# A combinatorial postsynaptic molecular mechanism converts patterns of nerve impulses into the behavioral repertoire

**DOI:** 10.1101/500447

**Authors:** Maksym V. Kopanitsa, Louie N. van de Lagemaat, Nurudeen O. Afinowi, Douglas J. Strathdee, Karen E. Strathdee, David G. Fricker, Eleanor J. Tuck, Kathryn A. Elsegood, Mike D. R. Croning, Noboru H. Komiyama, Seth G. N. Grant

## Abstract

How is the information encoded within patterns of nerve impulses converted into diverse behavioral responses? To address this question, we conducted the largest genetic study to date of the electrophysiological and behavioral properties of synapses. Postsynaptic responses to elementary patterns of activity in the hippocampal CA1 region were measured in 58 lines of mice carrying mutations in the principal classes of excitatory postsynaptic proteins. A combinatorial molecular mechanism was identified in which distinct subsets of proteins amplified or attenuated responses across timescales from milliseconds to an hour. The same mechanism controlled the diversity and magnitude of innate and learned behavioral responses. PSD95 supercomplex proteins were central components of this synaptic machinery. The capacity of vertebrate synapses to compute activity patterns increased with genome evolution and is impaired by disease-relevant mutations. We propose that this species-conserved molecular mechanism converts the temporally encoded information in nerve impulses into the repertoire of innate and learned behavior.

## Introduction

Information from the external world is converted into sequences of nerve impulses which are decoded or ‘read’ in the brain to generate representations, perceptions, memories, emotions, actions and all other behavioral responses. Individual impulses and pairs of impulses comprise the elementary syntax of longer sequences, such as bursts and rhythms. The timing of individual impulses or pairs of impulses separated even by milliseconds, has behavioral importance^1–6^. Identifying the molecular mechanisms underpinning the capacity of the brain to read patterns of neural activity and generate behavioral responses is a major goal in neuroscience.

The proteome of excitatory synapses is central to this question for several reasons. First, the output of excitatory synapses (postsynaptic depolarization) is adjusted depending on the incoming sequence of neural activity^7–10^ (Fig. 1A). Synapses can enhance or attenuate their postsynaptic responses during the pattern of activity itself as well as make long-lasting stable changes in synaptic strength^10^. This modulation is referred to as synaptic plasticity, and the many forms that have been described can be grouped according to the time scales when they are observed: milliseconds (e.g. paired-pulse facilitation, paired-pulse depression), seconds (e.g. augmentation, depletion), minutes (e.g. post-tetanic potentiation) and tens of minutes to hours (e.g. short-term potentiation, long-term potentiation [LTP], long-term depression)^11–16^. Second, the postsynaptic proteome of excitatory synapses in vertebrate species has a remarkable complexity, with ~1,000 highly conserved proteins from many classes, including neurotransmitter receptors, adhesion, scaffolding, signaling and structural proteins, and many of these proteins are known to play a role in synaptic plasticity^17–27^. However, it is unknown whether these proteins have specific roles in the detection and discrimination of patterns of activity, principally because there have been no side-by-side comparisons of their functions. Indeed, it remains unknown which of the many postsynaptic proteins are of greatest functional significance, and thus the core molecular mechanisms of the postsynaptic terminal of excitatory synapses are unclear. It has also been puzzling why the postsynaptic proteome is so complex and highly conserved when it has been thought that synaptic transmission and plasticity could be accomplished with only a small number of these proteins^28^. Finally, more than 130 monogenic and polygenic brain disorders, including schizophrenia, depression, autism and intellectual disability^20,29–34^ arise from mutations and variants in genes encoding postsynaptic proteins. The extent to which these mutations impact on the capacity of synapses to compute information encoded in patterns of activity is poorly understood.

**Figure 1.**
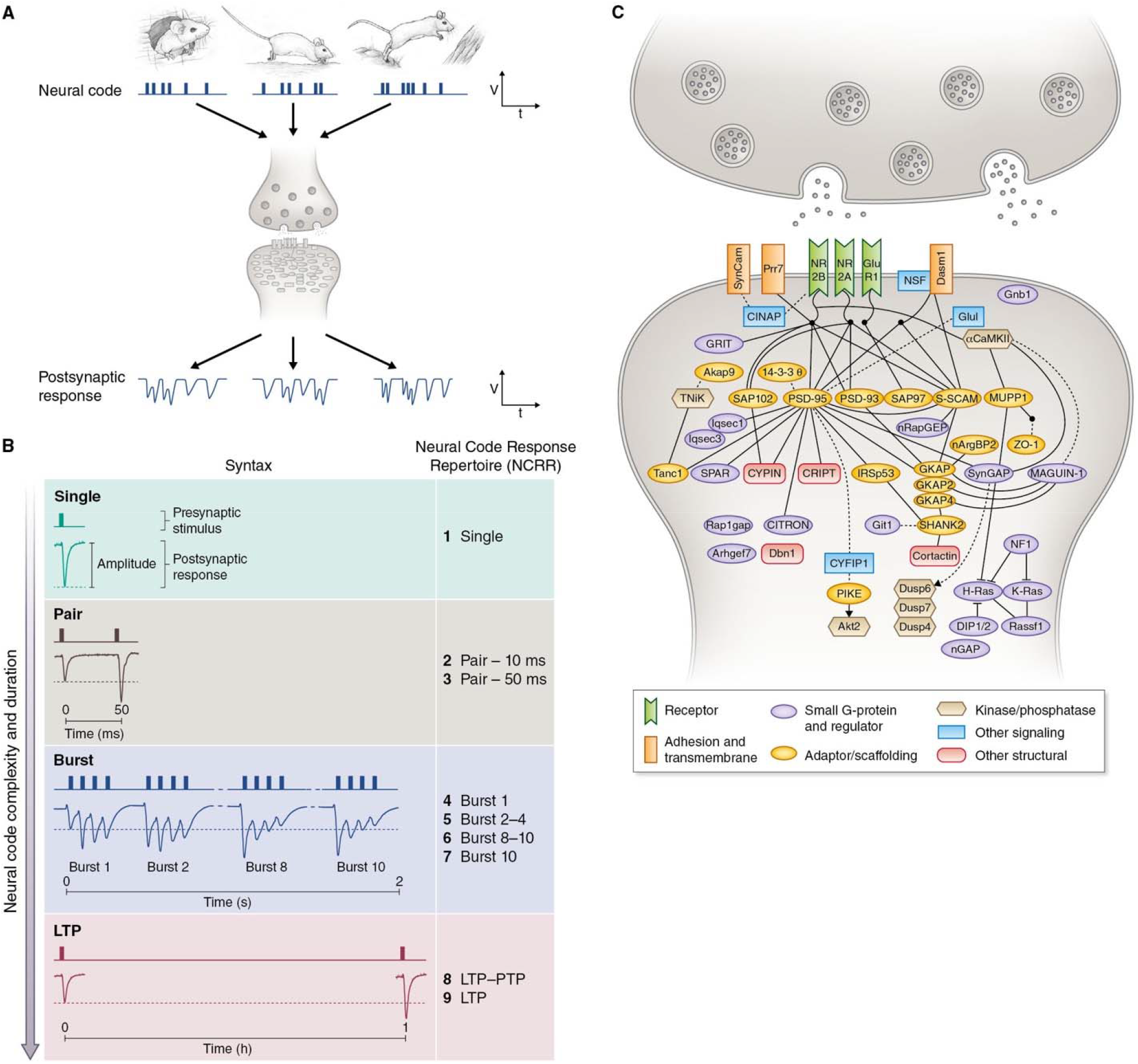
Genetic dissection of postsynaptic responses to neural codes. A. Different environmental stimuli are encoded into distinct patterns of nerve cell firing which arrive at excitatory synapses and generate postsynaptic responses of varying amplitude. V, voltage; t, time. B. The syntax of elementary sequences or ‘neural codes’ is represented by stimuli (Single, Pair, Burst, LTP) that together comprise a Neural Code Response Repertoire (NCRR) of 9 parameters spanning milliseconds to an hour. C. The postsynaptic proteins mutated in this study. Protein interactions are indicated (solid lines, binary interactions; dotted lines, other functional interactions). Key, major protein classes.

Here, we report a large-scale genetic dissection of the role of postsynaptic proteins in multiple forms of synaptic plasticity induced by elementary patterns of activity. In addition to a side-by-side comparative study of the electrophysiological phenotypes of mutations in many genes, we have measured innate and learned behavioral responses and hippocampal transcriptomes in the same mutant mouse lines^35^. These studies comprise the largest genetic analysis of the vertebrate synapse to date with phenotyping in molecular, electrophysiological and behavioral domains obtained using standardized quantitative approaches and analyzed with robust statistical methods. This data resource allowed us to uncover a novel molecular mechanism for synaptic computation that underpins the behavioral repertoire.

## Results

We studied fifty-eight lines of mice carrying engineered mutations in 51 genes of which several were well characterized and served as benchmarks for our experiments (Supplementary Fig.1, Supplementary Table 1). Most of the genes were previously uncharacterized and chosen because they represented major classes of postsynaptic proteins described in proteomic studies: glutamate receptors, surface adhesion proteins, adaptors and scaffolders, structural proteins, small G-proteins and regulators, signaling enzymes, kinases and phosphatases (Figs 1C, 2). Fifty-one lines harbored loss-of-function (LoF) mutations (homozygous null or heterozygous haploinsufficient), six lines carried knock-in mutations, and one line carried mutations in two genes encoding NMDA receptor subunits (Supplementary Data 1). These genes included paralogs from six protein families, phylogenetically conserved synapse proteins and orthologs of human disease genes (Fig. 2). All details of the mouse production and links to electrophysiological and behavioral data and other datasets are available on our website http://www.genes2cognition.org/publications/g2c/.

**Figure 2.**
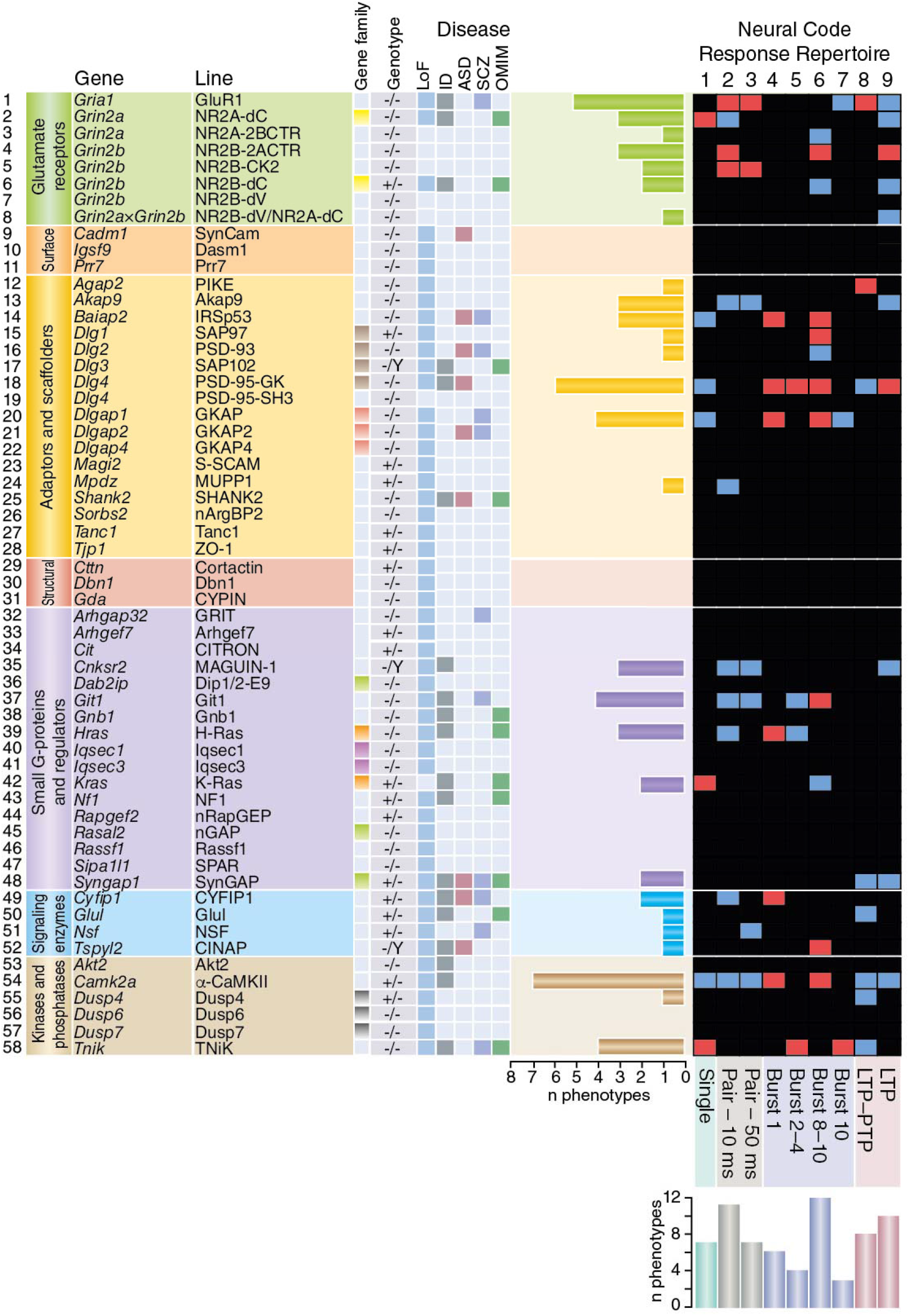
Summary of NCRR phenotypes. NCRR phenotypes (*P* < 0.05) in 58 lines of mutant mice are indicated by blue (attenuated electrophysiological response) or red (amplified electrophysiological response) squares; black indicates no significant phenotype compared with wild-type mice. The horizontal histogram shows the number of phenotypes for each gene/line and the vertical histogram (bottom) shows the number of phenotypes for each NCRR component. Gene family, boxes of the same color indicate paralogs; LoF, loss-of-function alleles; Orthologs of human disease genes: ID, intellectual disability; ASD, autism spectrum disorder; SCZ, schizophrenia; OMIM, neural diseases from Online Mendelian Inheritance of Man.

A highly standardized multi-electrode array (MEA)-based system was used to record field postsynaptic responses (fEPSPs) in the apical dendrites of CA1 pyramidal neurons following the stimulation of CA3 Schaffer collateral inputs in acute hippocampal slices from mutant and wild-type mice^36^ (Methods). Single, paired and multiple closely spaced stimuli (arranged in ten theta-bursts) evoked fEPSPs of differing amplitudes, consistent with previous literature (Fig. 1B). From each slice, we systematically acquired 18 measures of postsynaptic responses spanning the timescale of milliseconds to an hour. This was reduced to a set of nine minimally correlated measures, which we refer to as the Neural Code Response Repertoire (NCRR) (Fig. 1B, Methods). The nine NCRR parameters were subcategorized into four groups (Single, Pair, Burst, LTP) that modeled the increasing complexity and duration of activity patterns: *Single* was a measure of the absolute amplitude of the maximum fEPSP elicited by a single impulse; *Pair* defined the ratio between peak amplitudes of two submaximal responses separated by 10 or 50 ms (known as paired-pulse facilitation); *Burst* measures represented the submaximal fEPSP amplitudes during theta-bursts (1, 2–4, 8–10 and 10) from a 2-s-long train containing 10 bursts; *LTP* was the relative enhancement of the fEPSP at 60 min after a theta-burst stimulation (TBS), and LTP/PTP was the ratio of the LTP amplitude at the 60th min to the amplitude of the first fEPSP recorded after TBS, during post-tetanic potentiation (PTP) (Fig. 1C). A total of 2,836 slices were recorded from 58 lines of mice generating 51,048 postsynaptic measures from which 25,524 measures from the NCRR were used (9 measurements in 2,836 slices) (for numbers of slices and animals per line, see http://www.genes2cognition.org/publications/g2c/).

### Combinations of postsynaptic proteins decipher activity patterns

We first asked which postsynaptic proteins were required for each of the NCRR components (Fig. 2). Overall, from 51 lines of mice carrying single LoF mutations, the proportion showing phenotypes was 12.4% (68/459, *P* < 0.05). Remarkably, the response to a single impulse was sensitive to one combination of proteins, whereas another protein combination regulated pairs of impulses separated by 10 or 50 ms, and other combinations regulated the responses during bursts and LTP. Each of these combinations utilized all classes of postsynaptic signal transduction proteins, including glutamate receptors, adaptors and scaffolders, small G-proteins and regulators, signaling enzymes, kinases and phosphatases. Mutations in six structural and surface membrane molecules tested did not lead to significant electrophysiological phenotypes. Some proteins (e.g. PSD95, α-CamKII) were required for many of the NCRR components, indicating that they are involved in regulating responses over a wide temporal range, whereas other proteins had more restricted, specialized roles (see horizontal phenotype histograms, Fig. 2).

We were also struck by the observation that each combination comprised proteins that either attenuated or amplified the response magnitude (blue and red boxes, Fig. 2). The relative contribution made by each gene to amplification or attenuation is shown in plots ranking their effect size (Cohen’s d) value for each NCRR component (Fig. 3A). Counting the number of proteins that either amplified or attenuated each response revealed no detectable bias in this bidirectional modulation (χ^2^ test, *P* > 0.05) (Fig. 3B). Importantly, many mutations did not simply attenuate or amplify all affected NCRR components, but enhanced some and attenuated others, revealing that the repertoire of different temporal components is coordinately regulated. Together, this genetic dissection shows that the capacity of synapses to read and respond to patterns of activity is dependent on the coordinated function of combinations of postsynaptic proteins.

**Figure 3.**
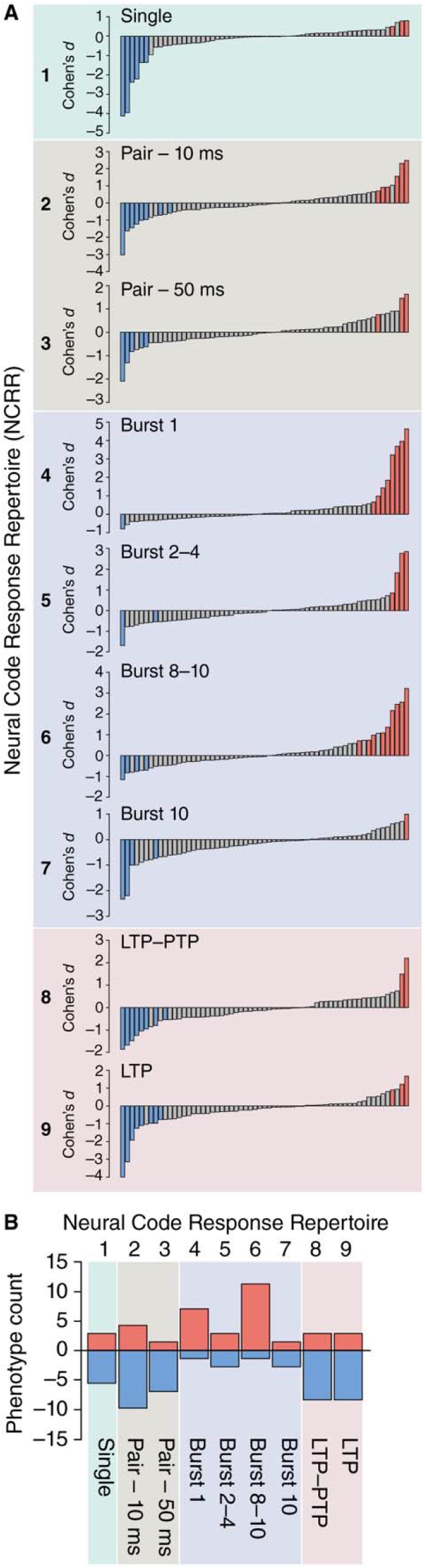
Bidirectional tuning and response optima. A. Ranking the direction and amplitude of LoF mutation Cohen’s *d* values of each NCRR variable (1-9 as indicated) for each LoF mutation. B. Number of LoF mutations amplifying (red) or attenuating (blue) each NCRR component.

### Major drivers of postsynaptic function

There have been no studies comparing the electrophysiological phenotypes of the many classes of postsynaptic proteins and therefore, the identity of the most important proteins for synaptic transmission and plasticity remains unknown. To address this, we derived an Overall NCRR score from the phenotype effect size (Cohen’s d) value of each NCRR parameter for each mutation. As shown in Figure 4, ranking the Overall NCRR scores for the 51 LoF lines showed that mutations in genes encoding the scaffold protein PSD95 and the enzyme α-CamKII had the strongest overall phenotype, followed by mutations in genes encoding GKAP, GRIA1, IRSp53, Git1, MAGUIN-1, NR2A, TNiK, NR2B and SynGAP. The relative contribution of each of these proteins to individual NCRR scores varied, indicating their differential contribution to the modulation of different temporal components of activity patterns (Fig. 4). This combination of postsynaptic proteins represents the prominent drivers of excitatory synapse function.

**Figure 4.**
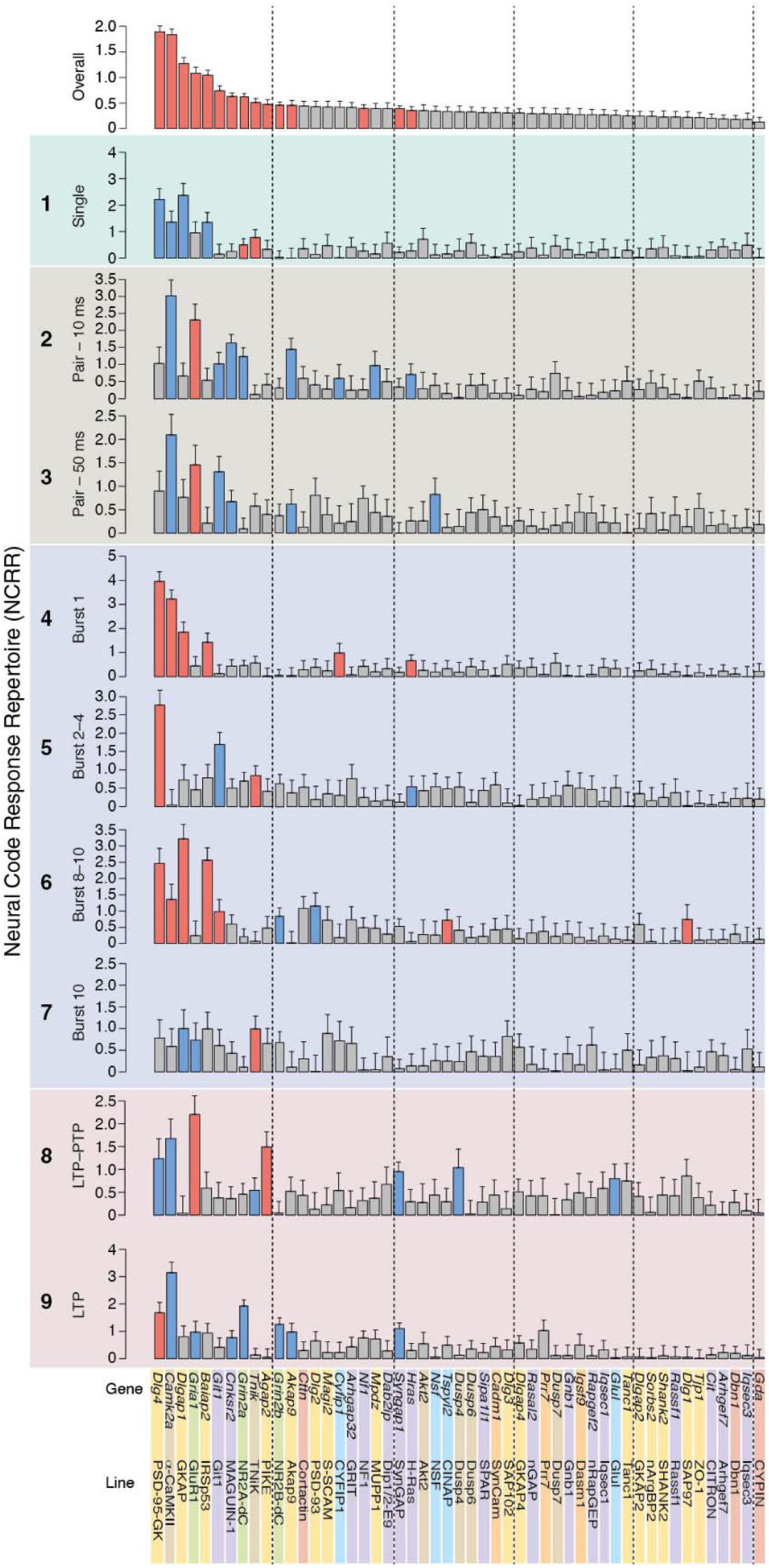
Overall and individual NCRR phenotype effect size values of individual mutants. The top panel shows the ranked Overall NCRR effect size (Cohen’s d) values for LoF mutations in 51 genes. Error bars represent standard error of the Cohen’s *d* estimate. Lower nine panels show mutant phenotypes for each NCRR component. Blue or red shading represents attenuated or amplified electrophysiological responses, respectively (*P* <0.05).

### Physiological and behavioral mechanisms are shared

There are three features in common between the results obtained from the genetic dissection of electrophysiological functions described above and the behavioral data presented in the companion manuscript^35^. First, combinations of postsynaptic proteins specify each innate or learned behavioral response, and each synaptic response to temporally encoded information. Second, each of these combinations is composed of proteins that amplified and attenuated the respective responses. Third, the major driver proteins in behavior (PSD95, TNiK, PSD93, GRIA1, Gnb1, IRSp53, GKAP4, Iqsec3)^35^ overlapped with those defined in the electrophysiological experiments.

To further examine the relationship between these sets of proteins, we tested the correlation between the Overall NCRR scores and the Overall Behavioral Repertoire (OBR)^35^ score using data from 45 LoF lines of mice that were tested in behavioral and electrophysiological experiments. This comparison showed a significant correlation (Pearson’s r = 0.401, *P* = 0.0067) (Fig. 5A) between behavior and physiology that was primarily dependent on seven proteins (PSD95, α-CamKII, GRIA1, TNiK, IRSp53, NR2A-dC, SynGAP) (Fig. 5A, B). These proteins are known constituents of PSD95 supercomplexes^24,29,30,37–39^, indicating that these signaling complexes are the major molecular machines in the postsynaptic terminal governing electrophysiological and behavioral responses.

**Figure 5.**
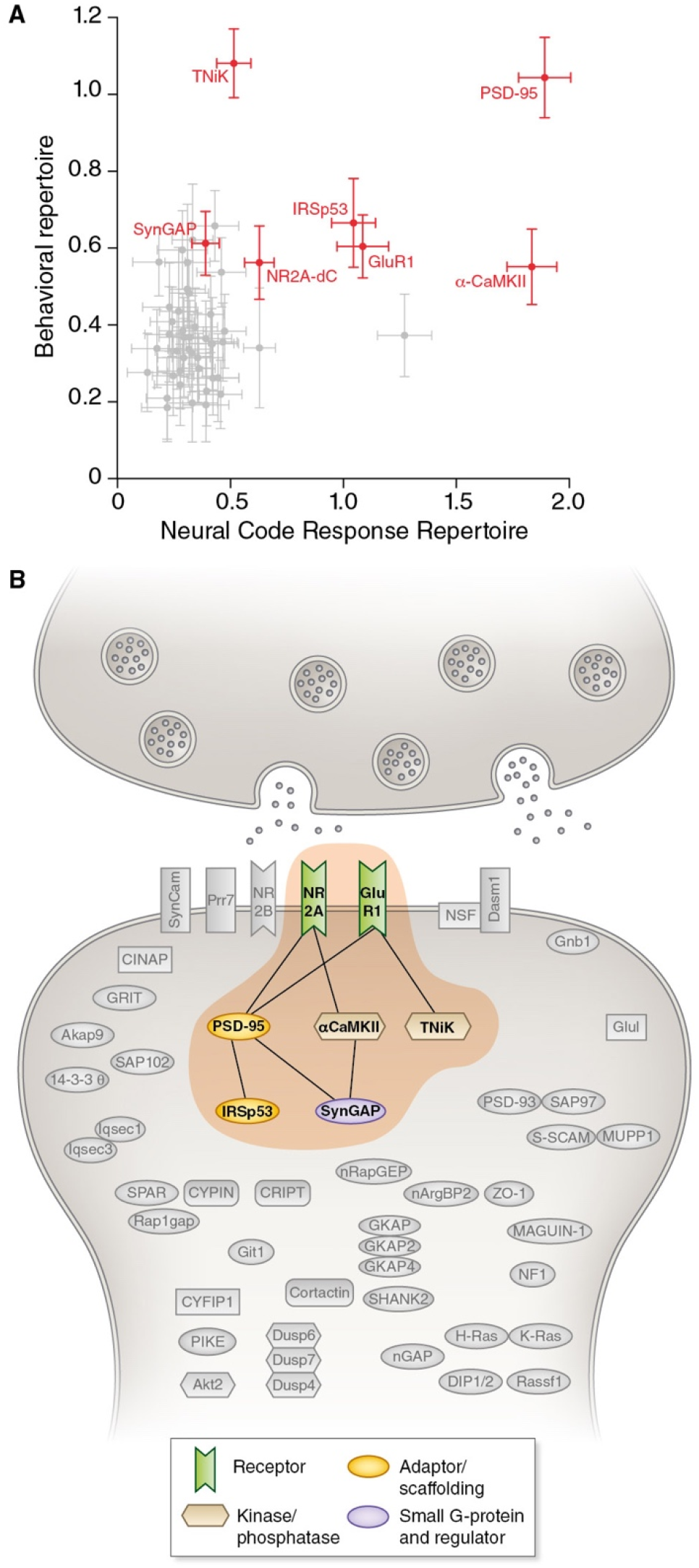
A core set of postsynaptic proteins controls the NCRR and behavioral repertoire. A. Correlation between Overall NCRR and Overall Behavioral Repertoire mutation effect size (Cohen’s d) scores. Large effect mutations in seven proteins (highlighted in red) were required for the correlation. Error bars represent standard error of the Cohen’s *d* estimate. B. The seven core proteins (brown zone) controlling NCRR and behavioral repertoire centered on PSD95 and satellite proteins (surrounding gray area in the postsynaptic terminal).

### Vertebrate proteome evolution generated response complexity

One of the most fundamental questions in neuroscience is how did humans and other vertebrates evolve their sophisticated cognitive functions from simpler organisms? Because our gene sets include paralogs in six families of postsynaptic proteins which arose from two whole-genome duplications ~550 million years ago^23,32,40–42^, our data afford a unique opportunity to ask if the evolution of these paralogs contributed to the capacity of vertebrate synapses to read information encoded in patterns of neural activity. As shown in Figure 6A, the different NCRR phenotypes in five out of six families of paralogs supports the conclusion that the genome duplications contributed greater molecular combinatorial complexity to vertebrate synapses, with a resultant increase in the capacity of synapses to read and respond to patterns of activity.

**Figure 6.**
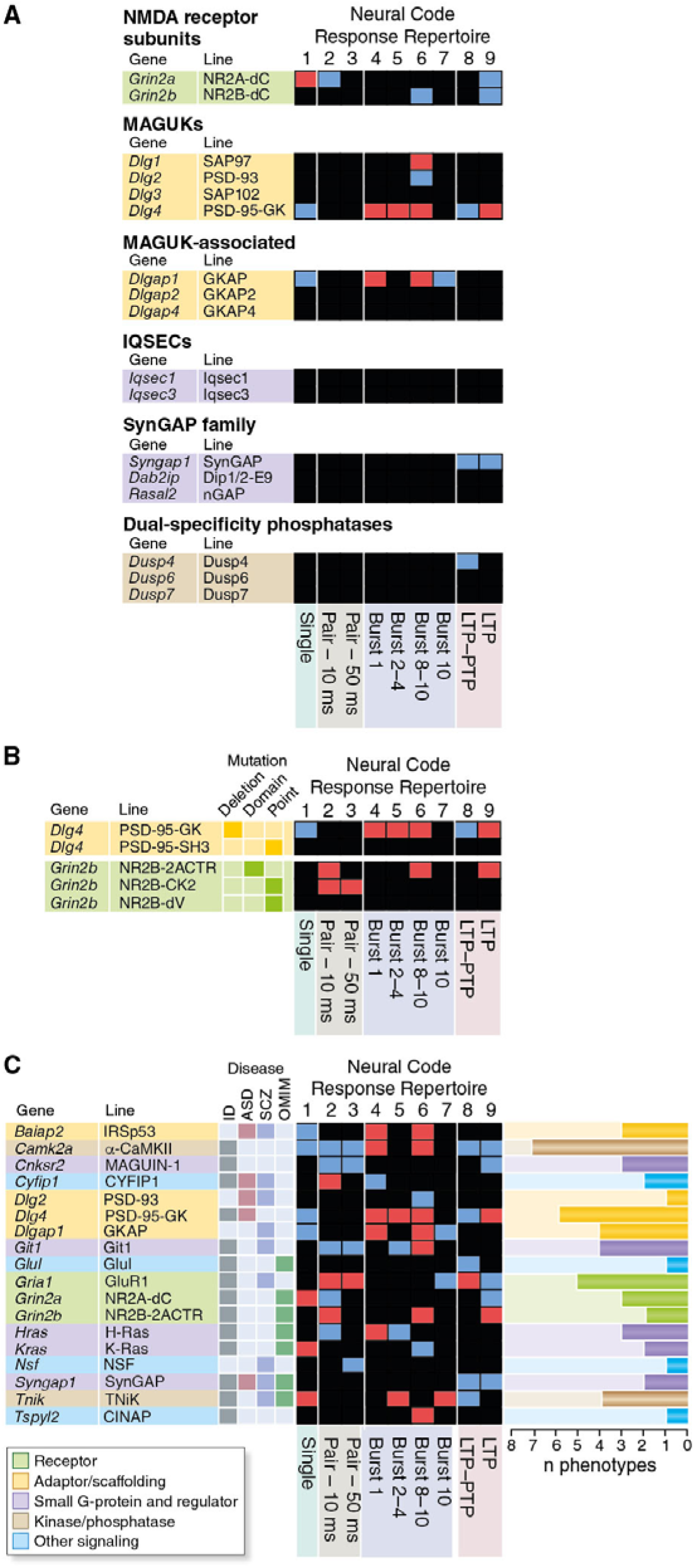
Vertebrate evolution and disease. A. NCRR phenotypes caused by mutations in vertebrate paralogs that belong to six protein classes. B. NCRR phenotypes caused by fine structural mutations. Gene deletion, domain deletion and point mutation are indicated. C. NCRR phenotypes caused by mutations in orthologs of human intellectual disability (ID), autism spectrum disorder (ASD) and schizophrenia (SCZ) genes; OMIM, neural diseases from Online Mendelian Inheritance of Man. Phenotypes (*P* < 0.05) are indicated by blue (attenuated electrophysiological response) or red (amplified electrophysiological response) squares; black indicates no significant phenotype compared with wild-type mice.

Because structural differences in paralogs arise by the accumulation of single nucleotide polymorphisms (SNPs) (and other mutations), it was of interest to ask how SNPs might reshape the NCRR. We examined lines of mice carrying engineered mutations in two gene families. A comparison of two different point mutations in the NR2B subunit of the NMDA receptor shows that the absence of the C-terminal valine (NR2B-dV mutation) had no significant phenotype, whereas the triple amino acid mutation disrupting the α-CamKII binding site (NR2B-CK2) had significant phenotypes in the Pair parameter (Fig. 6B). Mice carrying a complete LoF mutation in PSD95 (PSD-95-GK) showed major NCRR phenotypes, whereas a point mutation (PSD-95-SH3)^43^ introducing a single amino acid change in the binding site of the SH3 domain had no significant phenotype (Fig. 6B). Together, these results show that gene duplication and more subtle structural changes can shape the capacity of vertebrate synapses to process information in patterns of nerve cell activity.

### Diverse temporal response parameters are disrupted in human disorders

Mutations and genetic variation in postsynaptic proteins cause more than 130 monogenic and polygenic brain disorders, including schizophrenia, depression, autism and intellectual disability^20,29–34^. In addition, variation in IQ is associated with genetic variation in synaptic proteins and PSD95 supercomplexes^29,44^. Our datasets included mutations in 18 genes known to be mutated in intellectual disability (ID), autism spectrum disorder (ASD) and schizophrenia (SCZ) (Fig. 6C, Supplementary Table 2), providing a unique opportunity to ask if (i) these mutations result in impairments in the synaptic responses to patterns of activity and (ii) if there are any common or convergent phenotypes (e.g. LTP impairment) associated with diseases. As shown in Figure 6C, disease-relevant mutations disrupted all parameters. Furthermore, we observed both amplification and attenuation of the responses (Fig. 6C, red and blue boxes). These data suggest that ID, ASD, and SCZ are not caused by a reduction or increase in any single electrophysiological parameter but reflect a shift from the optimal response in multiple parameters.

## Discussion

To understand how the brain decodes information within the limitless number of patterns of neuronal activity that occur during behavior, we focused on a set of elementary patterns: single stimuli, pairs of stimuli and bursts. Remarkably, each of these elementary patterns required different combinations of postsynaptic proteins and each combination included proteins that amplified or attenuated each response. Patterns of increasing complexity (e.g. bursts) required multiple combinations of postsynaptic proteins at different points throughout the activity pattern, as judged from the sensitivity of burst elements to respective mutations. Thus, combinations of postsynaptic proteins enable synapses to read information encoded in patterns of action potentials and control the strength of synaptic transmission on short- and long-term timescales. Since the protein combinations regulate the elementary syntax that underlies more complex, naturally occurring patterns of activity, we suggest that the protein combinations identified in these experiments, together with many other combinations, are employed in physiological circumstances. The fact that the postsynaptic proteome is composed of >1,000 proteins^17–27^ suggests that there is a vast number of potential protein combinations available to read diverse patterns of activity.

Our integrated electrophysiological and behavioral data from the same mice show five important sets of convergent findings that point to a molecular and synaptic mechanism of behavior. First, each behavior and electrophysiological response utilized combinations of postsynaptic proteins. Second, the response magnitudes of all behavioral and electrophysiological parameters were found to be controlled by amplifying and attenuating proteins. Third, the main driver proteins affecting behavior and electrophysiology were the same and comprised those found in PSD95 supercomplexes. Fourth, paralogs diversified behavioral and electrophysiological responses. Fifth, comparison of SNPs with larger mutations showed similarly restricted and widespread phenotypes in electrophysiology and behavior.

These findings provide strong support for the following model. The postsynaptic proteome contains combinations of proteins that enable synapses to read temporally encoded information. These protein combinations set the optimal magnitude of diverse electrophysiological and behavioral responses. When the protein combinations are disturbed by pathogenic mutations or genetic variation, synapses ‘misread’ the information in patterns of activity and generate an inappropriate postsynaptic response that in turn drives neuronal activity leading to maladaptive behavioral responses. At the subsynaptic structural level, the combinations represent the physical organization of postsynaptic proteins into multiprotein complexes with PSD95 supercomplexes playing the major role. We suggest that these molecular machines are molecular building blocks for the repertoire of behaviors. We believe this molecular model complements Cajal’s ‘circuit model’ of behavior, in which neurons are the building blocks for the circuits mediating behaviors^45^, because excitatory synapses are molecularly diverse. Different combinations of postsynaptic proteins and multiprotein complexes are distributed into different synapses, neurons, circuits and brain regions^26,27,46,47^. Moreover, it has been shown that synapse molecular diversity by itself can generate spatio-temporal representations from incoming patterns of neural activity^47^. The rich anatomical complexity of synaptome maps reveals the spatial organization of protein complexes across brain regions controlling innate and learned behaviors^47^.

This combinatorial molecular mechanism also provides a basis for the evolution of vertebrate behavioral complexity and behavioral disorders arising from mutations. Our findings support the conclusion that vertebrate genome duplications and paralog diversification increased the capacity of synapses to compute more complex patterns of neuronal activity. The finding that different kinds of mutations (changes in single amino acids, proteins domains and gene deletions) affecting many protein classes all lead to changes in synaptic computation suggests that the many human disorders associated with genetic variation in the postsynaptic proteome have a direct impact on the capacity of synapses to compute temporally encoded information. All 18 disease-relevant mutations changed at least one synaptic response parameter, with most (14/18) changing multiple parameters. Our side-by-side comparison of electrophysiological phenotypes did not reveal any convergent phenotypes for ASD, schizophrenia or intellectual disability genes. These disorders could be considered as arising from a shift in the optimal response to patterns of activity and an overall defect in decoding temporal information.

In our studies, we have focused on elementary patterns of neural activity, and there are many electrophysiological stimulation paradigms, built from these elementary patterns, that induce forms of plasticity. We therefore expect that the combinatorial molecular principles would apply to these paradigms. We do not think that the combinatorial principles will be restricted to postsynaptic proteins because paralogs of presynaptic proteins, such as Munc13^48^, are known to play differential roles in synaptic plasticity. Finally, the data resources generated from this study can be used for many applications in computational neuroscience, behavior and physiology, medical genetics and genomics. We suggest that integrated analyses of large-scale phenotype data spanning molecules, synapse physiology and behavior can be applied to other sets of proteins in the nervous system.

## Methods

### Preparation of hippocampal slices

Experimental procedures have been previously described^36^. In brief, 3- to 8-month-old mutant and litter-matched, or in rare cases age-matched, wild-type mice were sacrificed by cervical dislocation and the brain immediately immersed in ice-cold “cutting” solution (110 mM sucrose, 60 mM NaCl, 28 mM NaHCO3, 1.25 mM NaH2PO4, 3 mM KCl, 7 mM MgSO_4_, 0.5 mM CaCl_2_, 5 mM glucose, 0.6 mM sodium ascorbate, 0.015 mM phenol red, pH 7.2) gassed with a mixture of 95% O_2_ and 5% CO_2_. Whole brain slices were cut at 350 μm thickness by a Vibroslice MA752 (Campden Instruments, Loughborough, UK) in such a way so that the blade would cut through hemispheres at an angle of 20–30° to their horizontal planes. “Cutting” solution in the temperature-controlled Peltier bath was maintained at 0–3 °C and constantly saturated with a mixture of 95% O_2_ and 5% CO_2_. Up to eight slices containing medial segments of the hippocampus with overlaying cortical areas were trimmed of the remaining tissue, placed into a well of a slice chamber (Fine Science Tools, Foster City, CA, USA) and kept interfaced between moist air and subfused fresh artificial cerebrospinal fluid (ACSF) that contained 124 mM NaCl, 25 mM NaHCO_3_, 1 mM NaH_2_PO_4_, 4.4 mM KCl, 1.2 mM MgSO_4_, 2 mM CaCl_2_, 10 mM glucose, and 0.015 mM phenol red (pH 7.3-7.4). Temperature in the chamber was slowly increased to 30 °C for the rest of the incubation time. Slices rested in these conditions for at least 2 h before experiments commenced.

### Electrophysiological recording

Field excitatory postsynaptic potentials (fEPSPs) were recorded by the MEA60 electrophysiological suite (Multi Channel Systems, Reutlingen, Germany). Four set-ups consisting of a MEA1060-BC pre-amplifier and a filter amplifier (gain 550×) were run simultaneously by a data acquisition card operated by the MC_Rack software. Raw electrode data were digitized at 10 kHz and stored on a PC hard disk for subsequent analysis. To record fEPSPs, a hippocampal slice was placed into the well of 5×13 3D MEA biochip (Qwane Biosciences, Lausanne, Switzerland). The slice was guided to a desired position with a fine paint brush and gently fixed over MEA electrodes by a silver ring with attached nylon mesh lowered vertically by a one-dimensional U-1C micromanipulator (Narishige, Tokyo, Japan). MEA biochips were fitted into the preamplifier case and fresh ACSF was delivered to the MEA well through a temperature-controlled perfusion cannula that warmed perfusing media to 32 °C. All experiments were carried out in submerged slices. Monopolar stimulation of the Schaffer collateral/commissural fibers through array electrodes was performed by STG2008 and STG4008 stimulus generators (Multi Channel Systems). Biphasic (positive/negative, 100 μs/phase) voltage pulses were used. Amplitude, duration and frequency of stimulation were controlled by the MC_Stimulus II software. As the data acquisition card could process data only from 128 channels, we recorded from only 30 electrodes per chip (out of a possible 60), which enabled to use four independent set-ups.

We performed all LTP experiments using two-pathway stimulation of Schaffer collateral/commissural fibers ^49,50^. Our previous experiments showed that largest LTP was recorded in proximal parts of the CA1 stratum radiatum^36^. We therefore picked a single principal recording electrode in the middle of the proximal part of CA1 and assigned two electrodes for stimulation of the control and test pathways on the subicular side and on the CA3 side of stratum radiatum, respectively. The distance from the recording electrode to the test stimulation electrode was 400–560 μm and to the control stimulation electrode was 316–560 μm. To evoke orthodromic fEPSPs, stimulation electrodes were activated at a frequency of 0.02 Hz. Following at least 15–30 min of equilibration period inside the MEA well, input/output relationships were obtained and baseline stimulation strength was set to evoke a response that corresponded to 40% of the maximal attainable fEPSP at the principal recording electrode. To investigate shortterm plasticity, we used paired-pulse stimulation of the test pathway with an inter-pulse interval of 10 and 50 ms. Paired-pulse ratio was calculated by dividing the amplitude of the second fEPSP in a pair by the amplitude of the first one. Average data from five paired-pulse stimulations were used for each slice. LTP was induced after 30 min period of stable baseline responses by applying theta-burst stimulation (TBS) train consisting of 10 bursts given at 5 Hz with 4 pulses given at 100 Hz per burst. Stimulus strength was not altered during TBS.

## Data analysis

Amplitude of the negative part of fEPSPs induced by the maximal attainable stimulus (4.2 V) in the test pathway was used as a measure of synaptic strength. Data were loaded into an R environment from .mcd files using the NeuroShare library freely available from Multi Channel Systems. LTP plots were scaled to the average of the first five baseline points. LTP magnitude was assessed by averaging normalized fEPSPs in the test pathway 60–65 min after TBS episode. To account for possible drift of baseline conditions, amplitude values in the test pathway were normalized by respective amplitudes in the control pathway prior to statistical comparison of LTP values. In addition to the peak fEPSP and LTP, 16 other variables were measured, including paired-pulse facilitation at 50 ms and 10 ms interpulse intervals, defacilitation observed in the third fEPSP of a 100 Hz train of four fEPSPs, two measures of plasticity and ten measures of fEPSP amplitude during TBS, and a measure of fEPSP amplitude decay after TBS. Thus, these 18 variables used measures of fEPSP amplitude to address synaptic transmission and plasticity in single fEPSPs, pairs of fEPSPs, and trains of fEPSPs. Retaining only one of the ten highly redundant measures of burst amplitude reduced the number of variables analyzed to nine. Definitions are provided in the table below and online (for example, see: http://www.genes2cognition.org/publications/g2c/electrophysiology/m00000082/)

| Main paper name | Short name | Description |
| --- | --- | --- |
| Single | Max fEPSP amplitude | Maximum field excitatory postsynaptic potential (fEPSP) amplitude. |
| Pair – 50 ms | PPF at 50ms, ampl ratio | Paired-pulse facilitation (PPF), pulses separated by 50 ms, amplitude ratio. |
| Pair – 10 ms | Burst 1, PPF at 10ms, ampl ratio | Paired-pulse facilitation (PPF), pulses separated by 10 ms, |
|  |  | amplitude ratio, observed during first two pulses of the first 100 Hz burst during theta-burst stimulation. |
| Burst 1 | Burst 1, EPSP3 depr, ampl ratio | Depression observed in third fEPSP relative to the second fEPSP of the first 100 Hz burst, amplitude ratio. |
| Burst 2-4 | Burst 2-4 ave amplitude vs Burst 1 | Facilitation observed in average amplitude of bursts 2-4, relative to average amplitude of burst 1. |
| Burst 8-10 | Burst 8-10 ave amplitude vs Burst 2-4 | Depression observed in average amplitude of bursts 8-10, relative to average amplitude of bursts 2-4. |
| Burst 10 | Burst 10, ave amplitude | Average amplitude of four fEPSPs in tenth burst. |
| LTP-PTP | LTP vs PTP, amplitude ratio | Reduction in potentiation from first EPSP after theta-burst stimulation to 1 hour later, fEPSP amplitude ratio. |
| LTP | LTP based on amplitude | Long-term potentiation, ratio of amplitudes of fEPSPs in test pathway and control pathway. |

Because several slices from the same animal were routinely analyzed, the significance of changes in these measures in slices from mutant mice compared with the values in slices from wild-type mice was assessed using nested ANOVA with genotype as “fixed” factor and mouse as “random” factor. In addition, the Satterthwaite’s correction for unequal sample sizes was used (McDonald, 2014). Phenotype effect sizes were calculated using measurements in individual slices. Standard error of effect size was calculated by 1,000 random samples of wild-type mouse identifiers, matched for the background strain. In each sampling step, the same numbers of mice were used as in the original experiments (usually five mice representing mutants and five representing wild types). Simulated effect sizes were approximately normally distributed about zero. Half the central 68th percentile range, which approximates the standard deviation of this distribution, was used as the standard error of the effect size.

## Acknowledgements

We thank R. Sprengel and P. Seeburg for Gria1/GluR1, Grin2a/NR2A-dC and Grin2b/NR2B-dC mice, D. Bredt for Dlg2/PSD93 mice, and Louise van der Weyden for Cadm1/SynCam and Rassf1 mice. We are grateful to T.J. O’Dell and P. Charlesworth for electrophysiology advice. Computational support was provided by J. Menendez Montes and J. Piatkowski; technical support was provided by V.J. Robinson, S. Syed Salim, G. Berry, E. Stebbings, E. Trenchard and R. Uren. Administrative support was provided by J.V. Turner. We thank M. van Rossum, T.J. O’Dell and E. Fransén for comments on the manuscript and D. Maizels for artwork. Funding: Wellcome Trust, Wellcome Trust Sanger Institute and European Union Seventh Framework Programme under grant agreement numbers HEALTH-F2-2009-241498 (“EUROSPIN” project), HEALTH-F2-2009-242167 (“SynSys project”) and HEALTH-F2-2009-241995 (“GENCODYS project”).

## Data availability

The raw and analyzed data are archived and freely available at http://www.genes2cognition.org/publications/g2c.

## Author contributions

Electrophysiology experiments, MVK, NOA; Gene targeting/mouse genetics NHK, DJS, KES, DGF, EJT, KAE; Project tracking and colony management, KAE, MVK, MDRC; Statistical analysis, LNL, MVK; Database and website, MDRC; Direction and writing, SGNG.

## Supplementary Materials

Fig S1

Table S1-2

Supplementary Data File 1

## References

1 Borst, A. & Theunissen, F. E. Information theory and neural coding. Nat Neurosci 2, 947–957, doi:10.1038/14731 (1999).

2 Maass, W. & Bishop, C. M. Pulsed neural networks. (MIT Press, 1999).

3 Pfeifer, R. Connectionism in perspective. (North-Holland; Distributors for the U.S. and Canada, Elsevier Science Pub. Co., 1989).

4 Rolls, E. T. & Tovee, M. J. Processing speed in the cerebral cortex and the neurophysiology of visual masking. Proc Biol Sci 257, 9–15, doi:10.1098/rspb.1994.0087 (1994).

5 Stanley, G. B. Reading and writing the neural code. Nat Neurosci 16, 259–263, doi:10.1038/nn.3330 (2013).

6 Thorpe, S. J. & Imbert, M. in Connectionism in perspective (eds R. Pfeifer, Z. Schreter, F. Fogelman-Soulie, & L. Steels) (North-Holland; Distributors for the U.S. and Canada, Elsevier Science Pub. Co., 1989).

7 Dittman, J. S., Kreitzer, A. C. & Regehr, W. G. Interplay between facilitation, depression, and residual calcium at three presynaptic terminals. J Neurosci 20, 1374–1385 (2000).

8 Markram, H., Gupta, A., Uziel, A., Wang, Y. & Tsodyks, M. Information processing with frequency-dependent synaptic connections. Neurobiol Learn Mem 70, 101–112, doi:10.1006/nlme.1998.3841 (1998).

9 Segundo, J. P., Moore, G. P., Stensaas, L. J. & Bullock, T. H. Sensitivity of Neurones in Aplysia to Temporal Pattern of Arriving Impulses. J Exp Biol 40, 643–667 (1963).

10 Abbott, L. F. & Regehr, W. G. Synaptic computation. Nature 431, 796–803, doi:10.1038/nature03010 (2004).

11 Bliss, T. V. & Lomo, T. Long-lasting potentiation of synaptic transmission in the dentate area of the anaesthetized rabbit following stimulation of the perforant path. J Physiol 232, 331–356 (1973).

12 Carlisle, H. J., Fink, A. E., Grant, S. G. & O’Dell, T. J. Opposing effects of PSD-93 and PSD-95 on long-term potentiation and spike timing-dependent plasticity. J Physiol 586, 5885–5900, doi:10.1113/jphysiol.2008.163469 (2008).

13 Citri, A. & Malenka, R. C. Synaptic plasticity: multiple forms, functions, and mechanisms. Neuropsychopharmacology 33, 18–41, doi:10.1038/sj.npp.1301559 (2008).

14 Ito, M. Long-term depression. Annual review of neuroscience 12, 85–102, doi:10.1146/annurev.ne.12.030189.000505 (1989).

15 Thomson, A. M. Facilitation, augmentation and potentiation at central synapses. Trends Neurosci 23, 305–312 (2000).

16 Zucker, R. S. Short-term synaptic plasticity. Annual review of neuroscience 12, 13–31, doi:10.1146/annurev.ne.12.030189.000305 (1989).

17 Bayes, A. et al. Comparative study of human and mouse postsynaptic proteomes finds high compositional conservation and abundance differences for key synaptic proteins. PLoS One 7, e46683, doi:10.1371/journal.pone.0046683 (2012).

18 Bayes, A. et al. Human post-mortem synapse proteome integrity screening for proteomic studies of postsynaptic complexes. Mol Brain 7, 88, doi:10.1186/s13041-014-0088-4 (2014).

19 Bayes, A. et al. Zebrafish synapse proteome complexity, evolution and ultrastructure. Nat Commun 8, 14613, doi:10.1038/ncomms14613 (2017).

20 Bayes, A. et al. Characterization of the proteome, diseases and evolution of the human postsynaptic density. Nat Neurosci 14, 19–21, doi:10.1038/nn.2719 (2011).

21 Collins, M. O. et al. Molecular characterization and comparison of the components and multiprotein complexes in the postsynaptic proteome. J Neurochem 97 Suppl 1, 16–23, doi:10.1111/j.1471-4159.2005.03507.x (2006).

22 Distler, U. et al. In-depth protein profiling of the postsynaptic density from mouse hippocampus using data-independent acquisition proteomics. Proteomics 14, 2607–2613, doi:10.1002/pmic.201300520 (2014).

23 Emes, R. D. et al. Evolutionary expansion and anatomical specialization of synapse proteome complexity. Nat Neurosci 11, 799–806, doi:10.1038/nn.2135 (2008).

24 Husi, H., Ward, M. A., Choudhary, J. S., Blackstock, W. P. & Grant, S. G. Proteomic analysis of NMDA receptor-adhesion protein signaling complexes. Nat Neurosci 3, 661–669 (2000).

25 Peng, J. et al. Semiquantitative proteomic analysis of rat forebrain postsynaptic density fractions by mass spectrometry. J Biol Chem 279, 21003–21011, doi:10.1074/jbc.M400103200 (2004).

26 Roy, M. et al. Regional Diversity in the Postsynaptic Proteome of the Mouse Brain. Proteomes 6, doi:10.3390/proteomes6030031 (2018).

27 Roy, M. et al. Proteomic analysis of postsynaptic proteins in regions of the human neocortex. Nat Neurosci 21, 130–138, doi:10.1038/s41593-017-0025-9 (2018).

28 Nicoll, R. A. A Brief History of Long-Term Potentiation. Neuron 93, 281–290, doi:10.1016/j.neuron.2016.12.015 (2017).

29 Fernandez, E. et al. Arc Requires PSD95 for Assembly into Postsynaptic Complexes Involved with Neural Dysfunction and Intelligence. Cell Rep 21, 679–691, doi:10.1016/j.celrep.2017.09.045 (2017).

30 Fernandez, E. et al. Targeted tandem affinity purification of PSD-95 recovers core postsynaptic complexes and schizophrenia susceptibility proteins. Mol Syst Biol 5, 269, doi:10.1038/msb.2009.27 (2009).

31 Fromer, M. et al. De novo mutations in schizophrenia implicate synaptic networks. Nature 506, 179–184, doi:10.1038/nature12929 (2014).

32 Nithianantharajah, J. et al. Synaptic scaffold evolution generated components of vertebrate cognitive complexity. Nat Neurosci 16, 16–24, doi:10.1038/nn.3276 (2013).

33 Purcell, S. M. et al. A polygenic burden of rare disruptive mutations in schizophrenia. Nature 506, 185–190, doi:10.1038/nature12975 (2014).

34 Howard, D. M. et al. Genome-wide association study of depression phenotypes in UK Biobank identifies variants in excitatory synaptic pathways. Nat Commun 9, 1470, doi:10.1038/s41467-018-03819-3 (2018).

35 Komiyama, N. H. et al. Synaptic combinatorial molecular mechanisms generate repertoires of innate and learned behavior. (submitted).

36 Kopanitsa, M. V., Afinowi, N. O. & Grant, S. G. Recording long-term potentiation of synaptic transmission by three-dimensional multi-electrode arrays. BMC Neurosci 7, 61, doi:10.1186/1471-2202-7-61 (2006).

37 Coba, M. P. et al. TNiK is required for postsynaptic and nuclear signaling pathways and cognitive function. J Neurosci 32, 13987–13999, doi:10.1523/JNEUROSCI.2433-12.2012 (2012).

38 Frank, R. A. et al. NMDA receptors are selectively partitioned into complexes and supercomplexes during synapse maturation. Nat Commun 7, 11264, doi:10.1038/ncomms11264 (2016).

39 Frank, R. A. W., Zhu, F., Komiyama, N. H. & Grant, S. G. N. Hierarchical organisation and genetically separable subfamilies of PSD95 postsynaptic supercomplexes. J Neurochem, doi:10.1111/jnc.14056 (2017).

40 Emes, R. D. & Grant, S. G. The human postsynaptic density shares conserved elements with proteomes of unicellular eukaryotes and prokaryotes. Front Neurosci 5, 44, doi:10.3389/fnins.2011.00044 (2011).

41 Emes, R. D. & Grant, S. G. Evolution of synapse complexity and diversity. Annual review of neuroscience 35, 111–131, doi:10.1146/annurev-neuro-062111-150433 (2012).

42 Ryan, T. J. et al. Evolution of GluN2A/B cytoplasmic domains diversified vertebrate synaptic plasticity and behavior. Nat Neurosci 16, 25–32, doi:10.1038/nn.3277 (2013).

43 Arbuckle, M. I. et al. The SH3 domain of postsynaptic density 95 mediates inflammatory pain through phosphatidylinositol-3-kinase recruitment. EMBO Rep 11, 473–478, doi:10.1038/embor.2010.63 (2010).

44 Hill, W. D. et al. Human cognitive ability is influenced by genetic variation in components of postsynaptic signalling complexes assembled by NMDA receptors and MAGUK proteins. Transl Psychiatry 4, e341, doi:10.1038/tp.2013.114 (2014).

45 Shepherd, G. M. Foundations of the neuron doctrine. 25th anniversary edition. edn, (Oxford University Press, 2016).

46 Emes, R. D. et al. Evolutionary expansion and anatomical specialization of synapse proteome complexity. Nat Neurosci 11, 799–806, doi:10.1038/nn.2135 (2008).

47 Zhu, F. et al. Architecture of the Mouse Brain Synaptome. Neuron 99, 781–799 e710, doi:10.1016/j.neuron.2018.07.007 (2018).

48 Rosenmund, C. et al. Differential control of vesicle priming and short-term plasticity by Munc13 isoforms. Neuron 33, 411–424 (2002).

49 Andersen, P., Sundberg, S. H., Sveen, O. & Wigstrom, H. Specific long-lasting potentiation of synaptic transmission in hippocampal slices. Nature 266, 736–737 (1977).

50 Schwartzkroin, P. A. & Wester, K. Long-lasting facilitation of a synaptic potential following tetanization in the in vitro hippocampal slice. Brain Res 89, 107–119 (1975).

