## Supplementary Data 1 for "A combinatorial postsynaptic molecular mechanism converts patterns of nerve impulses into the behavioral repertoire"

**Contents:**

For each mouse line (alphabetical order) the details of mouse generation, genotyping and breeding is provided. Further information is available on the G2C website: http://www.genes2cognition.org/publications/g2c

### *Ywhaq* (14-3-3 *theta*)

#### 1 Mouse generation

##### 1.1 Mutation

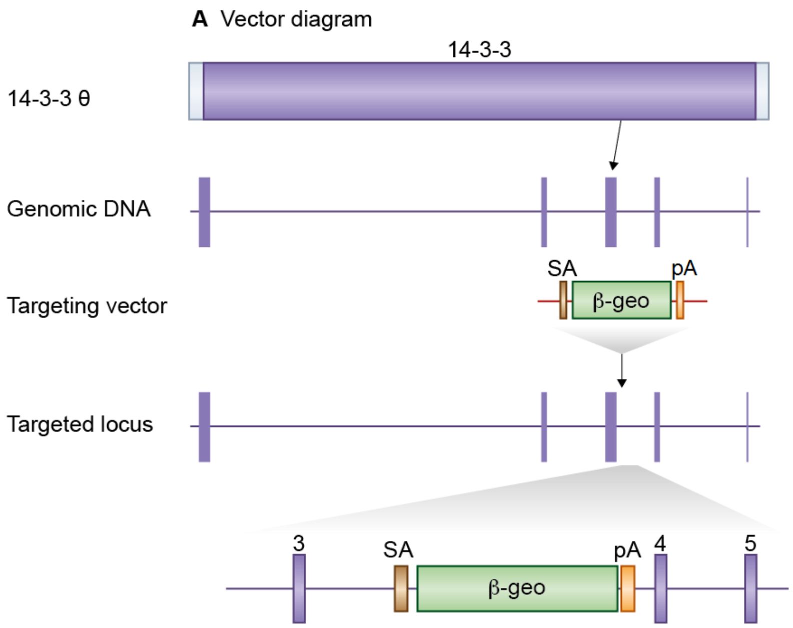


**Figure 1:** *Details of the mutation*

A mouse embryonic stem (ES) cell line (AL0734, strain 129/Ola) with an insertional mutation in *Ywhaq* was obtained from Sanger Institute Gene Trap Resource (SIGTR - sanger.ac.uk/PostGenomics/genetrap/). The insertional mutation in AL0734 by the gene-trapping vector, pGT0lxr, that was designed to create an in-frame fusion between the 5' exons of the trapped gene and a reporter, beta-geo (a fusion of beta-galactosidase and neomycin phosphotransferase II) occurred in intron 3-4. Thus, the gene-trapped locus is predicted to yield a fusion transcript containing exons 1-3 of *Ywhaq* and ß-geo. The ES cells were injected into C57BL/6 blastocysts to create chimeric mice, which were bred with 129S5 mice to generate heterozygous *Ywhaq* mutant mice. Those F1 heterozygous mice had been backcrossed with 129S5 mice for 1-2 times before being used for intercrossing to produce homozygous mutants. Location of *Ywhaq* gene trap. *Ywhaq* is a 5 exon gene which encodes a protein with a 14-3-3 domain (top). The *Ywhaq* gene trap is located in intron 3-4.

#### 1.2 Genotyping

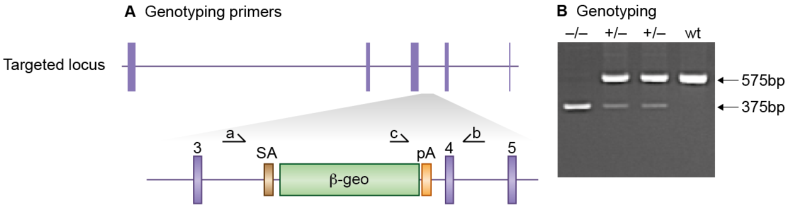


**Figure 2:** *Genotyping details*

Genomic DNA was isolated from ES cells or mouse tissues by Wizard SV 96 Genomic DNA purification system (Promega Cat A2371). Genotyping PCR consisted of a 575bp product amplified from the wild-type (wt) allele using a forward primer A (5'- CAATGCAACTAATCCAGAGAG -3') upstream of the cassette and a reverse primer B (5'- CAGATTACTACCCTACATGTG -3') in the wt sequence deleted by targeted mutation. A 375bp product was amplified from the targeted allele using primer B with forward primer C (5'- GATCTGCACTGTCCCGGATG -3'), within the ß-geo cassette. After enzymatic amplification for 35 cycles (45 seconds at 94 degC, 45 seconds at 55 degC, and 1 minute at 72 degC), the PCR products were size-fractionated on a 2% agarose gel in 1x Tris borate-EDTA buffer. Primers used for genotyping (a,b, c). PCR genotyping of gene trap 14-3-3 $\theta$ mice using a common reverse primer, b, and forward primers a and c to amplify the wt and mutant alleles respectively.

#### 1.3 Breeding

Birth of 14-3-3 $\theta^{-/-}$ mice followed Mendelian ratios with 19% of offspring being homozygous knockouts. Genotypes of 3-week-old pups from 14-3-3 $\theta^{+/-}$ intercrosses identified 33 wt, 51 14-3-3 $\theta^{+/-}$ and 19 14-3-3 $\theta^{-/-}$ progeny (Chi-squared p= 0.148). Male and female 14-3-3 $\theta$-/- mice developed normally to adulthood, exhibited normal body size and no gross abnormalities. 14-3-3 $\theta$ mice were maintained by backcrossing onto the 129S5/SvEvBrd background; heterozygous males and females were fertile and used to set up intercrosses to generate homozygous and wildtype mice to study.

### *Akap9* (Akap9)

#### 1 Mouse generation

##### 1.1 Mutation

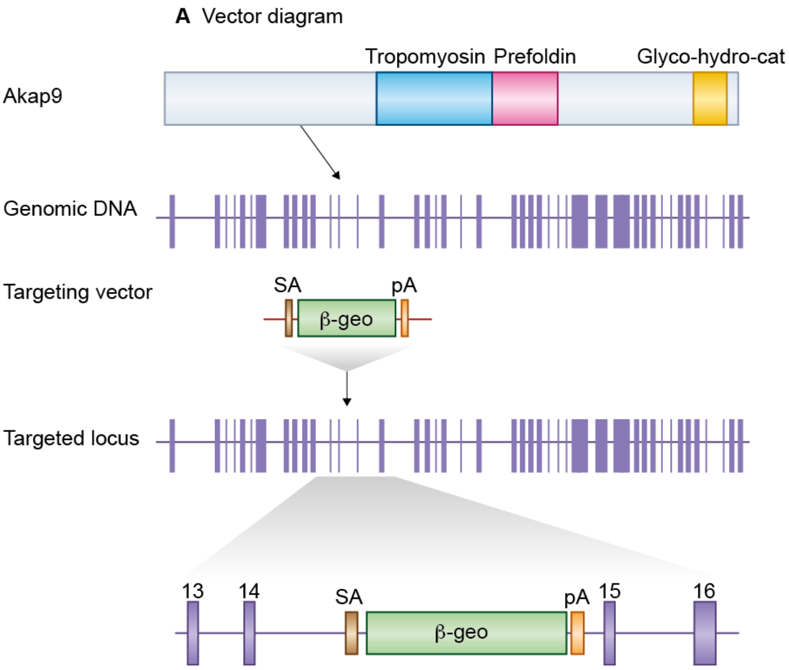


**Figure 1:** *Details of the mutation*

A mouse embryonic stem (ES) cell line (XP0050, strain 129/Ola) with an insertional mutation in *Akap9* was obtained from Sanger Institute Gene Trap Resource (SIGTR - sanger.ac.uk/PostGenomics/genetrap/). The insertional mutation in XP0050 by the gene-trapping vector, pGT0lxr, that was designed to create an in-frame fusion between the 5' exons of the trapped gene and a reporter, beta-geo (a fusion of beta-galactosidase and neomycin phosphotransferase II) occurred in intron 14-15. Thus, the gene-trapped locus is predicted to yield a fusion transcript containing exons 1-14 of *Akap9* and ß-geo. The ES cells were injected into C57BL/6 blastocysts to create chimeric mice, which were bred with 129S5 mice to generate heterozygous *Akap9* mutant mice. Location of *Akap9* gene trap. *Akap9* is a 48 exon gene encoding a protein which contains Tropomysin, Prefoldin and Glyco-hydro-cat domains (top). The *Akap9* gene trap is located in intron 14-15.

#### 1.2 Genotyping

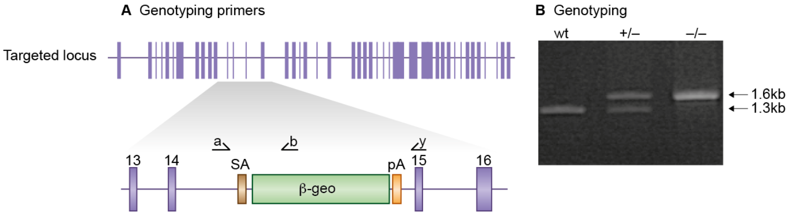


**Figure 2:** *Genotyping details*

Genomic DNA was isolated from ES cells or mouse tissues by Wizard SV 96 Genomic DNA purification system (Promega Cat A2371). Genotyping PCR consisted of a 1.3kb product amplified from the wild-type (wt) allele using a forward primer A (5'- CTTAGCCTGAGATGACTCAG -3') upstream of the cassette with reverse primer Y. A 1.6kb product was amplified from the targeted allele using primer A with reverse primer B (5'- GGTTACGTTGGTGTAGATGG -3'), within the beta-geo cassette. After enzymatic amplification for 35 cycles (30 seconds at 94 degC, 1 minute at 55 degC, and 3 minutes at 72 degC), the PCR products were size-fractionated on a 1% agarose gel in 1x Tris borate-EDTA buffer. Primers used for genotyping (a,b, y). PCR genotyping of gene trap *Akap9* mice using a common forward primer, a, and reverse primers y and b to amplify the wt and mutant alleles respectively.

#### 1.3 Breeding

Birth of *Akap9*$^{-/-}$ mice followed Mendelian ratios with 25% of offspring being homozygous knockouts. Genotypes of 3-week-old pups from *Akap9*$^{+/-}$ intercrosses identified 67 wt, 120 *Akap9*$^{+/-}$ and 56 *Akap9*$^{-/-}$progeny (Chi-squared p= 0.597). Male and female *Akap9*$^{-/-}$ mice developed normally to adulthood, exhibited normal body size and no gross abnormalities *Akap9* mice were maintained by backcrossing onto the 129S5/SvEvBrd background; heterozygous males and females were fertile and used to set up intercrosses to generate homozygous and wildtype mice to study.

### *Akt2* (Akt2)

### 1 Mouse generation

#### 1.1 Mutation

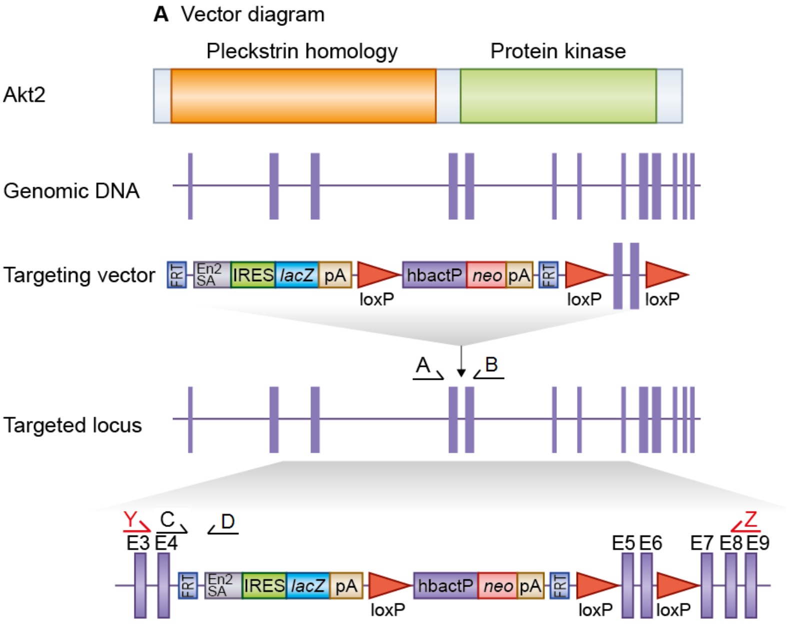


**Figure 1:** *Details of the mutation*

Akt2 mice with trapped allele *Akt2*$^{tm1Wcs}$ (colony MAAE) were supplied by the Mouse Genome Project (Sanger Institute). Generation of *Akt2* targeted mice. *Akt2* is a 14 exon gene encoding a protein which contains Pleckstrin homology and protein kinase domains (top). A selection cassette was inserted into intron 4-5 whilst exons 5,6 were floxed creating a frame shift between exons 4 and 7.

#### 1.2 Genotyping

Genotyping PCR consisted of a 583bp product amplified from the wild-type (wt) allele using a forward primer A (CGGGCAGTATTCAAAGCCAG) and reverse primer B (TAATCCCAACACTCGGGAGG). A 316bp product was amplified from the targeted allele using forward primer C (AGACTGGGGCAGGCTAAGTG) with reverse primer D (TCGTGGTATCGTTATGCGCC). After enzymatic amplification for 35 cycles (45 seconds at 94 degC, 45 seconds at 55 degC, and 1 minute at 72 degC), the PCR products were size-fractionated on a 2% agarose gel in 1x Tris borate-EDTA buffer (image not shown).

#### 1.3 Breeding

Birth of *Akt2* mice followed Mendelian ratios with 33% of offspring being homozygous knockouts. Genotypes of 3-week-old pups from *Akt2*$^{-/-}$ intercrosses identified 7 wt, 30 *Akt2*$^{+/-}$ and 18 *Akt2*$^{-/-}$ progeny (Chi-squared p= 0.088). Male and female *Akt2*$^{-/-}$ mice developed normally to adulthood, exhibited normal body size and no gross abnormalities. *Akt2* mice were maintained by backcrossing onto the C57BL/6J background; heterozygous males and females were fertile and used to set up intercrosses to generate homozygous and wildtype mice to study.

### *Arhgef7* (Arhgef7)

#### 1 Mouse generation

Generation of *Arhgef7* mutant mice described in Omelchenko et al (2014)

##### 1.2 Genotyping

*Arhgef7* genotyping described in Omelchenko *et al* (2014)

##### 1.3 Breeding

*Arhgef7*-/- mice were known to be lethal so *Arhgef7*+/- mice were produced for phenotypic analysis. Male and female *Arhgef7*+/- mice developed normally to adulthood, were fertile, exhibited normal body size and no gross abnormalities. Birth of *Arhgef7*+/- mice followed Mendelian ratios with 49% of offspring being homozygous knockouts. Genotypes of 3-week old-pups from *Arhgef7*+/- back-crosses identified 98 wildtype and 94 *Arhgef7*+/- progeny (chi-squared p= 0.77). Backcrosses onto the C57BL/6J background were used to maintain the colony.

### Camk2a (*alpha*-CaMKII)

#### 1 Mouse generation

##### 1.1 Mutation

Generation of *Camk2a* mutant mice described in PMID:1378648, Silva et al (1992)

##### 1.2 Genotyping

*Camk2a* genotyping described in PMID:1378648, Silva et al (1992)

##### 1.3 Breeding

Heterozygous *Camk2a+/-* mice and wildtype littermates were generated f om backcrosses for analysis, with litters fostered to C57BL6c-/c- parents since female heterozygous *Camk2a*+/- mice were found to be poor mothers. Genotypes of 3-week-old pups from *Camk2a*+/- backcrosses identified 63 wt and 52 *Camk2a*+/- progeny (Chi-squared p= 0.789). Male and female *Camk2a*+/- mice developed normally to adulthood, exhibited normal body size and no gross abnormalities. *Camk2a*+/- mice were maintained by backcrossing onto the C57BL/6J background.

### *Tspyl2* (CINAP)

#### 1 Mouse generation

##### 1.1 Mutation

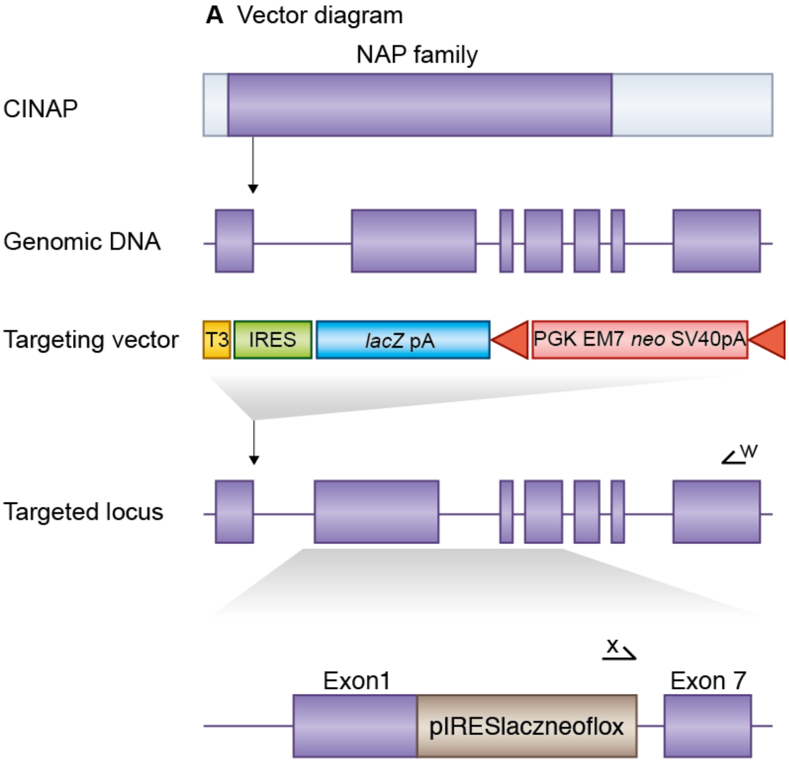


**Figure 1:** *Details of the mutation*

E14TG2a mouse embryonic stem (ES) cells were targeted with a vector containing 6.9kb and 2.9kb of flanking genomic DNA. This replaced 3.5kb of *Tspyl2* genomic DNA (X148773141 to X148776721; Ensemble Build 56) with IRES-lacZ-neo cassette. Correctly targeted ES cells were identified by long range PCR using Expand Long Template PCR system (Roche Cat 11681842001). The PCR contained primer X (5'-GAGCTATTCCAGAAGTAGTGAG-3') and primer W (5'- CAGTCCCTGAGAAGCACTTG -3') that correspond to sequence in the IRES-lacZ-neo cassette and sequence outside the 2.9kb flanking region respectively. The correctly targeted ES cells were injected into C57BL/6 blastocysts to create chimeric mice, which were bred with 129S5 mice to generate heterozygous *Tspyl2* mutant mice. Location of *Tspyl2* gene trap. *Tspyl2* is a 7 exon gene encoding the CINAP protein which contains a NAP family domain (top). We replaced *Tspyl2* exons 2-6 with a selection cassette in targeted mice and created a frameshift between exons 1 and 7. Primers used for targeted clone identification (w,x) are shown.

#### 1.2 Genotyping

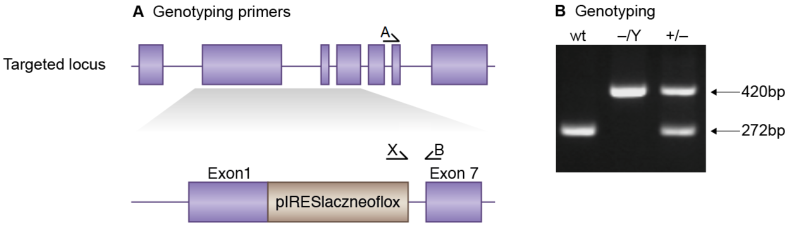


**Figure 2:** *Genotyping details*

Genomic DNA was isolated from ES cells by Wizard SV 96 Genomic DNA purification system (Promega Cat A2371). Genotyping PCR consisted of a 272bp product amplified from the wild-type (wt) allele using a forward primer A (CAGACCTCTTGACCACAAAAC) in the wt sequence deleted by targeted mutation and a reverse primer B (CTGCACACTTTCACCCTTAG) downstream of the cassette. A 420bp product was amplified from the targeted allele using primer B with forward primer X, within the selection cassette. After enzymatic amplification for 35 cycles (45 seconds at 94 degC, 45 seconds at 55 degC, and 1 minute at 72 degC), the PCR products were size-fractionated on a 2% agarose gel in 1x Tris borate-EDTA buffer. Primers used for genotyping (A,B, X). PCR genotyping of targeted *Tspyl2* mice using a common reverse primer, B, and forward primers A and X to amplify the wt and mutant alleles respectively.

#### 1.3 Breeding

*Tspyl2* resides on the X chromosome: birth of *Tspyl2*$^{-/Y}$ mice followed Mendelian ratios with 21% of offspring being hemizygous knockouts but no female *Tspyl2*$^{-/-}$ mice were born. Genotypes of 3-week-old pups from backcrosses identified 106 wt, 56 female *Tspyl2*$^{-/+}$ and 45 *Tspyl2*$^{-/Y}$ male progeny (Chi-squared p= 0.077). Male *Tspyl2*$^{-/Y}$ developed normally to adulthood, exhibited normal body size and no gross abnormalities. Backcrosses onto the 129S5/SvEvBrd background were used to maintain the colony and to generate hemizygous and wildtype mice to study.

### *Cit* (CITRON)

#### 1 Mouse generation

CITRON mice were supplied by the Mouse Genome Project (Sanger Institute).

##### 1.1 Mutation

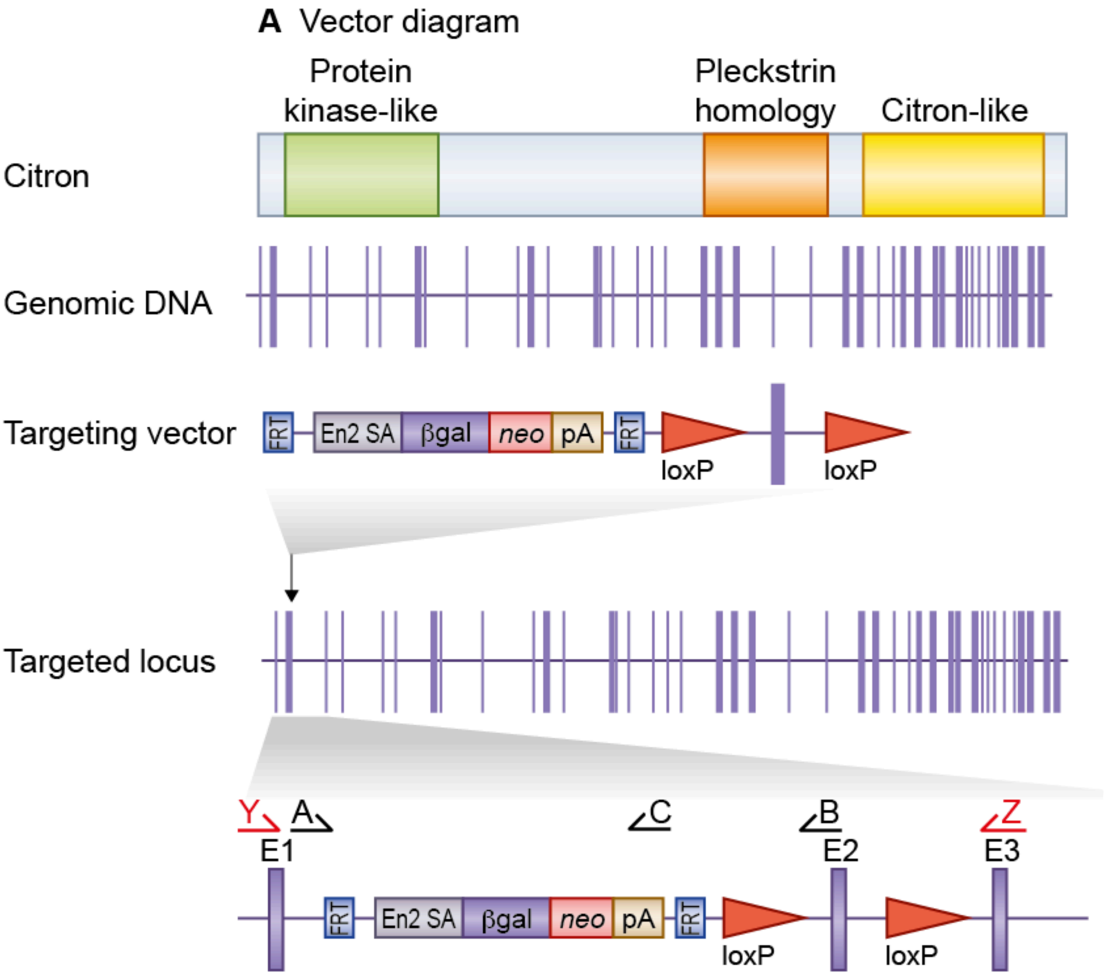


**Figure 1:** *Details of the mutation*

Generation of *Cit* targeted mice **A**. *Cit* is a 47 exon gene encoding the CITRON protein which contains Protein kinase-like, Pleckstrin homology and Citron-like domains (top). A selection cassette was inserted into intron 1-2 whilst exon 2 was floxed creating a frame shift between exons 1 and 3.

#### 1.2 Genotyping

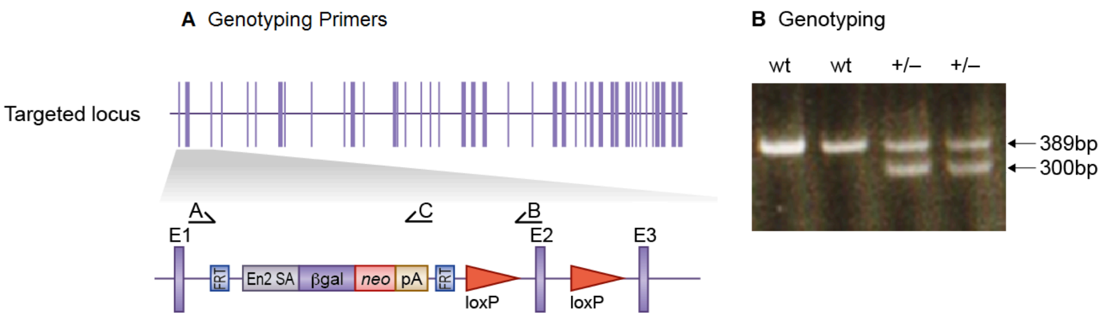


**Figure 2:** *Genotyping details*

Genotyping PCR consisted of a 389bp product amplified from the wild-type (wt) allele using a forward primer A (TTGGCTTCCCAACTCAGGTC) and reverse primer B (ACATTCCTTCCCTGTGTGGC). A 300bp product was amplified from the targeted allele using forward primer A with reverse primer C (TCGTGGTATCGTTATGCGCC). After enzymatic amplification for 35 cycles (45 seconds at 94 °C, 45 seconds at 55 °C, and 1 minute at 72 °C), the PCR products were size-fractionated on a 2% agarose gel in 1x Tris borate-EDTA buffer.

#### 1.3 Breeding

*Cit* $^{-/-}$ mice were known to be sub-viable so *Cit* $^{+/-}$ mice were produced for phenotypic analysis. Male and female *Cit* $^{+/-}$ developed normally to adulthood, were fertile, exhibited normal body size and no gross abnormalities. Genotypes of 3-week-old pups from *Cit* $^{+/-}$ back-crosses identified 103 wildtype and 114 *Cit* $^{+/-}$ progeny. Backcrosses onto the C57BL/6J background were used to maintain the colony.

### *Cttn* (Cortactin)

#### 1 Mouse generation

##### 1.1 Mutation

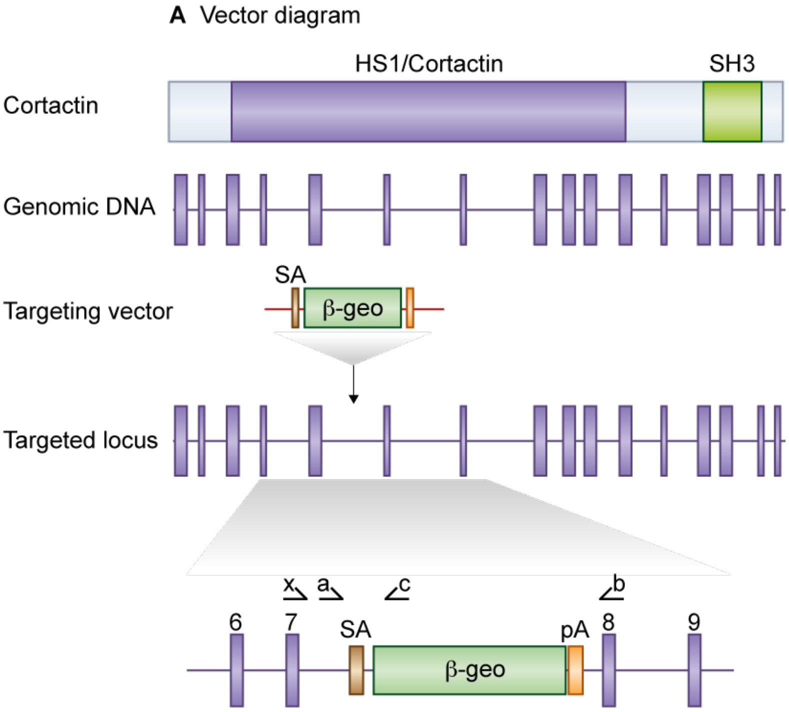


**Figure 1:** *Details of the mutation*

A mouse embryonic stem (ES) cell line (RRS284, strain 129/Ola) with an insertional mutation in *Cttn* was obtained from BayGenomics (baygenomics.ucsf.edu/). The insertional mutation in RRS284, by the gene-trapping vector, pGT1lxf, that was designed to create an in-frame fusion between the 5' exons of the trapped gene and a reporter, ß-geo (a fusion of ß-galactosidase and neomycin phosphotransferase II), occurred in intron 7-8. Thus, the gene-trapped locus is predicted to yield a fusion transcript containing exons 1-7 of *Cttn* and ß-geo. The ES cells were injected into C57BL/6 blastocysts to create chimeric mice, which were bred with 129S5 mice to generate heterozygous *Cttn* mutant mice. Location of *Cttn* gene trap mice. *Cttn* is an 18 exon gene, encoding a protein containing a HS1/Cortactin domain and SH3 Domain (top). The *Cttn* gene trap is located in intron 7-8. Primers used for genotyping (a,b, c) and RT-PCR (x,b) are shown.

#### 1.2 Genotyping

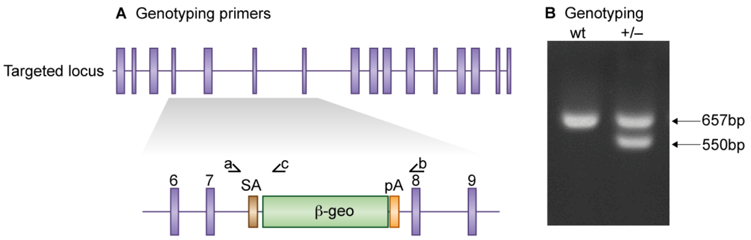


**Figure 2:** *Genotyping details*

Genomic DNA was isolated from ES cells or mouse tissues by Wizard SV 96 Genomic DNA purification system (Promega Cat A2371). Genotyping PCR consisted of a 657bp product amplified from the wild-type (wt) allele using a forward primer a (5'- GAACTCTGTCTGCACAGTTTG -3') upstream of the cassette with primer b. A 550bp product was amplified from the targeted allele using primer a with reverse primer c (5'-CAGTCCTCTTCACATCCATG -3') within the ß-geo cassette. After enzymatic amplification for 35 cycles (45 seconds at 94 degC, 45 seconds at 55 degC, and 1 minute at 72 degC), the PCR products were size-fractionated on a 2% agarose gel in 1x Tris borate-EDTA buffer.

#### 1.3 Breeding

No *Cttn*$^{-/-}$ mice were produced from *Cttn*$^{+/-}$ intercrosses. Male and female *Cttn*$^{+/-}$ mice de-veloped normally to adulthood, were fertile, exhibited normal body size and no gross abnormalities. Genotypes of 3-week-old pups from *Cttn*$^{+/-}$ intercrosses identified 22 wt, and 59 *Cttn*$^{+/-}$ progeny (Chi-squared p= $<$0.001). Backcrosses onto the 129S5/SvEvBrd background were used to maintain the colony and to gen-erate heterozygous and wildtype mice to study.

### *Cript* (CRIPT)

#### 1 Mouse generation

##### 1.1 Mutation

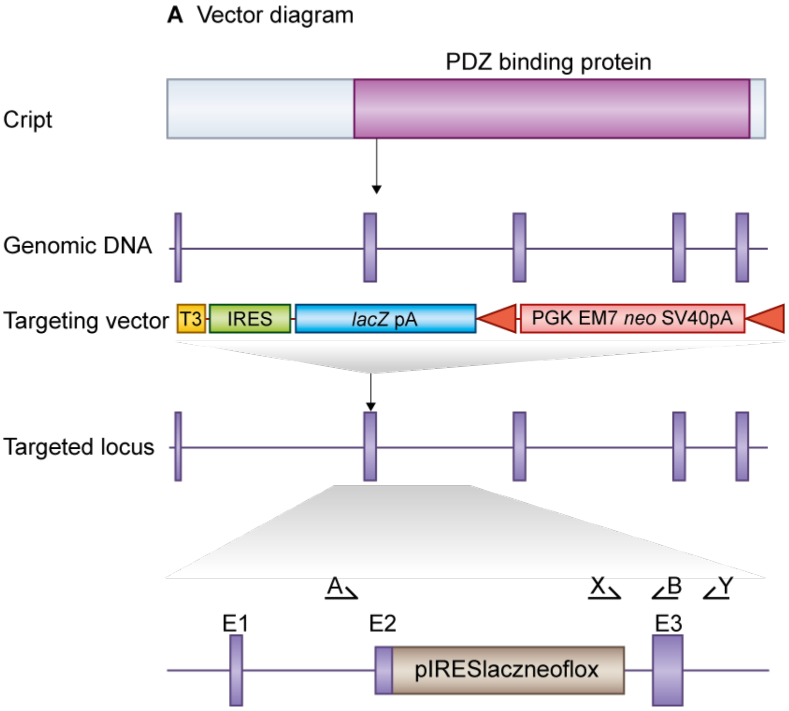


**Figure 1:** *Details of the mutation*

E14TG2a mouse embryonic stem (ES) cells were targeted with a vector containing 6kb and 2.5kb of flanking genomic DNA. This replaced 2kb of *Cript* genomic DNA (X87427053 to X87429095; Ensemble Build 55) with an IRES-lacZ-neo cassette. Genomic DNA was isolated from ES cells by Wizard SV 96 Genomic DNA purification system (Promega Cat A2371). Correctly targeted ES cells were identified by long range PCR using Expand Long Template PCR system (Roche Cat 11681842001). The PCR contained primer X (5'-GAGCTATTCCAGAAGTAGTGAG-3') and primer Y (5'- CCAAACTGAACTCAGATCCTC -3') that correspond to sequence in the IRES-lacZ-neo cassette and sequence outside the 2.5kb flanking region respectively. The correctly targeted ES cells were injected into C57BL/6 blastocysts to create chimeric mice, which were bred with 129S5 mice to generate heterozygous *Cript* mutant mice. Generation of *Cript* targeted mice. *Cript* is 5 exon gene which encodes a protein that contains a PDZ protein binding domain (top). We replaced most of *Cript* exon 2 with a selection cassette in targeted mice and created a frameshift between exons 2 and 3. Primers used for genotyping(a,b, x) and targeted clone identification (x,y) are shown.

#### 1.2 Genotyping

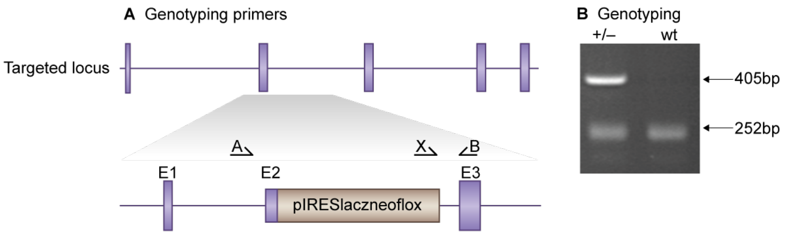


**Figure 2:** *Genotyping details*

Genotyping PCR consisted of a 252bp product amplified from the wild-type (wt) allele using a forward primer A (GAACTACTGGGACAATTGATG) in the wt sequence deleted by targeted mutation and a reverse primer B (CTCACAGTGTCAAGCTTCAG) downstream of the cassette. A 405bp product was amplified from the targeted allele using primer B with forward primer X, within the selection cassette. After enzymatic amplification for 35 cycles (45 seconds at 94 degC, 45 seconds at 55 degC, and 1 minute at 72 degC), the PCR products were size-fractionated on a 2% agarose gel in 1x Tris borate-EDTA buffer.

#### 1.3 Breeding

No *Cript*$^{-/-}$ mice were produced from *Cript*$^{+/-}$ intercrosses. Male and female *Cript*$^{+/-}$ mice developed normally to adulthood, were fertile, exhibited normal body size and no gross abnormalities. Genotypes of 3-week-old pups from *Cript*$^{+/-}$ intercrosses identified 36 wt, and 95 *Cript*$^{+/-}$ progeny (Chi-squared p= <0.001). Backcrosses onto the 129S5/SvEvBrd background were used to maintain the colony and to generate heterozygous and wildtype mice to study.

### *Cyfip1* (CYFIP1)

#### 1 Mouse generation

##### 1.1 Mutation

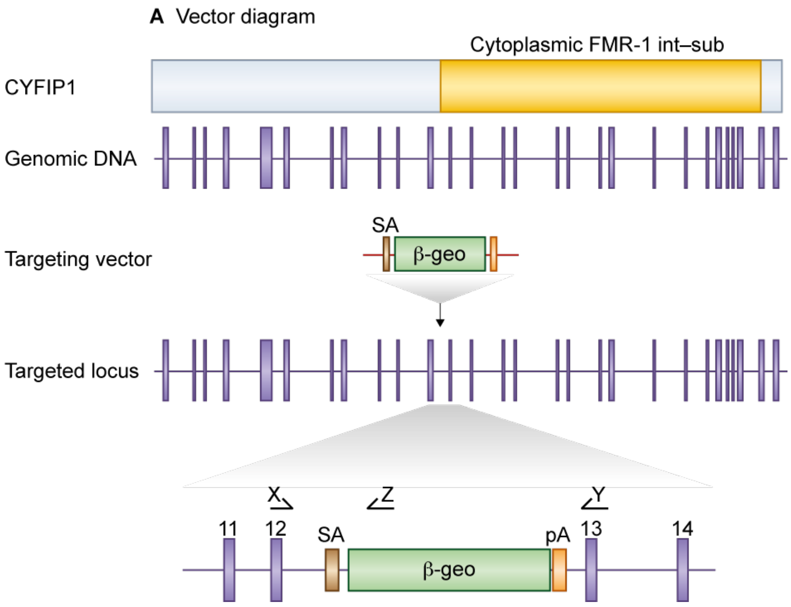


**Figure 1:** *Details of the mutation*

A mouse embryonic stem (ES) cell line (XG769, strain 129/Ola) with an insertional mutation in *Cyfip1* was obtained from BayGenomics (baygenomics.ucsf.edu/). The insertional mutation in XG769, by the gene-trapping vector, pGT1lxf, that was designed to create an in-frame fusion between the 5' exons of the trapped gene and a reporter, ß-geo (a fusion of ß-galactosidase and neomycin phosphotransferase II), occurred in intron 12-13. Thus, the gene-trapped locus is predicted to yield a fusion transcript containing exons 1-12 of *Cyfip1* and beta-geo. The ES cells were injected into C57BL/6 blastocysts to create chimeric mice, which were bred with 129S5 mice to generate heterozygous *Cyfip1* mutant mice.

#### 1.2 Genotyping

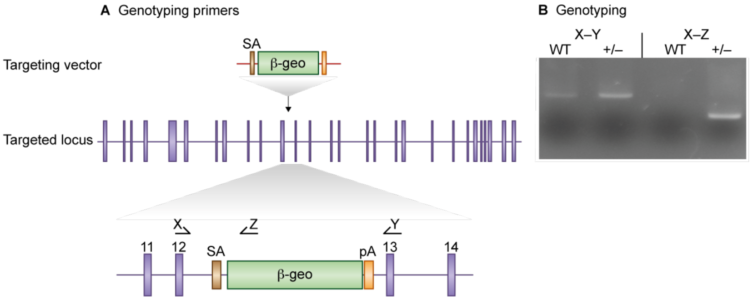


**Figure 2:** *Genotyping details*

Genomic DNA was isolated from ES cells or mouse tissues by Wizard SV 96 Genomic DNA purification system (Promega Cat A2371). Genotyping PCR consisted of a 1.2kb product amplified from the wild-type (wt) allele using a forward primer X and a reverse primer Y as above. A 1kb product was amplified from the targeted allele using primer X with reverse primer Z (5'- GGTTACGTTGGTGTAGATGG -3'), within the ß-geo cassette. After enzymatic amplification for 35 cycles (45 seconds at 94 degC, 45 seconds at 55 degC, and 1 minute at 72 degC), the PCR products were size-fractionated on a 2% agarose gel in 1x Tris borate-EDTA buffer. Primers used for genotyping (X,Y, Z). PCR genotyping of gene trap *Cyfip1* mice using a common forward primer, X, and reverse primers Y and Z to amplify the wt and mutant alleles respectively.

#### 1.3 Breeding

No *Cyfip1*$^{-/-}$ mice were produced from *Cyfip1*$^{+/-}$ intercrosses. Male and female *Cyfip1*$^{+/-}$ mice devel-oped normally to adulthood, were fertile, exhibited normal body size and no gross abnormalities. Genotypes of 3-week-old pups from *Cyfip1*$^{+/-}$ intercrosses identified 8 wt and 18 *Cyfip1*$^{+/-}$ progeny (Chi-squared p= 0.0125). Backcrosses onto the 129S5/SvEvBrd background were used to maintain the colony and to generate heterozygous and wildtype mice to study.

### *Gda* (CYPIN)

### 1 Mouse generation

#### 1.1 Mutation

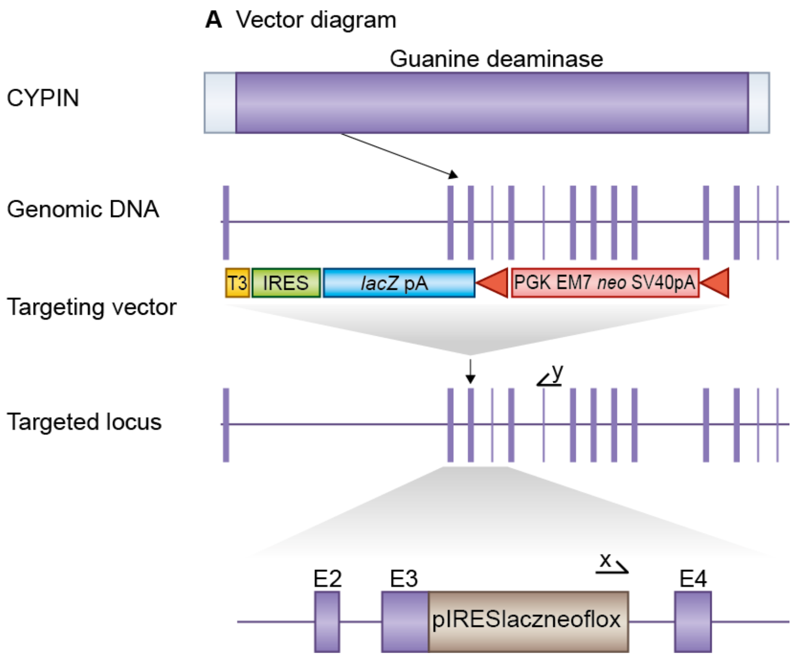


**Figure 1:** *Details of the mutation*

E14TG2a mouse embryonic stem (ES) cells were targeted with a vector containing 8.7kb and 2.7kb of flanking genomic DNA. This replaced 3.1kb of *Gda* genomic DNA (X21503036 to X21506136; Ensemble Build 50) with IRES-lacZ-neo cassette. Correctly targeted ES cells were identified by long range PCR using Expand Long Template PCR system (Roche Cat 11681842001). The PCR contained primer X (5'-GAGCTATTCCAGAAGTAGTGAG-3') and primer Y (5'- CTCAGCTAGAGTGTACTTTAG -3') that correspond to sequence in the IRES-lacZ-neo cassette and sequence outside the 2.7kb flanking region respectively. The correctly targeted ES cells were injected into C57BL/6 blastocysts to create chimeric mice, which were bred with 129S5 mice to generate heterozygous *Gda* mutant mice. Location of *Gda* gene trap. *Gda* is a 14 exon gene encoding a protein that contains a Guanine deaminase domain (top). We replaced most of *Gda* exon 3 with a selection cassette in targeted mice and created a frameshift between exons 3 and 4. Primers used for targeted clone identification (x,y) are shown.

#### 1.2 Genotyping

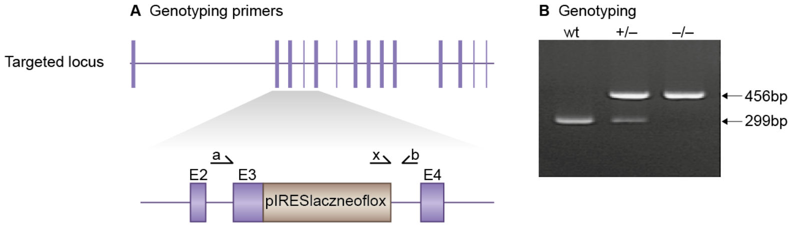


**Figure 2:** *Genotyping details*

Genomic DNA was isolated from ES cells by Wizard SV 96 Genomic DNA purification system (Promega Cat A2371). Genotyping PCR consisted of a 299bp product amplified from the wild-type (wt) allele using a forward primer A (GTCAAGCAAAGACAACCAGAG) in the wt sequence, deleted by targeted mutation, and a reverse primer B (GCTAACTGTCAAGACCTATCC) downstream of the cassette. A 456bp product was amplified from the targeted allele using primer B with forward primer X, within the selection cassette. After enzymatic amplification for 35 cycles (45 seconds at 94 degC, 45 seconds at 55 degC, and 1 minute at 72 degC), the PCR products were size-fractionated on a 2% agarose gel in 1x Tris borate-EDTA buffer. Primers used for genotyping (a,b, x). PCR genotyping of targeted *Gda* mice using a common reverse primer, b, and forward primers a,x to amplify the wt and mutant alleles respectively.

#### 1.3 Breeding

Birth of *Gda*$^{-/-}$ mice followed Mendelian ratios with 20% of offspring being homozygous knockouts. Genotypes of 3-week-old pups from *Gda*$^{+/-}$ intercrosses identified 78 wt, 159 *Gda*$^{+/-}$ and 61 *Gda*$^{-/-}$progeny (Chi-squared p= 0.194). Male and female *Gda*$^{-/-}$ mice showed increased mortality rates between birth and 12 weeks, exhibited small body size but with no gross abnormalities. *Gda* mice were maintained by backcrossing onto the 129S5/ SvEvBrd background; heterozygous males and females were fertile and used to set up intercrosses to generate homozygous and wildtype mice to study.

### *Igsf9* (Dasm1)

#### 1 Mouse generation

##### 1.1 Mutation

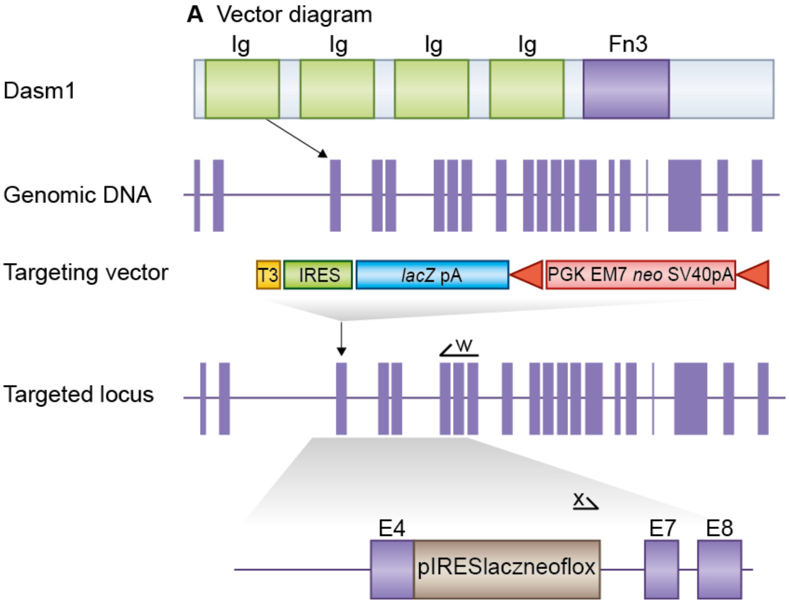


**Figure 1:** *Details of the mutation*

E14TG2a mouse embryonic stem (ES) cells were targeted with a vector containing 7kb and 3.1kb of flanking genomic DNA. This replaced 1.7kb of *Igsf9* genomic DNA (X174419685 to X174421368; Ensemble Build 55) with IRES-lacZ-neo cassette. Correctly targeted ES cells were identified by long range PCR using Expand Long Template PCR system (Roche Cat 11681842001). The PCR contained forward primer X (5'-GAGCTATTCCAGAAGTAGTGAG-3') and reverse primer W (5'- CTAACACTGGCATCTGGTAAG -3') that correspond to sequence in the IRES-lacZ-neo cassette and sequence outside the 3.1kb flanking region respectively. The correctly targeted ES cells were injected into C57BL/6 blastocysts to create chimeric mice, which were bred with 129S5 mice to generate heterozygous *Igsf9* mutant mice. Location of *Igsf9* gene trap. *Igsf9* is a 21 exon gene encoding the Dasm1 protein which contains 4 Ig (immunoglobulin) domains and a Fibronecin type 3 domain (top). We replaced *Igsf9* exons 4-6 with a selection cassette in targeted mice and created a frameshift between exons 3 and 7. Primers used for targeted clone identification (w,x) are shown.

#### 1.2 Genotyping

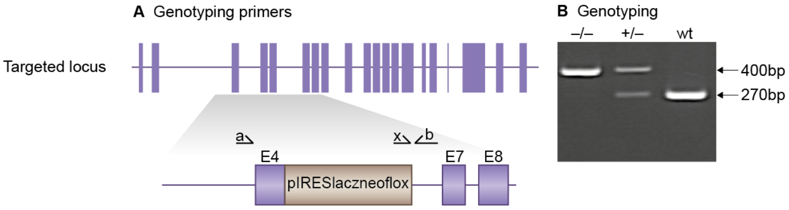


**Figure 2:** *Genotyping details*

Genomic DNA was isolated from ES cells by Wizard SV 96 Genomic DNA purification system (Promega Cat A2371). Genotyping PCR consisted of a 270bp product amplified from the wild-type (wt) allele using a forward primer A (GACATATCGCCCATGTGTG) in the wt sequence deleted by targeted mutation and a reverse primer B (CTTCTGCTGAGAAACTGTATG) downstream of the cassette. A 400bp product was amplified from the targeted allele using primer B with forward primer X, within the selection cassette. After enzymatic amplification for 35 cycles (45 seconds at 94 degC, 45 seconds at 55 degC, and 1 minute at 72 degC), the PCR products were size-fractionated on a 2% agarose gel in 1x Tris borate-EDTA buffer. Primers used for genotyping (a,b, x). PCR genotyping of targeted *Igsf9* mice using a common reverse primer, b, and forward primers a and x to amplify the wt and mutant alleles respectively.

#### 1.3 Breeding

Birth of *Igsf9*$^{-/-}$ mice followed Mendelian ratios with 24% of offspring being homozygous knockouts. Geno-types of 3-week-old pups from *Igsf9*$^{+/-}$ intercrosses identified 30 wt, 93 *Igsf9*$^{+/-}$ and 39 *Igsf9*$^{-/-}$progeny (Chi-squared p= 0.103). Male and female Igsf9$^{-/-}$ mice developed normally to adulthood, exhibited normal body size and no gross abnormalities. *Igsf9* mice were maintained by backcrossing onto the 129S5/SvEvBrd background; heterozygous males and females were fertile and used to set up intercrosses to generate homozygous and wildtype mice to study.

### *Dab2ip* (Dip1/2-E9)

#### 1 Mouse generation

##### 1.1 Mutation

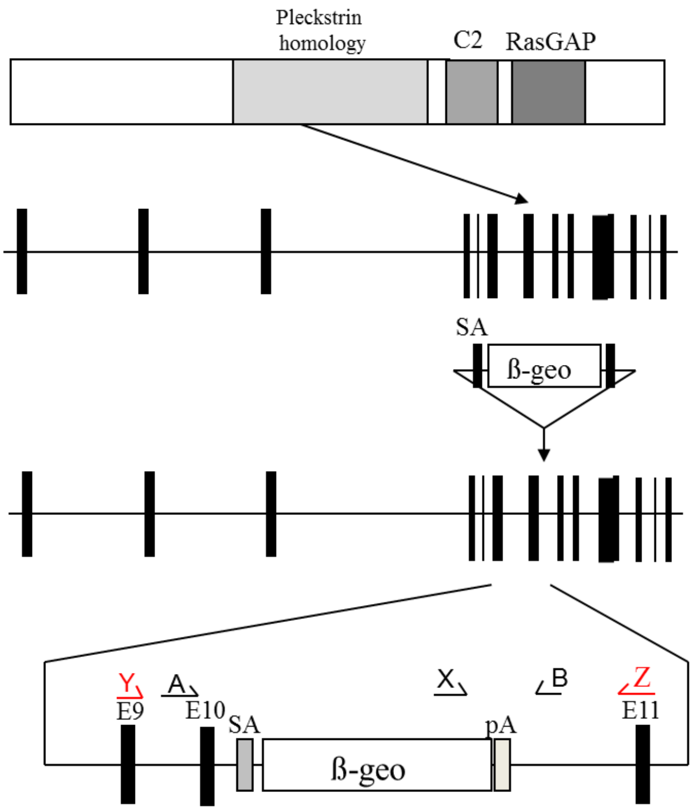


**Figure 1:** *Details of the mutation*

A mouse embryonic stem (ES) cell line (XT0252, strain 129/Ola) with an insertional mutation in *Dab2ip* was obtained from BayGenomics (baygenomics.ucsf.edu/). The insertional mutation in XT0252, by the gene-trapping vector, pGT1lxf, that was designed to create an in-frame fusion between the 5' exons of the trapped gene and a reporter, ß-geo (a fusion of ß-galactosidase and neomycin phosphotransferase II), occurred in intron 10-11. Thus, the gene-trapped locus is predicted to yield a fusion transcript containing exon 1-10 of *Dab2ip* and ß-geo. The ES cells were injected into C57BL/6 blastocysts to create chimeric mice, which were bred with 129S5 mice to generate heterozygous *Dab2ip* mutant mice. Location of *Dab2ip* gene trap. *Dab2ip* is a 16 exon gene encoding a protein which contains Pleckstrin homology, C2 and RasGAP domains (top). The *Dab2ip* gene trap is located in intron 10-11. Primers used for genotyping (A,B, X) and RT-PCR (Y,Z) are shown.

#### 1.2 Genotyping

Genomic DNA was isolated from ES cells or mouse tissues by Wizard SV 96 Genomic DNA purification system (Promega Cat A2371). Genotyping PCR consisted of a 1.6kb product amplified from the wild-type (wt) allele using a forward primer A (5'- GAGACCCTTTCCAACACAGC - 3') upstream of the cassette and a reverse primer B (5'- CCATCTTCTGAAGCCCAGCA - 3') downstream of the cassette. A 1kb product was amplified from the targeted allele using primer B with forward primer X (5'- ATTCAGGCTGCGCAACTGTTGGG - 3'), within the beta-geo cassette. After enzymatic amplification for 35 cycles (45 seconds at 94 degC, 45 seconds at 55 degC, and 2 minutes at 72 degC), the PCR products were size-fractionated on a 1% agarose gel i n 1x Tris borate-EDTA buffer (image not shown).

#### 1.3 Breeding

Birth of *Dab2ip*$^{-/-}$ mice followed Mendelian ratios with 22% of offspring being homozygous knockouts. Genotypes of 3-week-old pups from *Dab2ip*$^{+/-}$ intercrosses i dentified 47 wt, 80 *Dab2ip*$^{+/-}$ and 35 *Dab2ip*$^{-/-}$ progeny (Chi-squared$^2$ p= 0.406). Male and female *Dab2ip*$^{-/-}$ mice developed normally to adulthood, exhibited normal body size and no gross abnormalities. *Dab2ip* mice were maintained by backcrossing onto the 129S5/SvEvBrd background; heterozygous males and females were fertile and used to s et up intercrosses to generate homozygous and wildtype mice to study.

### *Dbn1* (Drebrin)

#### 1 Mouse generation

Drebrin mice were supplied by the Mouse Genome Project (Sanger Institute).

##### 1.1 Mutation

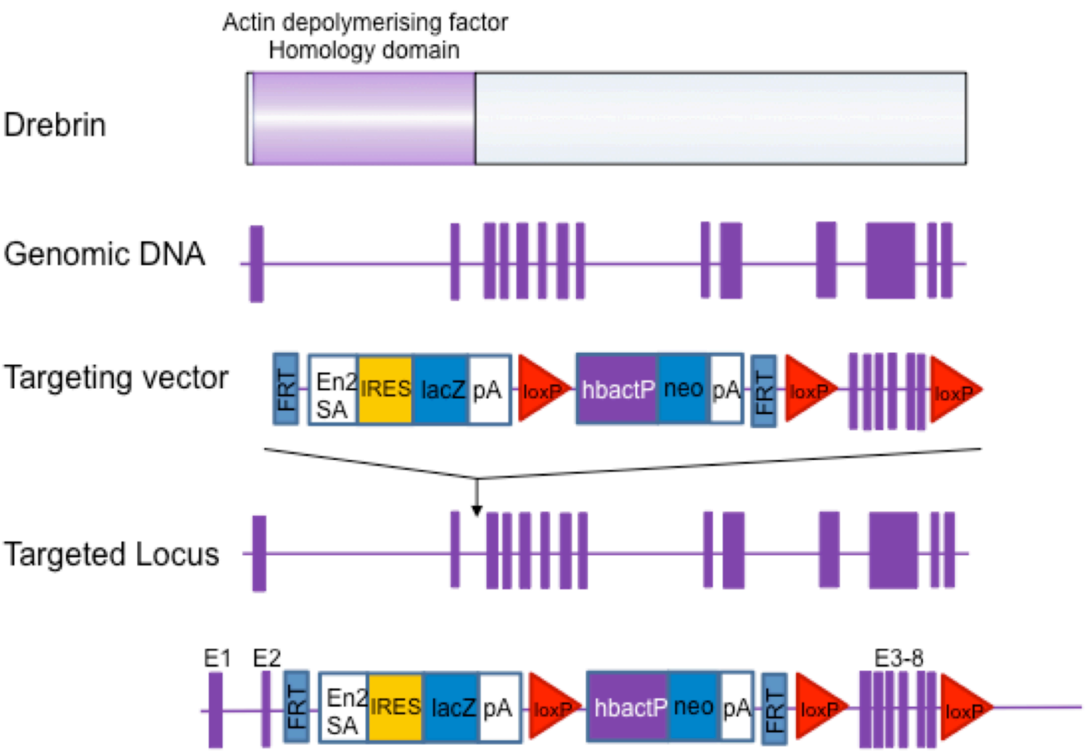


**Figure 1:** *Details of the mutation*

Generation of *Dbn1* targeted mice A. *Dbn1* is a 14 exon gene which encodes an Actin-depolymerising factor homology domain (top). A selection cassette was inserted into intron 2-3 whilst exons 3-8 were floxed creating a frame shift between exons 2 and 9.

#### 1.2 Genotyping

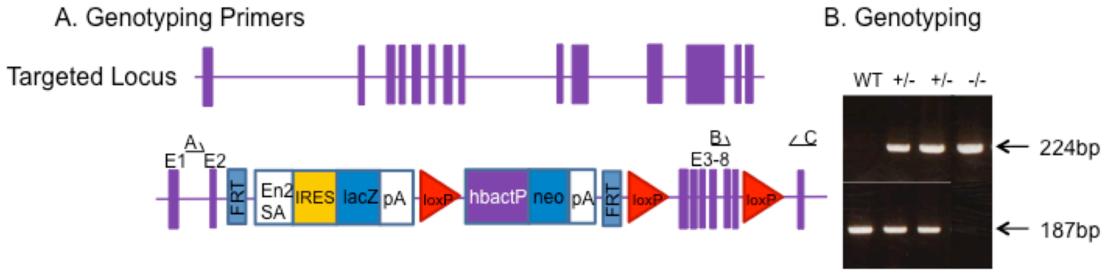


**Figure 2:** *Genotyping details*

Genomic DNA was isolated from ES cells by Wizard SV 96 Genomic DNA purification system (Promega Cat A2371). Genotyping PCR consisted of a 187bp product amplified from the wild-type (wt) allele using a forward primer B (CTATCATGAGTGTGTCAGGTC) and reverse primer C (CAGAGCCCAAGACTAATACAC). A 224bp product was amplified from the targeted allele using forward primer A (CTTGTCCACGGTTGTCCTTC) with reverse primer C. After enzymatic amplification for 35 cycles (45 seconds at 94 °C, 45 seconds at 55 °C, and 1 minute at 72 °C), the PCR products were size-fractionated on a 2% agarose gel in 1x Tris borate-EDTA buffer.

**Primers used for genotyping (A,B & C.** PCR genotyping of targeted Drebrin mice using a common reverse primer, C, and forward primers B and A to amplify the wt and mutant alleles respectively.

#### 1.3 Breeding

Birth of *Dbn1*-/- mice followed Mendelian ratios with 21% of offspring being homozygous knockouts. Genotypes of 3-week-old pups from Drebrin+/- intercrosses identified 60 wt, 95 *Dbn1*+/- and 40 *Dbn1*-/- progeny (chi-squared p= 0.12). Male and female *Dbn1*-/- mice developed normally to adulthood, exhibited normal body size and no gross abnormalities. *Dbn1*+/- mice were maintained by backcrossing onto the C57BL/6J (Charles River) background; heterozygous males and females were fertile and used to set up intercrosses to generate homozygous and wildtype mice to study.

### *Dusp4* (Dusp4)

### 1 Mouse generation

#### 1.1 Mutation

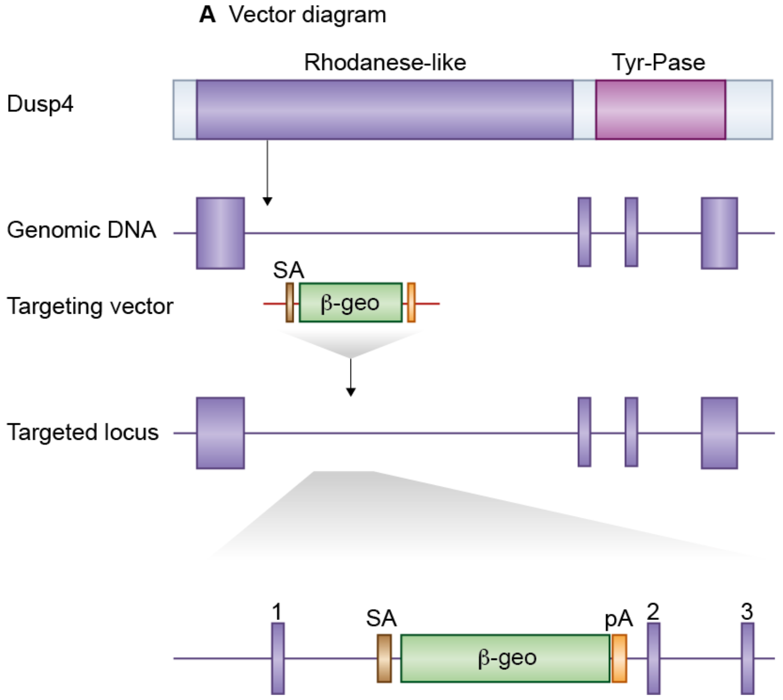


**Figure 1:** *Details of the mutation*

A mouse embryonic stem (ES) cell line (XG164, strain 129/Ola) with an insertional mutation in *Dusp4* was obtained from BayGenomics (baygenomics.ucsf.edu/). The insertional mutation in XG164 by the gene-trapping vector, pGT1lxf, that was designed to create an in-frame fusion between the 5' exon of the trapped gene and a reporter, ß-geo (a fusion of ß-galactosidase and neomycin phosphotransferase II) occurred in intron 1-2. Thus, the gene-trapped locus is predicted to yield a fusion transcript containing exon 1 of *Dusp4* and ß-geo. The ES cells were injected into C57BL/6 blastocysts to create chimeric mice, which were bred with 129S5 mice to generate heterozygous *Dusp4* mutant mice. Location of *Dusp4* gene trap. *Dusp4* is a 4 exon gene that encodes a protein containing a Rhodanese-like domain and a Tyr-Pase domain (top). The *Dusp4* gene trap is located in intron 1-2.

#### 1.2 Genotyping

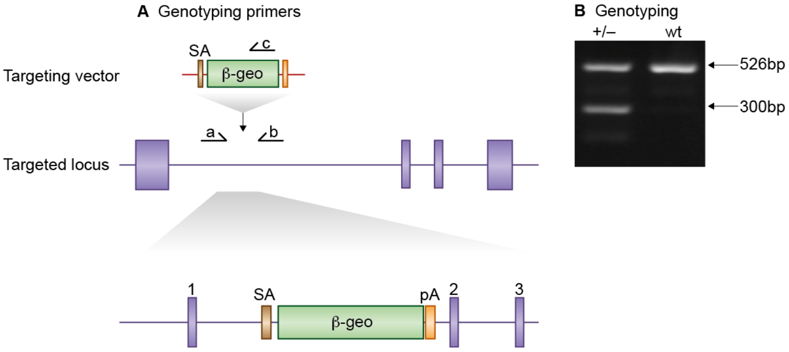


**Figure 2:** *Genotyping details*

Genomic DNA was isolated from ES cells or mouse tissues by Wizard SV 96 Genomic DNA purification system (Promega Cat A2371). Genotyping PCR consisted of a 526bp product amplified from the wild-type (wt) allele using a forward primer a (5'- GGTACATACTAGAAATCTTATG -3') upstream of the cassette and a reverse primer b (5'- GAGTCCTCCACTCAACTCAG -3') in the wt sequence deleted by targeted mutation. A 300bp product was amplified from the targeted allele using primer a with reverse primer c (5'- CAGTC-CTCTTCACATCCATG -3'), within the beta-geo cassette. After enzymatic amplification for 35 cycles (45 seconds at 94 degC, 45 seconds at 55 degC, and 1 minute at 72 degC), the PCR products were size-fractionated on a 2% agarose gel in 1x Tris borate-EDTA buffer. Primers used for genotyping (a,b, c). PCR genotyping of gene trap Dusp4 mice using a common forward primer, a, and reverse primers b and c to amplify the wt and mutant alleles respectively.

#### 1.3 Breeding

No *Dusp4*$^{-/-}$ mice were produced from *Dusp4*$^{+/-}$ intercrosses. Male and female *Dusp4*$^{+/-}$ mice developed normally to adulthood, were fertile, exhibited normal body size and no gross abnormalities. Genotypes of 3-week-old pups from *Dusp4*$^{+/-}$ intercrosses identified 58 wt and 144 Dusp4$^{+/-}$ progeny (Chi-squared p= <0.001). Backcrosses onto the 129S5/SvEvBrd background were used to maintain the colony and to generate heterozygous and wildtype mice to study.

### *Dusp6* (Dusp6)

#### 1 Mouse generation

##### 1.1 Mutation

A. Vector Diagram

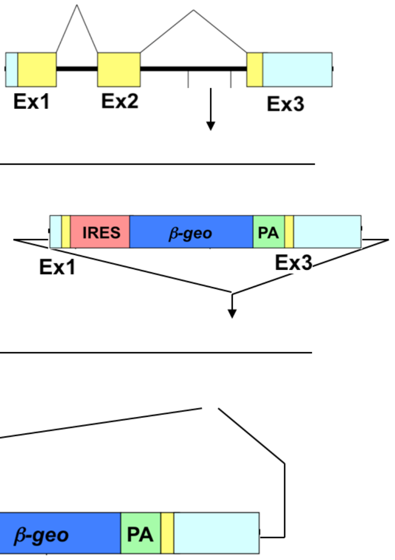


**Figure 1:** *Details of the mutation*

HMI mouse embryonic stem (ES) cells were targeted with a vector for the *Dusp6* locus which comprised 0.7kb and 4.3 kb bands of 5' and 3' genomic DNA flanking an IRES-ß-geo-PA cassette. The targeting vector makes a 2.2 kb deletion in the *Dusp6* locus removing the majority of the gene coding sequence from the OliI site in exon 1, to the BstBI site in exon 3.The correctly targeted ES cells were injected into C57BL/6 blastocysts to create chimeric mice, which were bred with 129S5 mice to generate heterozygous *Dusp6* mutant mice.

#### 1.2 Genotyping

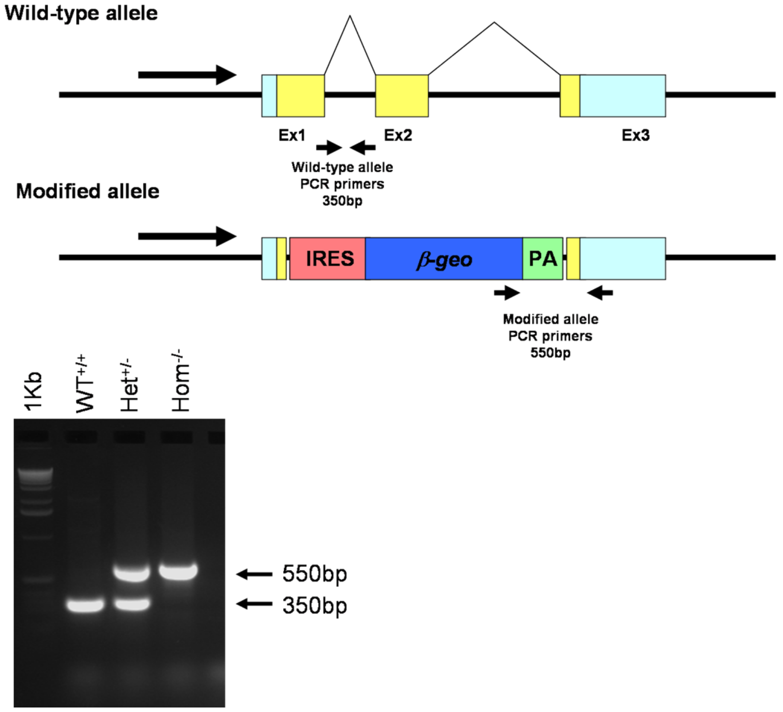


**Figure 2:** *Genotyping details*

Genotyping PCR consisted of a 350bp product amplified from the wild-type (wt) allele using a forward primer A (CAGGCTGGGGAAAATGTTAG) and reverse primer B (AACACTATGGACAGGCACCC), both within intron 1. A 550bp product was amplified from the targeted allele using forward primer C (TCGCCTTCTTGAC-GAGTTCT) and reverse primer D (TGGGTGCCTTTCACGTAGAC). After enzymatic amplification for 30 cycles (30 seconds at 94 degC, 30 seconds at 56 degC, and 1 minute at 72 degC), the PCR products were size-fractionated on a 2% agarose gel in 1x Tris borate-EDTA buffer. Primers used for genotyping (A, B, C and D). PCR genotyping using forward primers A and B in intron 1 gives a 350 bp product from the wt allele. Using primers C (which anneals to beta-geo) and D (annealing to exon 3) a 550bp product was produced from the targeted allele.

#### 1.3 Breeding

Birth of *Dusp6*$^{-/-}$ mice followed Mendelian ratios with 25% of offspring being homozygous knockouts. Genotypes of 3-week-old pups from *Dusp6*$^{+/-}$ intercrosses identified 63 wt, 140 *Dusp6*$^{+/-}$ and 66 *Dusp6*$^{-/-}$progeny (Chi-squared p=0.77). Male and female *Dusp6*$^{-/-}$ mice developed normally to adulthood, exhibited normal body size and no gross abnormalities. *Dusp6* mice were maintained by backcrossing onto the 129S5/SvEvBrd background; heterozygous males and females were fertile and used to set up intercrosses to generate homozygous and wildtype mice to study.

### *Dusp7* (Dusp7)

#### 1 Mouse generation

##### 1.1 Mutation

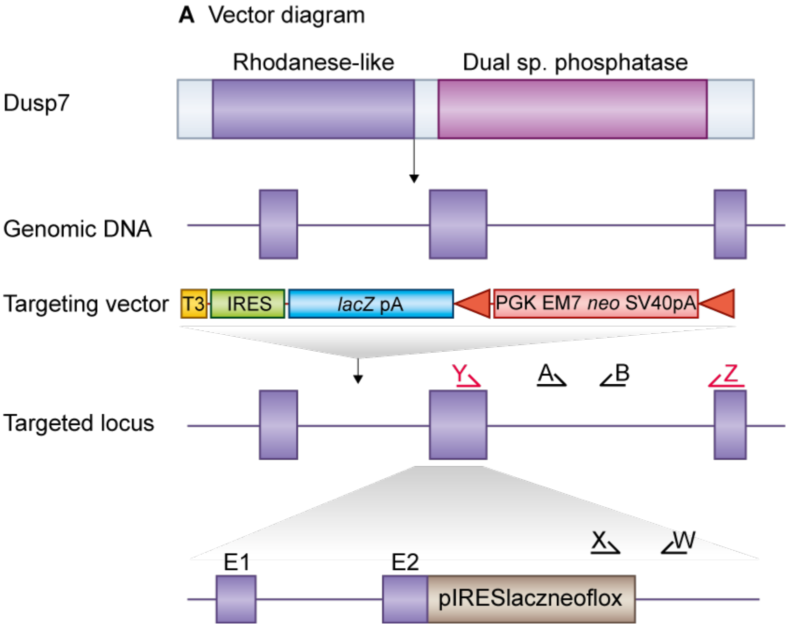


**Figure 1:** *Details of the mutation*

E14TG2a mouse embryonic stem (ES) cells were targeted with a vector containing *Dusp7* flanking genomic DNA sequence. This replaced 3.2kb of *Dusp7* genomic DNA (X106370900 to X106374100; Ensemble Build 75) with I RES-lacZ-neo cassette. Genomic DNA was isolated from ES cells by Wizard SV 96 Genomic DNA purification system (Promega Cat A2371). Correctly targeted ES cells were identified by long range PCR using Expand Long Template PCR system (Roche Cat 11681842001). The correctly targeted ES cells were injected into C57BL/6 blastocysts to create chimeric mice, which were bred with 129S5 mice to generate heterozygous *Dusp7* mutant mice. Generation of *Dusp7* targeted mice. *Dusp7* i s a 3 exon gene encoding a protein which contains Rhodanese-like and Dual specificity phosphatase domains (top). We replaced *Dusp7* exons 2-3 with a s election cassette i n targeted mice which deleted the r est of the gene. Primers used for genotyping (a,b,x, w) and RT-PCR (y,z)

#### 1.2 Genotyping

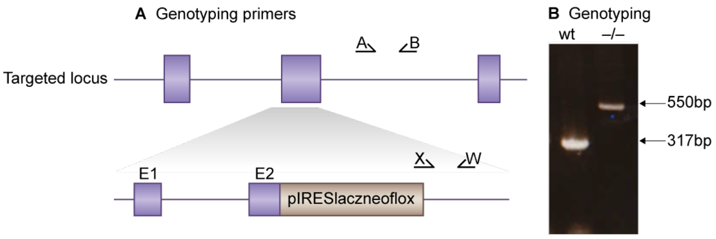


**Figure 2:** *Genotyping details*

PCR genotyping of targeted *Dusp7* mice using primer pairs (A-B) and (X-W) to amplify the wt and mutant alleles respectively. Genotyping PCR consisted of a 317bp product amplified from the wild-type (wt) allele using a forward primer A (GTCTCTGTGGGATCCAGGTG) in the wt sequence deleted by targeted mutation and a reverse primer B (AGCCCAGCTGACACTAAACG) downstream of the cassette. A 550bp product was amplified from the targeted allele using forward primer X (TCGCCTTCTTGACGAGTTCT), within the selection cassette with reverse primer W (GCTATCAGCCGAATGGATGT). After enzymatic amplification for 35 cycles (45 seconds at 94 degC, 45 seconds at 55 degC, and 1 minute at 72 degC), the PCR products were size-fractionated on a 2% agarose gel in 1x Tris borate-EDTA buffer.

#### 1.3 Breeding

Birth of *Dusp7*$^{-/-}$ mice followed Mendelian ratios with 23% of offspring being homozygous knockouts. Genotypes of 3-week-old pups from *Dusp7* $^{+/-}$ intercrosses identified 53 wt, 121 *Dusp7* $^{+/-}$ and 52 *Dusp7*$^{-/-}$progeny (Chi-squared p= 0.565). Male and female *Dusp7*$^{-/-}$ mice developed normally to adulthood, exhibited normal body size and no gross abnormalities *Dusp7* mice were maintained by backcrossing onto the 129S5/SvEvBrd background; heterozygous males and females were fertile and used to set up intercrosses to generate homozygous and wildtype mice to study.

### *Git1* (Git1)

#### 1 Mouse generation

##### 1.1 Mutation

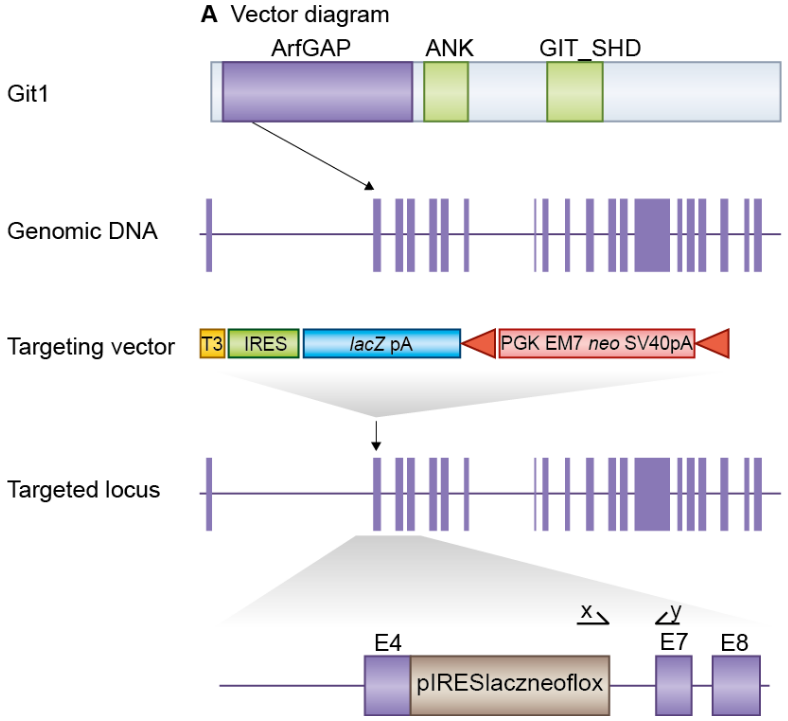


**Figure 1:** *Details of the mutation*

E14TG2a mouse embryonic stem (ES) cells were targeted with a vector containing 6.5kb and 2.7kb of flanking genomic DNA. This replaced 1.1 kb of *Git1* genomic DNA (X77312499 to X77313637; Ensemble Build 55) with IRES-lacZ-neo cassette. Correctly targeted ES cells were identified by long range PCR using Expand Long Template PCR system (Roche Cat 11681842001). The PCR contained primer X (5'-GAGCTATTCCAGAAGTAGTGAG-3') and primer Y (5'- CTGTGTCCTTGCTCTTTACAG -3') that correspond to sequence in the IRES-lacZ-neo cassette and sequence outside the 2.7kb flanking region respectively. The correctly targeted ES cells were injected into C57BL/6 blastocysts to create chimeric mice, which were bred with 129S5 mice to generate heterozygous *Git1* mutant mice. Location of *Git1* gene trap. *Git1* is a 20 exon gene encoding the Git1 protein which contains ArfGAP, Ankyrin and GIT-SHD domains (top). We replaced *Git1* exons 5,6 with a selection cassette in targeted mice and created a frameshift between exons 4 and 7. Primers used for targeted clone identification (x,y) are shown.

#### 1.2 Genotyping

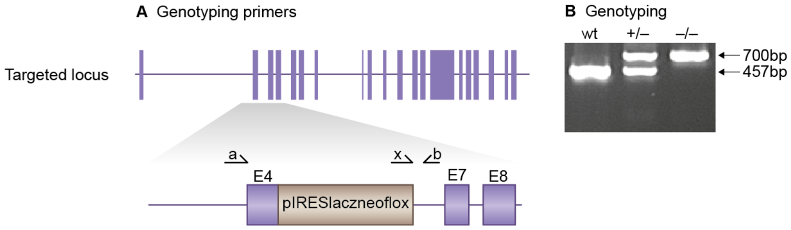


**Figure 2:** *Genotyping details*

Genomic DNA was isolated from ES cells by Wizard SV 96 Genomic DNA purification system (Promega Cat A2371). Genotyping PCR consisted of a 457bp product amplified from the wild-type (wt) allele using a forward primer A (GTCTGCCACTTGAAAGCTAG) in the wt sequence deleted by targeted mutation and a reverse primer B (CACATGTAGAGGTGTAGTGC) downstream of the cassette. A 700bp product was amplified from the targeted allele using primer B with forward primer X, within the selection cassette. After enzymatic amplification for 35 cycles (45 seconds at 94 degC, 45 seconds at 55 degC, and 1 minute at 72 degC), the PCR products were size-fractionated on a 2% agarose gel in 1x Tris borate-EDTA buffer. Primers used for genotyping (a,b, x). PCR genotyping of targeted Git1 mice using a common reverse primer, b, and forward primers a and x to amplify the wt and mutant alleles respectively.

#### 1.3 Breeding

Birth of *Git1*$^{-/-}$ mice did not follow Mendelian ratios with 7% of offspring being homozygous knockouts. Genotypes of 3-week-old pups from *Git1*$^{+/-}$ intercrosses identified 30 wt, 78 *Git1*$^{+/-}$ and 9 *Git1*$^{-/-}$ progeny (Chi-squared p= <0.001). Male and female *Git1*$^{-/-}$ mice developed normally to adulthood, exhibited normal body size but smaller brains and no gross abnormalities. *Git1* mice were maintained by backcrossing onto the 129S5/SvEvBrd background; heterozygous males and females were fertile and used to set up intercrosses to generate homozygous, heterozygous and wildtype mice to study.

### *Dlgap1* (GKAP)

#### 1 Mouse generation

##### 1.1 Mutation

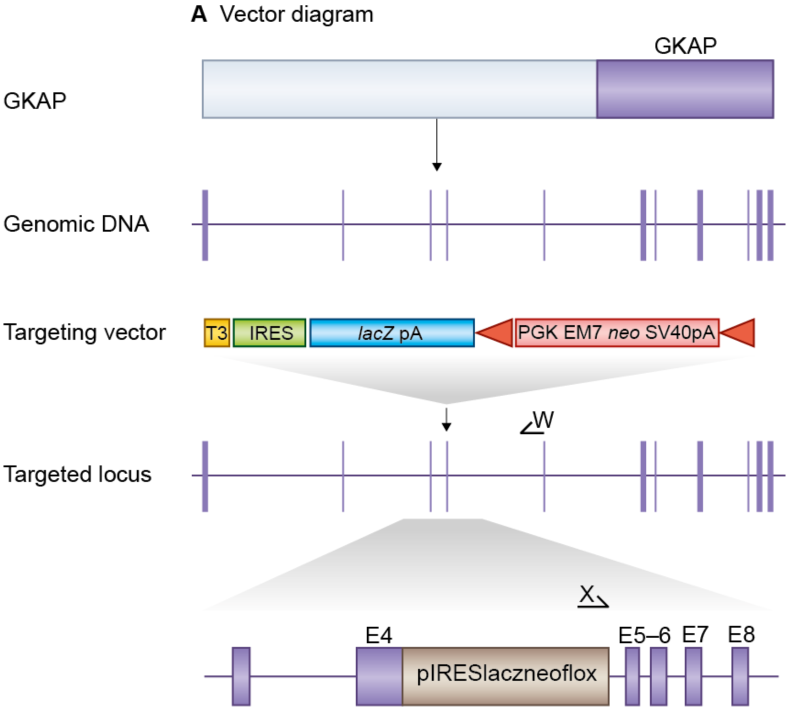


**Figure 1:** *Details of the mutation*

E14TG2a mouse embryonic stem (ES) cells were targeted with a vector containing 5.3kb and 2.6kb of flanking genomic DNA. This replaced 795bp of *Dlgap1* genomic DNA (X70662625 to X70663420; Ensemble Build 68) with IRES-lacZ-neo cassette. Correctly targeted ES cells were identified by long range PCR using Expand Long Template PCR system (Roche Cat 11681842001). The PCR contained primer X (5'-GAGCTATTCCAGAAGTAGTGAG-3') and primer W (5'-GTTGTTGATTGAGAGTGAAGC-3') that correspond to sequence in the IRES-lacZ-neo cassette and sequence outside the 2.6kb flanking region respectively. The correctly targeted ES cells were injected into C57BL/6 blastocysts to create chimeric mice, which were bred with 129S5 mice to generate heterozygous *Dlgap1* mutant mice. Location of *Dlgap1* gene trap. *Dlgap1* is a 11 exon gene encoding the GKAP protein which contains a GKAP domain (top). We replaced *Dlgap1* exon 4 with a selection cassette in targeted mice and created a frameshift between exons 4 and 5.

#### 1.2 Genotyping

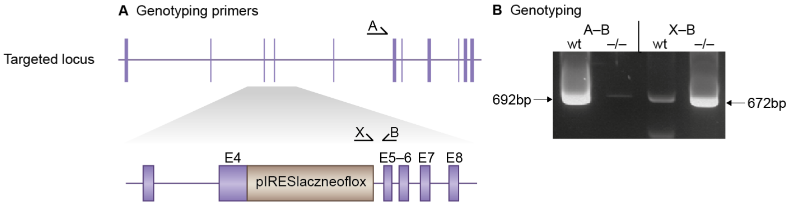


**Figure 2:** *Genotyping details*

Genomic DNA was isolated from ES cells by Wizard SV 96 Genomic DNA purification system (Promega Cat A2371). Genotyping PCR consisted of a 692bp product amplified from the wild-type (wt) allele using a forward primer A (GTGGTCATGGTATGACTCTC) in the wt sequence deleted by targeted mutation and a reverse primer B (GGATATTGTCTGACACTGCTC) downstream of the cassette. A 672bp product was amplified from the targeted allele using primer B with forward primer X, within the selection cassette. After enzymatic amplification for 35 cycles (45 seconds at 94,deg;C, 45 seconds at 55,deg;C, and 1 minute at 72,deg;C), the PCR products were size-fractionated on a 2% agarose gel in 1x Tris borate-EDTA buffer. Primers used for genotyping(a,b, x) PCR genotyping of targeted *Dlgap1* mice using a common reverse primer (b) and reverse primers a and x to amplify the wt and mutant alleles respectively.

#### 1.3 Breeding

Birth of *Dlgap1*$^{-/-}$ mice followed Mendelian ratios with 28% of offspring being homozygous knockouts. Genotypes of 3-week-old pups from *Dlgap1*$^{+/-}$ intercrosses identified 43 wt, 81 Dlgap1$^{+/-}$ and 48 *Dlgap1*$^{-/-}$ progeny (Chi-squared p= 0.647). Male and female *GKAP*$^{-/-}$ mice showed no gross abnormalities. *Dlgap1* mice were maintained by backcrossing onto the 129S5/SvEvBrd background; heterozygous males and females were fertile and used to set up intercrosses to generate homozygous and wildtype mice to study.

### *Dlgap2* (GKAP2)

### 1 Mouse generation

#### 1.1 Mutation

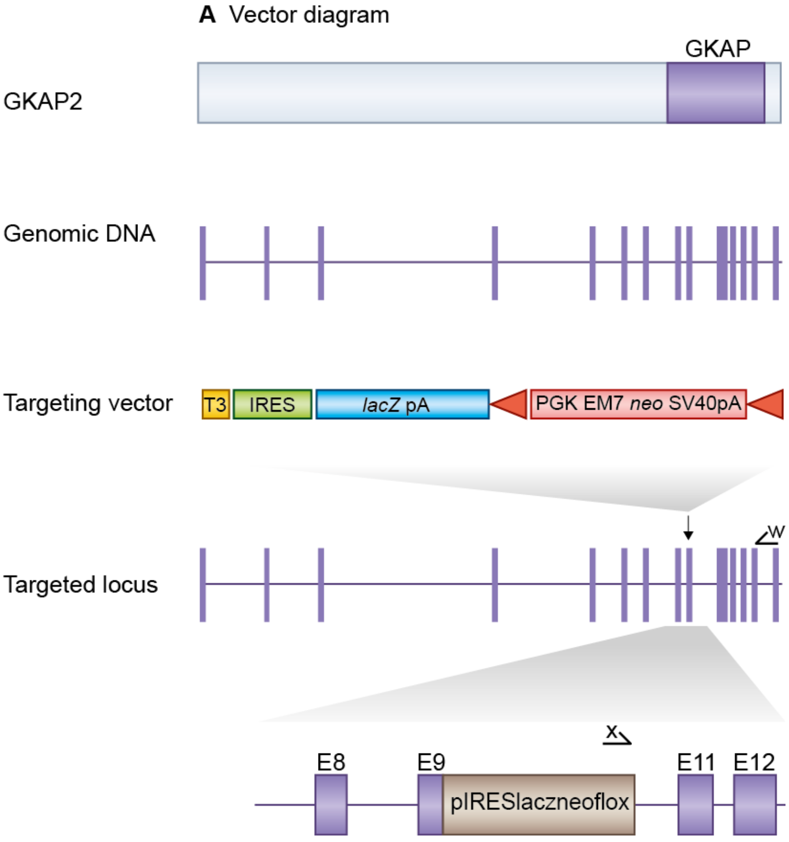


**Figure 1:** *Details of the mutation*

E14TG2a mouse embryonic stem (ES) cells were targeted with a vector containing 6kb and 2.9kb of flanking genomic DNA. This replaced 1.8kb of *Dlgap2* genomic DNA (X14822401 to X14824243; Ensemble Build 69) with IRES-lacZ-neo cassette. Correctly targeted ES cells were identified by long range PCR using Expand Long Template PCR system (Roche Cat 11681842001). The PCR contained forward primer X (5'-GAGCTATTCCAGAAGTAGTGAG-3') and reverse primer W (5'- GCTAGTGGTACAAACACTAGG -3') that correspond to sequence in the IRES-lacZ-neo cassette and sequence outside the 2.9kb flanking region respectively. The correctly targeted ES cells were injected into C57BL/6 blastocysts to create chimeric mice, which were bred with 129S5 mice to generate heterozygous *Dlgap2* mutant mice. Those F1 heterozygous mice had been backcrossed with 129S5 mice for 1-2 times before being used for intercrossing. Location of *Dlgap2* gene trap. *Dlgap2* is a 15 exon gene encoding the GKAP2 protein which contains a GKAP domain (top). We replaced *Dlgap2* exons 9-10 with a selection cassette in targeted mice and created a frameshift between exons 8 and 11. Primers used for targeted clone identification (w,x) are shown.

#### 1.2 Genotyping

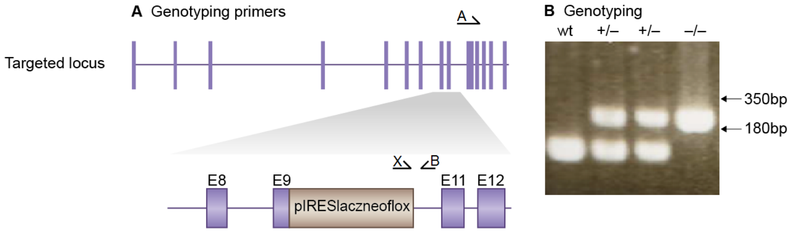


**Figure 2:** *Genotyping details*

Genomic DNA was isolated from ES cells by Wizard SV 96 Genomic DNA purification system (Promega Cat A2371). Genotyping PCR consisted of a 180bp product amplified from the wild-type (wt) allele using a forward primer A (CTATCCTCCCTCACTCTGTTG) in the wt sequence deleted by targeted mutation and a reverse primer B (GCATCTTGATGTTCACACATC) downstream of the cassette. A 350bp product was amplified from the targeted allele using primer B with forward primer X, within the selection cassette. After enzymatic amplification for 35 cycles (45 seconds at 94 degC, 45 seconds at 55 degC, and 1 minute at 72 degC), the PCR products were size-fractionated on a 2% agarose gel in 1x Tris borate-EDTA buffer. Primers used for genotyping (A,B, X). PCR genotyping of targeted *Dlgap2* mice using a common reverse primer, B, and forward primers A and X to amplify the wt and mutant alleles respectively.

#### 1.3 Breeding

Birth of *Dlgap2*$^{-/-}$ mice followed Mendelian ratios with 26% of offspring being homozygous knockouts. Genotypes of 3-week-old pups from *Dlgap2*$^{+/-}$ intercrosses identified 22 wt, 57 *Dlgap2*$^{+/-}$ and 28 *Dlgap2*$^{-/-}$ progeny (Chi-squared p= 0.568). Male and female *Dlgap2*$^{-/-}$ mice developed normally to adulthood, exhibited normal body size and no gross abnormalities. *Dlgap2* mice were maintained by backcrossing onto the 129S5/SvEvBrd background; heterozygous males and females were fertile and used to set up intercrosses to generate homozygous and wildtype mice to study.

### *Dlgap4* (GKAP4)

#### 1 Mouse generation

##### 1.1 Mutation

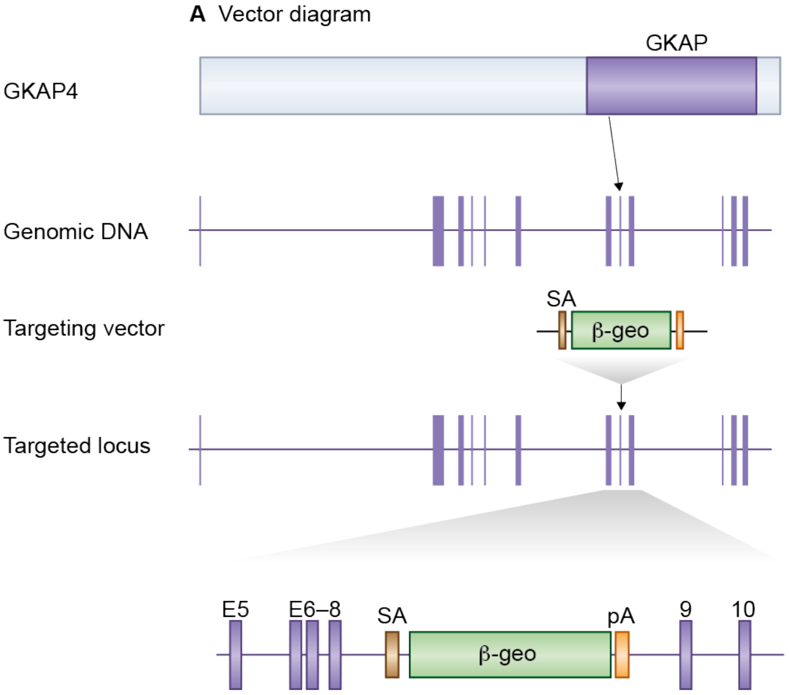


**Figure 1:** *Details of the mutation*

A mouse embryonic stem (ES) cell line (CA0132, strain 129/Ola) with an insertional mutation in *Dlgap4* was obtained from Sanger Institute Gene Trap Resource (SIGTR - sanger.ac.uk/PostGenomics/genetrap/). The insertional mutation in CA0132, by the gene-trapping vector, pGT0lxr, that was designed to create an in-frame fusion between the 5' exons of the trapped gene and a reporter, ß-geo (a fusion of ß-galactosidase and neomycin phosphotransferase II), occurred in intron 8-9. Thus, the gene-trapped locus is predicted to yield a fusion transcript containing exons 1-8 of *Dlgap4* and ß-geo. The ES cells were injected into C57BL/6 blastocysts to create chimeric mice, which were bred with 129S5 mice to generate heterozygous *Dlgap4*-mutant mice. Location of *Dlgap4* gene trap. *Dlgap4* is a 13 exon gene encoding a protein that contains a GKAP domain (top). The *Dlgap4* gene trap is located in intron 8-9.

#### 1.2 Genotyping

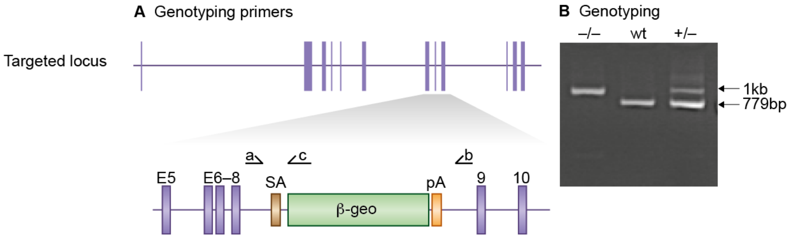


**Figure 2:** *Genotyping details*

Genomic DNA was isolated from ES cells or mouse tissues by Wizard SV 96 Genomic DNA purification system (Promega Cat A2371). Genotyping PCR consisted of a 779bp product amplified from the wild-type (wt) allele using a forward primer A (5'- GGTTCTTTTACTCAGAGCATG -3') in the wt sequence deleted by targeted mutation and a reverse primer B (5'- CAAACAGACATGTGGACAGAC -3') upstream of the cassette. A 1kb product was amplified from the targeted allele using primer A with reverse primer C (5'-GGTTACGTTGGTGTAGATGG -3'), within the beta-geo cassette. After enzymatic amplification for 35 cycles (45 seconds at 94 degC, 45 seconds at 55 degC, and 1 minute at 72 degC), the PCR products were size-fractionated on a 2% agarose gel in 1x Tris borate-EDTA buffer. Primers used for genotyping (a,b, c). PCR genotyping of gene trap GKAP4 mice using a common forward primer, a, and reverse primers b and c to amplify the wt and mutant alleles respectively.

#### 1.3 Breeding

Birth of *Dlgap4*$^{-/-}$ mice did not follow Mendelian ratios with 30% of offspring being homozygous knockouts. Genotypes of 3-week-old pups from *Dlgap4*$^{+/-}$ intercrosses identified 21 wt, 85 *Dlgap4*$^{+/-}$ and 46 *Dlgap4*$^{-/-}$ progeny (Chi-squared p= 0.006). Male and female *Dlgap4*$^{-/-}$ mice developed normally to adulthood, exhibited normal body size and no gross abnormalities. *Dlgap4* mice were maintained by backcrossing onto the 129S5/SvEvBrd background; heterozygous males and females were fertile and used to set up intercrosses to generate homozygous and wildtype mice to study.

### *Glul* (Glul)

#### 1 Mouse generation

##### 1.1 Mutation

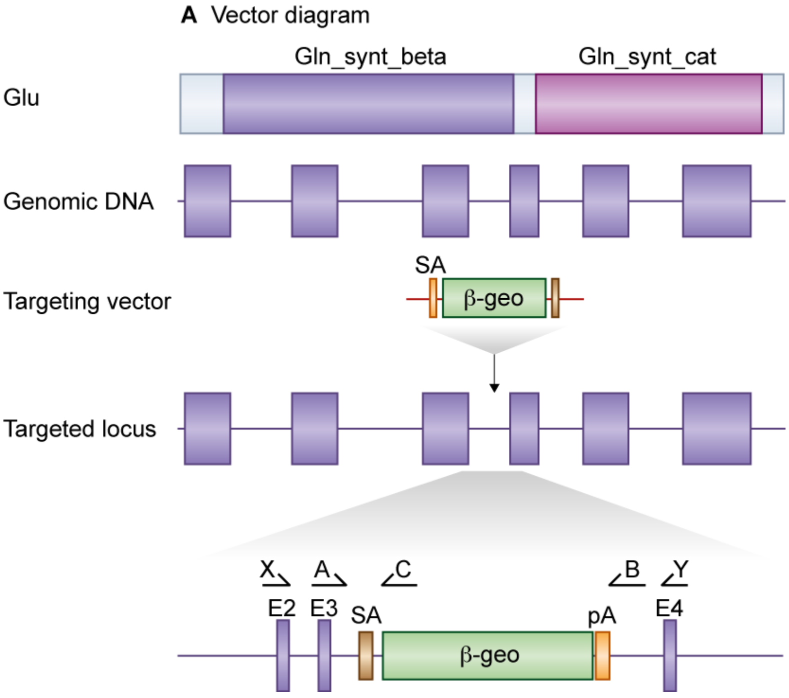


**Figure 1:** *Details of the mutation*

A mouse embryonic stem (ES) cell line (HMA290, strain 129/Ola) with an insertional mutation in *Glul* was obtained from BayGenomics (baygenomics.ucsf.edu/). The insertional mutation in CE0101, by the gene-trapping vector, pGT1lxf, that was designed to create an in-frame fusion between the 5' exons of the trapped gene and a reporter, ß-geo (a fusion of ß-galactosidase and neomycin phosphotransferase II), occurred within intron 3-4. Thus, the gene-trapped locus is predicted to yield a fusion transcript containing exons 1-3 of Glul and ß-geo. The ES cells were injected into C57BL/6-J blastocysts to create chimeric mice, which were bred with 129S5 mice to generate heterozygous *Glul* mutant mice. Location of *Glul* gene trap. *Glul* is a 7 exon gene encoding a protein which contains Gln synthetase beta-Grasp and Gln synthetase catalytic domains (top). The Glul gene trap is located in intron 3-4. Primers used for genotyping (A,B,C) and RT-PCR (X,Y) are shown

#### 1.2 Genotyping

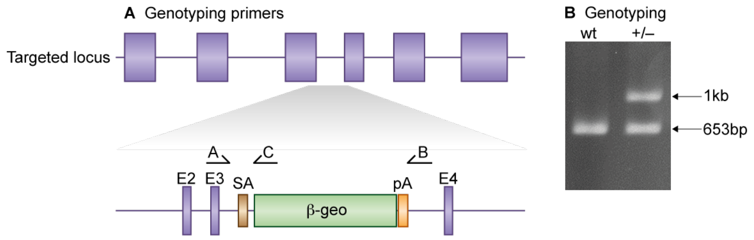


**Figure 2:** *Genotyping details*

Genomic DNA was isolated from ES cells or mouse tissues by Wizard SV 96 Genomic DNA purification system (Promega Cat A2371). Genotyping PCR consisted of a 653bp product amplified from the wild-type (wt) allele using a forward primer A (5'- CATGTACCTCCATCCTGTTG -3') and a reverse primer B (5'- CACGGTCATTCATGTGATC -3'). A 1kb product was amplified from the targeted allele using primer A with reverse primer C (5'- GATCCTCTAGAGTCCAGATCTG -3'), within the ß-geo cassette. After enzymatic amplification for 35 cycles (45 seconds at 94 degC, 45 seconds at 55 degC, and 1 minute at 72 degC), the PCR products were size-fractionated on a 2% agarose gel in 1x Tris borate-EDTA buffer.

#### 1.3 Breeding

No *Glul*$^{-/-}$ mice were produced from *Glul*$^{+/-}$ intercrosses. Male and female *Glul*$^{+/-}$ mice developed normally to adulthood, were fertile, exhibited normal body size and no gross abnormalities. Genotypes of 3-week-old-pups from *Glul*$^{+/-}$ intercrosses identified 95 wt, and 147 *Glul*$^{+/-}$ progeny (Chi-squared $p= <0.001$). Backcrosses onto the 129S5/SvEvBrd background were used to maintain the colony and to generate heterozygous and wildtype mice to study.

### *Gria1* (GluR1)

#### 1 Mouse generation

##### 1.1 Mutation

Generation of *Gria1* mutant mice described in PMID:10364547, Zamanillo *et al* (1999)

##### 1.2 Genotyping

*Gria1* genotyping described in PMID:10364547, Zamanillo *et al* (1999)

##### 1.3 Breeding

Birth of GluR1-/- mice did not follow Mendelian ratios with 17% of offspring being homozygous knockouts. Genotypes from GluR1+/- intercrosses identified 105 wt, 171 GluR1+/- and 57 GluR1-/- progeny (Chi-squared p= 0.001). Male and female GluR1-/- mice developed normally to adulthood, exhibited normal body size and no gross abnormalities. GluR1 mice were maintained by backcrossing onto the C57BL/6J background; heterozygous males and females were fertile and used to set up intercrosses to generate homozygous and wildtype mice to study.

### *Gnb1* (Gnb1)

#### 1 Mouse generation

##### 1.1 Mutation

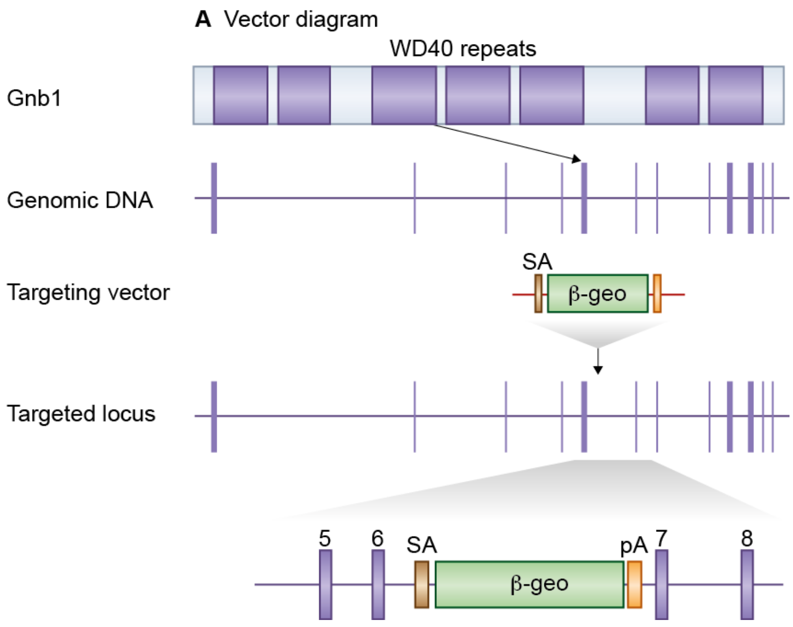


**Figure 1:** *Details of the mutation*

A mouse embryonic stem (ES) cell line (AN0901, strain 129/Ola) with an insertional mutation in *Gnb1* was obtained from Sanger Institute Gene Trap Resource (SIGTR - sanger.ac.uk/PostGenomics/genetrap/). The insertional mutation in AN0901 by the gene-trapping vector, pGT0lxr, that was designed to create an in-frame fusion between the 5' exons of the trapped gene and a reporter, ß-geo (a fusion of ß-galactosidase and neomycin phosphotransferase II) occurred in intron 6-7. Thus, the gene-trapped locus is predicted to yield a fusion transcript containing exons 1-6 of *Gnb1* and ß-geo. The ES cells were injected into C57/BL6 blastocysts to create chimeric mice, which were bred with 129S5 mice to generate heterozygous *Gnb1*-mutant mice. Location of *Gnb1* gene trap. *Gnb1* is a 12 exon protein and contains a WD40 repeat domain (top). The GNB1 gene trap is located in intron 6-7.

#### 1.2 Genotyping

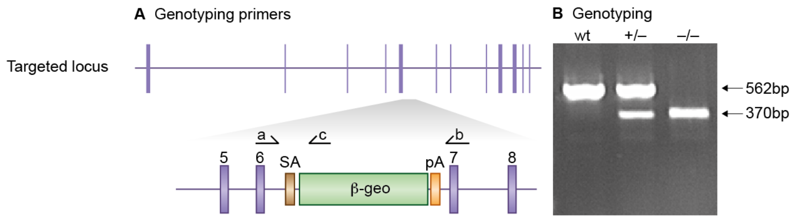


**Figure 2:** *Genotyping details*

Genomic DNA was isolated from ES cells or mouse tissues by Wizard SV 96 Genomic DNA purification system (Promega Cat A2371). Genotyping PCR consisted of a 562bp product amplified from the wild-type (wt) allele using a forward primer A (5'- GAGTATCTGAGGATAGCTAC -3') upstream of the cassette and a reverse primer B (5'- CTGCAACCACTACGCAGAC -3') in the wt sequence deleted by targeted mutation. A 370bp product was amplified from the targeted allele using primer A with reverse primer C (5'- CAGTCCTCTTCA-CATCCATG -3'), within the ß-geo cassette. After enzymatic amplification for 35 cycles (45 seconds at 94 degC, 45 seconds at 55 degC, and 1 minute at 72 degC), the PCR products were size-fractionated on a 2%agarose gel in 1x Tris borate-EDTA buffer. Primers used for genotyping (a,b, c). PCR genotyping of gene trap Gnb1 mice using a common forward primer, a, and reverse primers b and c to amplify the wt and mutant alleles respectively.

#### 1.3 Breeding

Birth of *Gnb1*$^{-/-}$ mice did not follow Mendelian ratios with 17% of offspring being homozygous knockouts. Genotypes from *Gnb1*$^{+/-}$ intercrosses identified 53 wt, 148 *Gnb1*$^{+/-}$ and 41 *Gnb1*$^{-/-}$ progeny (Chi-squared p= 0.001). Male and female *Gnb1*$^{-/-}$ mice developed normally to adulthood, exhibited normal body size and no gross abnormalities. *Gnb1* mice were maintained by backcrossing onto the 129S5/SvEvBrd background; heterozygous males and females were fertile and used to set up intercrosses to generate homozygous and wildtype mice to study.

### *Arhgap32* (GRIT)

#### 1 Mouse generation

##### 1.1 Mutation

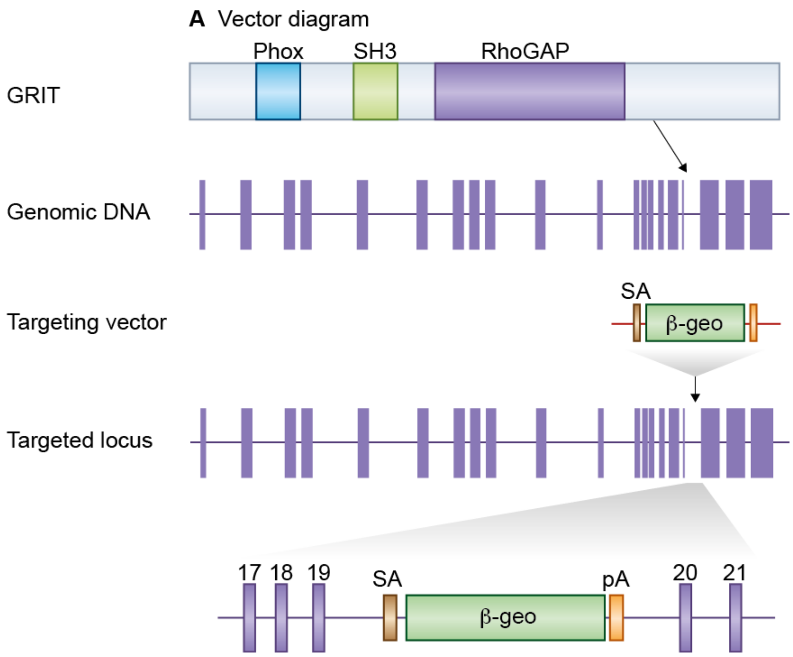


**Figure 1:** *Details of the mutation*

A mouse embryonic stem (ES) cell line (XG279, strain 129/Ola) with an insertional mutation in *Arhgap32* was obtained from BayGenomics (baygenomics.ucsf.edu/). The insertional mutation in XG279 by the gene-trapping vector, pGT1lxf, that was designed to create an in-frame fusion between the 5' exons of the trapped gene and a reporter, ß-geo (a fusion of ß-galactosidase and neomycin phosphotransferase II) occurred in intron 19-20. Thus, the gene-trapped locus is predicted to yield a fusion transcript containing exons 1-19 of *Arhgap32* and ß-geo. The ES cells were injected into C57BL/6 blastocysts to create chimeric mice, which were bred with 129S5 mice to generate heterozygous *Arhgap32* mutant mice. Location of *Arhgap32* gene trap. *Arhgap32* is a 22 exon gene encoding the GRIT protein which contains Phox, SH3 and RhoGAP domains (top). The *Arhgap32* gene trap is located in intron 19-20.

#### 1.2 Genotyping

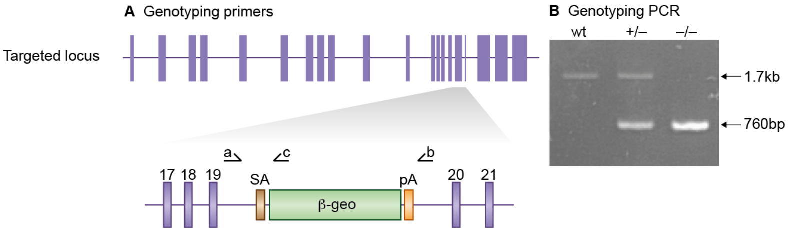


**Figure 2:** *Genotyping details*

Genomic DNA was isolated from ES cells or mouse tissues by Wizard SV 96 Genomic DNA purification system (Promega Cat A2371). Genotyping PCR consisted of a 1.7kb product amplified from the wild-type (wt) allele using a forward primer A (5'- GTCCCTTACTTCTCTCCATG -3') upstream of the cassette and a reverse primer B (5'- CTGAGCTACAGTATTGGAGAC -3') downstream of the trap. A 760bp product was amplified from the targeted allele using primer A with reverse primer C (5'- CATCCACTACTCAGTGCAGTG -3'), within the ß-geo cassette. After enzymatic amplification for 30 cycles (45 seconds at 94 degC, 1 minute at 59 degC, and 2 minutes at 72 degC), the PCR products were size-fractionated on a 1% agarose gel in 1x Tris borate-EDTA buffer. Primers used for genotyping (a,b, c). PCR genotyping of gene trap GRIT mice using a common forward primer, a, and reverse primers b and c to amplify the wt and mutant alleles respectively.

#### 1.3 Breeding

Birth of *Arhgap32*$^{-/-}$ mice followed Mendelian ratios with 27% of offspring being homozygous knockouts. Genotypes of 3-week-old pups from *Arhgap32*$^{+/-}$ intercrosses identified 69 wt, 93 *Arhgap32*$^{+/-}$ and 61 *Arhgap32*$^{-/-}$ progeny (Chi-squared p= 0.035). Male and female *Arhgap32*$^{-/-}$ mice developed normally to adulthood, exhibited normal body size and no gross abnormalities. *Arhgap32* mice were maintained by backcrossing onto the C57BL/6J back-ground; heterozygous males and females were fertile and used to set up intercrosses to generate homozygous and wildtype mice to study.

### *Hras* (H-Ras)

#### 1 Mouse generation

##### 1.1 Mutation

Generation of *Hras* mutant mice described in PMID:12427827, Komiyama *et al* (2002)

##### 1.2 Genotyping

*Hras* genotyping described in PMID:12427827, Komiyama *et al* (2002)

##### 1.3 Breeding

Birth of *H-Ras*$^{-/-}$ mice followed Mendelian ratios with 25% of offspring being homozygous knockouts. Genotypes of 3-week-old pups from *H-Ras*$^{+/-}$ intercrosses identified 81 wt, 129 *H-Ras*$^{+/-}$ and 70 *H-Ras*$^{-/-}$ progeny (Chi-squared p= 0.274). Male and female *H-Ras*$^{-/-}$ mice developed normally to adulthood, exhibited normal body size and no gross abnormalities. *H-Ras* mice were maintained by backcrossing onto the C57BL/6J background; heterozygous males and females were fertile and used to set up intercrosses to generate homozygous and wildtype mice to study.

### *Iqsec1* (Iqsec1)

### 1 Mouse generation

#### 1.1 Mutation

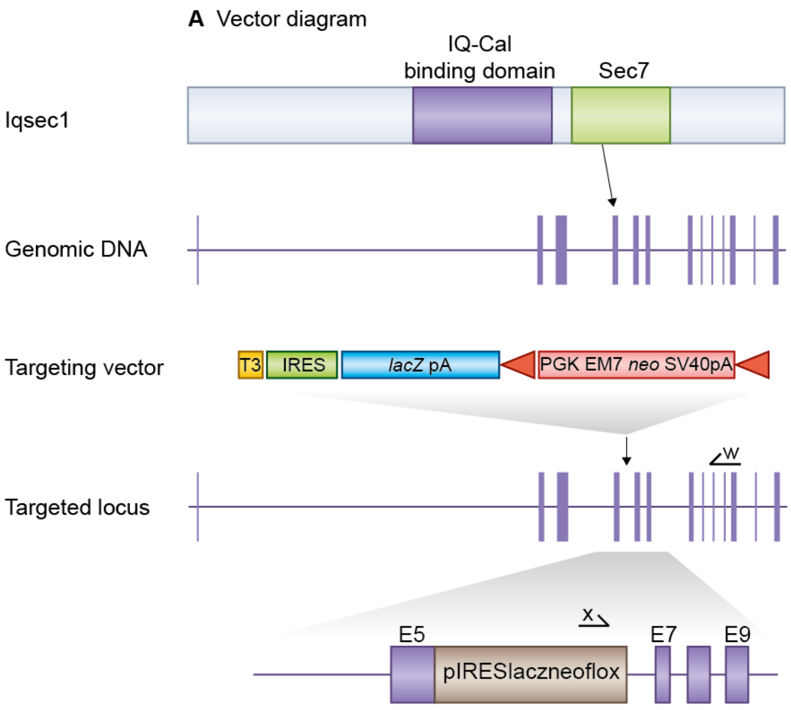


**Figure 1:** *Details of the mutation*

E14TG2a mouse embryonic stem (ES) cells were targeted with a vector containing 6.1kb and 3.2kb of flanking genomic DNA. This replaced 1.5kb of *Iqsec1* genomic DNA (X90628029 to X90629529; Ensemble Build 50) with IRES-lacZ-neo cassette. Correctly targeted ES cells were identified by long range PCR using Expand Long Template PCR system (Roche Cat 11681842001). The PCR contained primer X (5'-GAGCTATTCCAGAAGTAGTGAG-3') and primer W (5'- CCACTCAGACATCATACTGTG -3') that correspond to sequence in the IRES-lacZ-neo cassette and sequence outside the 3.2kb flanking region respectively. The correctly targeted ES cells were injected into C57BL/6 blastocysts to create chimeric mice, which were bred with 129S5 mice to generate heterozygous *Iqsec1* mutant mice. Location of *Iqsec1* gene trap. *Iqsec1* is a 14 exon gene encoding the Iqsec1 protein which contains a IQ-Cal binding domain and Sec7 domain (top). We replaced *Iqsec1* exon 6 with a selection cassette in targeted mice and created a frameshift between exons 5 and 7. Primers used for targeted clone identification (w,x) are shown.

#### 1.2 Genotyping

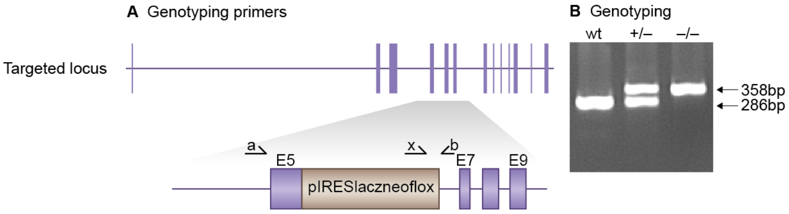


**Figure 2:** *Genotyping details*

Genomic DNA was isolated from ES cells by Wizard SV 96 Genomic DNA purification system (Promega Cat A2371). Genotyping PCR consisted of a 286bp product amplified from the wild-type (wt) allele using a forward primer A (CAGACACCATCTTCATCCTG) in the wt sequence, deleted by targeted mutation, and a reverse primer B (CACTTCACTCTGTTGAGCATG) downstream of the cassette. A 358bp product was amplified from the targeted allele using primer B with forward primer X, within the selection cassette. After enzymatic amplification for 35 cycles (45 seconds at 94 degC, 45 seconds at 55 degC, and 1 minute at 72 degC), the PCR products were size-fractionated on a 2% agarose gel in 1x Tris borate-EDTA buffer. Primers used for genotyping (a,b, x). PCR genotyping of targeted Iqsec1 mice using a common reverse primer, b, and forward primers a and x to amplify the wt and mutant alleles respectively.

#### 1.3 Breeding

Birth of *Iqsec1*$^{-/-}$ mice did not follow Mendelian ratios with 11% of offspring being homozygous knockouts. Genotypes of 3-week-old pups from *Iqsec1*$^{+/-}$ intercrosses identified 45 wt, 76 *Iqsec1*$^{+/-}$ and 15 *Iqsec1*$^{-/-}$ progeny (Chi-squared p= 0.001). Male and female Iqsec1$^{-/-}$ mice developed normally to adulthood, exhibited normal body size and no gross abnormalities. *Iqsec1* mice were maintained by backcrossing onto the 129S5/SvEvBrd background; heterozygous males and females were fertile and used to set up intercrosses to generate homozygous, heterozygous and wildtype mice to study.

### *Iqsec3* (Iqsec3)

#### 1 Mouse generation

##### 1.1 Mutation

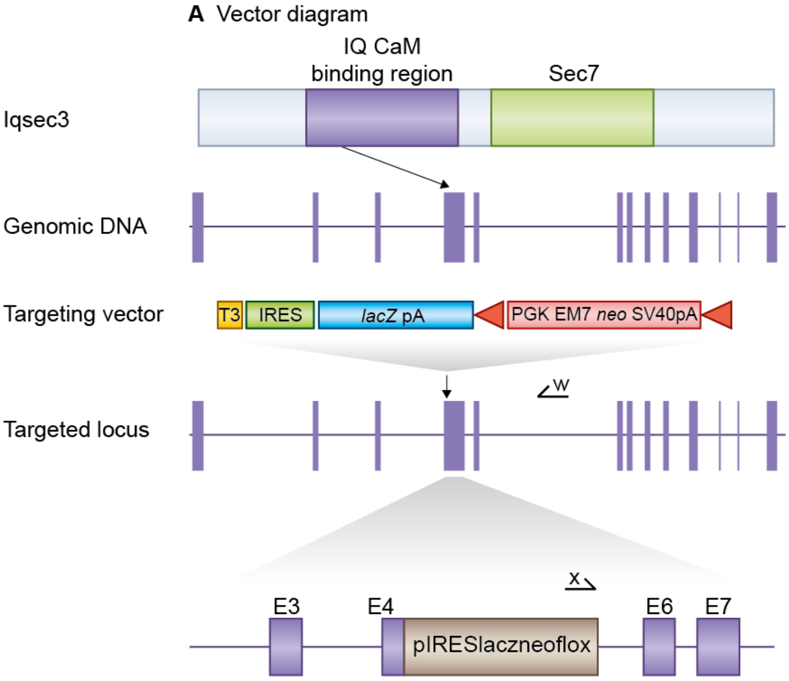


**Figure 1:** *Details of the mutation*

E14TG2a mouse embryonic stem (ES) cells were targeted with a vector containing 6.9kb and 3.1kb of flanking genomic DNA. This replaced 3kb of *Iqsec3* genomic DNA (X121360567 to X121363526; Ensemble Build 55) with IRES-lacZ-neo cassette. Correctly targeted ES cells were identified by long range PCR using Expand Long Template PCR system (Roche Cat 11681842001). The PCR contained primer X (5'-GAGCTATTCCAGAAGTAGTGAG-3') and primer W (5'- CTCATCACTGTGATCAAACAC -3') that correspond to sequence in the IRES-lacZ-neo cassette and sequence outside the 3.1kb flanking region respectively. The correctly targeted ES cells were injected into C57BL/6 blastocysts to create chimeric mice, which were bred with 129S5 mice to generate heterozygous *Iqsec3* mutant mice. Location of *Iqsec3* gene trap. *Iqsec3* is a 14 exon gene, encoding the Iqsec3 protein which contains a IQ CAM binding region and Sec7 domain(top). We replaced exons 4 and 5 with a selection cassette in targeted mice and created a frameshift between exons 3 and 6. Primers used for targeted clone identification (w,x) are shown.

#### 1.2 Genotyping

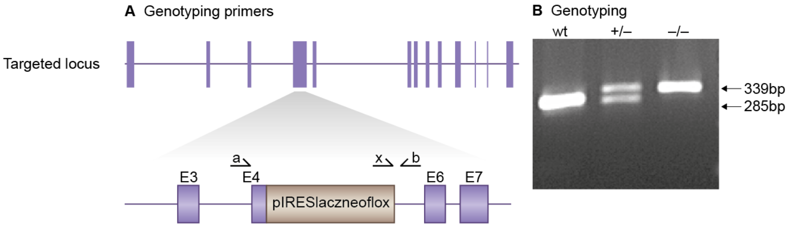


**Figure 2:** *Genotyping details*

Genomic DNA was isolated from ES cells by Wizard SV 96 Genomic DNA purification system (Promega Cat A2371). Genotyping PCR consisted of a 285bp product amplified from the wild-type (wt) allele using a forward primer A (CATCCAGTTCCTGATCTCAC) in the wt sequence deleted by targeted mutation and a reverse primer B (CCATCAAATGAATGACTGCAC) downstream of the cassette. A 339bp product was amplified from the targeted allele using primer B with forward primer X, within the selection cassette. After enzymatic amplification for 35 cycles (45 seconds at 94 degC, 45 seconds at 55 degC, and 1 minute at 72 degC), the PCR products were size-fractionated on a 2% agarose gel in 1x Tris borate-EDTA buffer. Primers used for genotyping (a,b, x). PCR genotyping of targeted Iqsec3 mice using a common reverse primer, b, and forward primers a and x to amplify the wt and mutant alleles respectively.

#### 1.3 Breeding

Birth of *Iqsec3*$^{-/-}$ mice did not follow Mendelian ratios with 16% of offspring being homozygous knockouts. Genotypes of 3-week-old pups from *Iqsec3*$^{+/-}$ intercrosses identified 132 wt, 176 *Iqsec3*$^{+/-}$ and 59 *Iqsec3*$^{-/-}$ progeny (Chi-squared p= <0.001). Male and female *Iqsec3*$^{-/-}$ mice developed normally to adulthood, exhibited normal body size and no gross abnormalities. *Iqsec3* mice were maintained by backcrossing onto the 129S5/SvEvBrd background; heterozygous males and females were fertile and used to set up intercrosses to generate homozygous and wildtype mice to study.

### *Baiap2* (IRSp53)

### 1 Mouse generation

#### 1.1 Mutation

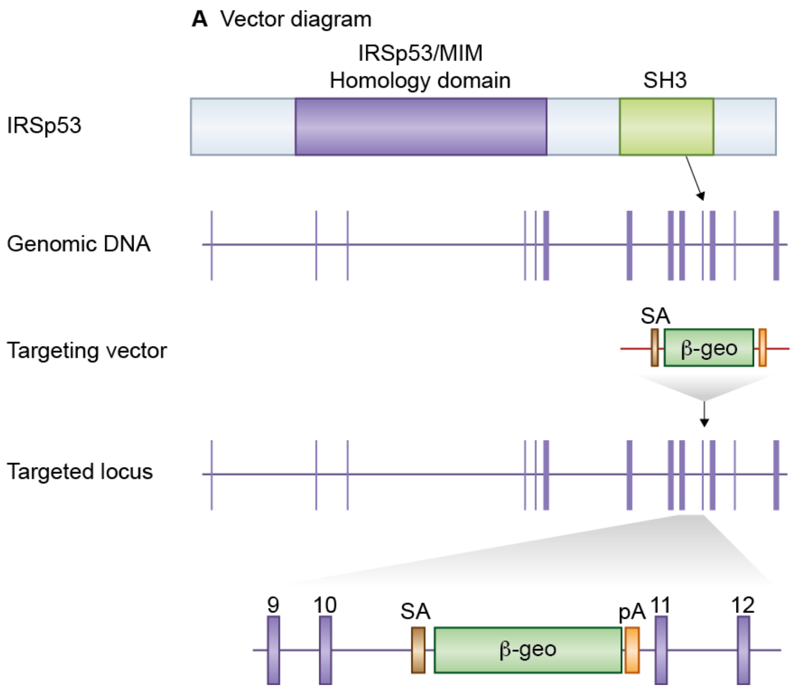


**Figure 1:** *Details of the mutation*

A mouse embryonic stem (ES) cell line (XN660, strain 129/Ola) with an insertional mutation in *Baiap2* was obtained from BayGenomics (baygenomics.ucsf.edu/). The insertional mutation in XN660 by the gene-trapping vector, pGT1lxf, that was designed to create an in-frame fusion between the 5' exons of the trapped gene and a reporter, ß-geo (a fusion of ß-galactosidase and neomycin phosphotransferase II) occurred in intron 10-11. Thus, the gene-trapped locus is predicted to yield a fusion transcript containing exons 1-10 of *Baiap2* and ß-geo. The ES cells were injected into C57BL/6 blastocysts to create chimeric mice, which were bred with 129S5 mice to generate heterozygous *Baiap2* mutant mice. Location of *Baiap2* gene trap. *Baiap2* is a 14 exon gene encoding the IRSp53 protein which contains IRSp53 homology and SH3 domains (top). The *Baiap2* gene trap is located in intron 10-11.

#### 1.2 Genotyping

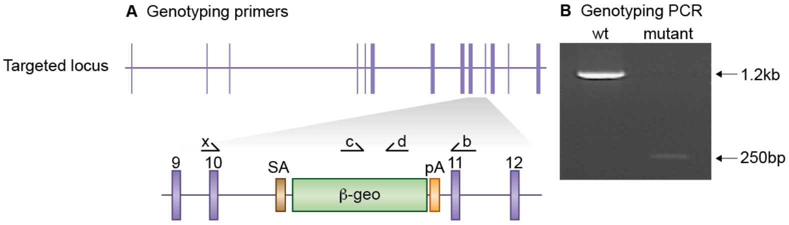


**Figure 2:** *Genotyping details*

Genomic DNA was isolated from ES cells or mouse tissues by Wizard SV 96 Genomic DNA purification system (Promega Cat A2371). Genotyping PCR consisted of 2 independent reactions. A 1.2kb product amplified from the wild-type (wt) allele using a forward primer X and a reverse primer B (5'- CTGTCACTTCCGTCACTGTC -3') in the wt sequence deleted by targeted mutation. The targeted allele is detected using forward primer C (5'- GCCATTGAACAAGATGGATTGCA- 3') with reverse primer D (5'- TTCCCGCTTCAGTGACAACGTC -3') that amplify a 250bp product within the ß-geo cassette. After enzymatic amplification for 35 cycles (45 seconds at 94 degC, 45 seconds at 55 degC, and 1 minute at 72 degC), the PCR products were size-fractionated on a 2% agarose gel in 1x Tris borate-EDTA buffer. Primers used for genotyping (x,b,c, d). PCR genotyping of gene trap IRSp53 mice using primers x,b to generate a 1.2kb wt product and primers c,d to amplify the mutant allele.

#### 1.3 Breeding

Birth of *IRSp53*$^{-/-}$ mice did not follow Mendelian ratios with 7% of offspring being homozygous knockouts. Genotypes of 3-week-old pups from *IRSp53*$^{+/-}$ intercrosses identified 150 wt, 170 *IRSp53*$^{+/-}$ and 63 *IRSp53*$^{-/-}$ progeny (Chi-squared p= $<$0.001). Male and female *IRSp53*$^{-/-}$ mice developed normally to adulthood, exhibited normal body size and no gross abnormalities. *IRSp53* mice were maintained by backcrossing onto the 129S5/SvEvBrd background; heterozygous males and females were fertile and used to set up intercrosses to generate homozygous and wildtype mice to study.

### *Kras* (K-Ras)

#### 1 Mouse generation

##### 1.1 Mutation

Conditional knockout of *Kras* in the brain was achieved by crossing LSL-Kras-G12D (K-ras$^{tm4Tyj}$), available from Jackson Laboratories and previously described by DA Tuveson et al. (Cancer Cell, 2004), and Camkcre4 (Tg(Camk2a-cre)1Gsc), previously described by T Mantamadiotis et al. (Nature Genetics, 2002).

##### 1.2 Genotyping

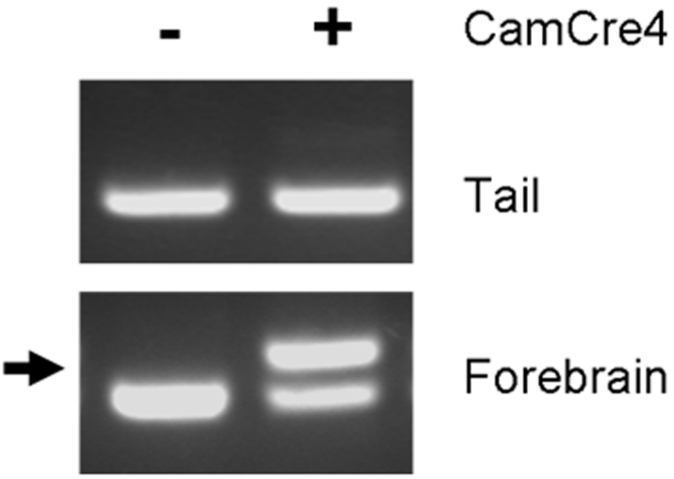


**Figure 1:** *Genotyping details*

Genomic DNA was extracted and prepared from tail and forebrain. Genotyping and analysis of recombination was done by PCR using HotStar DNA taq polymerase (Qiagen) according to manufacturers instructions. LSL-Kras-G12D mice were genotyped with primers; 5' CCTTTACAAGCGCACGCAGACTGTAGA 3' and 5' AGCTAGCCACCATGGCTTGAGTAAGTCTGCA 3'. Camkcre4 mice were genotyped with primers; 5'-TCGATGCAACGAGTGATGAG-3' and 5'-CCCAGAAATGCCAGATTACG -3'. Analysis of recombination was performed using *Kras* specific primers; 5'-GCAGCTAATGGCTCTCAAAGG-3' and 5'-TTGGCTCCAACACAGATGTTC-3'. Recombination of LSL-Kras-G12D occurs specifically in the forebrain of double transgenic mice. A 240bp band specific for recombination in targeted *Kras* gene (arrow) is seen in DNA extracted from forebrain but not DNA extracted from tail.

#### 1.3 Breeding

Birth of *K-Ras* mice from *K-Ras*$^{+/-}$-CAM-Cre$^{+/-}$ intercrosses was in line with expected ratios, with 19% of offspring being homozygous knockouts. Genotypes of 3-week-old pups from intercrosses identified 11 wt-wt, 9 wt-CAM-Cre$^{+/-}$, 15 *K-Ras*$^{+/-}$-wt, and 8 *K-Ras*$^{+/-}$-CAM-Cre$^{+/-}$ progeny (Chi-squared p= 0.445). Male and female double heterozygous mice showed no gross abnormalities. *K-Ras*$^{+/-}$ mice were maintained by backcrossing onto the C57BL/6-J background and CAM-Cre$^{+/-}$; mice were maintained by backcrossing onto the 129 background. A male K-Ras$^{+/-}$ mouse and female CAM-Cre$^{+/-}$ mice were used to set up intercrosses to generate homozygous and wildtype mice to study.

### *Cnksr2* (MAGUIN-1)

#### 1 Mouse generation

##### 1.1 Mutation

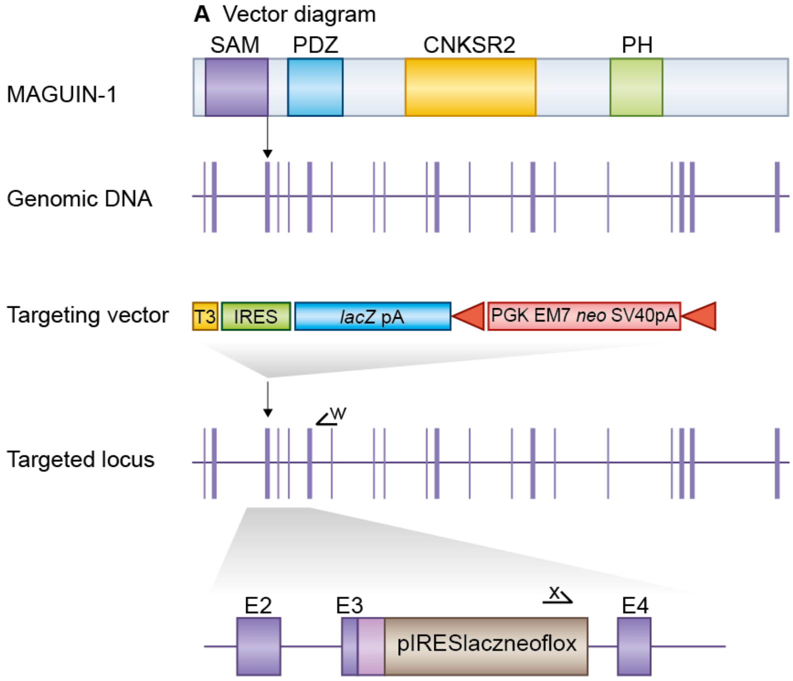


**Figure 1:** *Details of the mutation*

E14TG2a mouse embryonic stem (ES) cells were targeted with a vector containing 2.7kb and 6.6kb of flanking genomic DNA. This replaced 1.9kb of *Cnksr2* genomic DNA (X154431703 to X154433564; Ensemble Build 50) with IRES-lacZ-neo cassette. Correctly targeted ES cells were identified by long range PCR using Expand Long Template PCR system (Roche Cat 11681842001). The PCR contained primer X (5'- GAGC-TATTCCAGAAGTAGTGAG -3') and primer W (5'- GTAGGTCTTGATGACTGTTTGC -3') that correspond to sequence in the IRES-lacZ-neo cassette and sequence outside the 2.7kb flanking region respectively. The correctly targeted ES cells were injected into C57BL/6 blastocysts to create chimeric mice, which were bred with 129S5 mice to generate heterozygous *Cnksr2* mutant mice. Location of *Cnksr2* gene trap. *Cnksr2* is a 22 exon gene encoding the MAGUIN-1 protein which contains SAM, PDZ, Cnksr2 and PH domains (top). We replaced most of *Cnksr2* 3rd exon with a selection cassette in targeted mice and created a frameshift between exons 3 and 4. Primers used for targeted clone identification (w,x) are shown.

#### 1.2 Genotyping

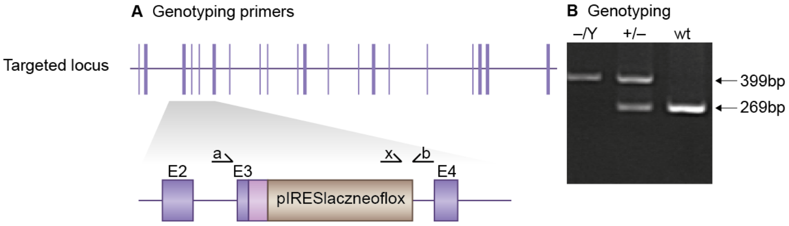


**Figure 2:** *Genotyping details*

Genomic DNA was isolated from ES cells by Wizard SV 96 Genomic DNA purification system (Promega Cat A2371). Genotyping PCR consisted of a 269bp product amplified from the wild-type (wt) allele using a forward primer A (CACTCTTCCTCCAACTCTTTC) in the wt sequence deleted by targeted mutation and a reverse primer B (CTCTTACTTGTCCCTGTGCAG) downstream of the cassette. A 399bp product was amplified from the targeted allele using primer B with forward primer X, within the selection cassette. After enzymatic amplification for 35 cycles (45 seconds at 94 degC, 45 seconds at 55 degC, and 1 minute at 72 degC), the PCR products were size-fractionated on a 2% agarose gel in 1x Tris borate-EDTA buffer. Primers used for genotyping (a,b, x). PCR genotyping of targeted MAGUIN-1 mice using a common reverse primer, b, and forward primers a and x to amplify the wt and mutant alleles respectively.

#### 1.3 Breeding

*Cnksr2* resides on the X chromosome: birth of *Cnksr2* $^{-/Y}$ mice followed Mendelian ratios with 24% of offspring being hemizygous knockouts but no female *Cnksr2*$^{-/-}$ mice were born. Genotypes of 3-week-old pups from Cnksr2 backcrosses identified 64 wt, 34 female *Cnksr2* $^{-/+}$ and 28 *Cnksr2* $^{-/Y}$ male progeny (Chi-squared p= 0.401). Male *Cnksr2* $^{-/Y}$ developed normally to adulthood, exhibited normal body size and no gross abnormalities. Backcrosses onto the 129S5/SvEvBrd background were used to maintain the colony and to generate hemizygous and wildtype mice to study.

### *Mpdz* (MUPP1)

### 1 Mouse generation

#### 1.1 Mutation

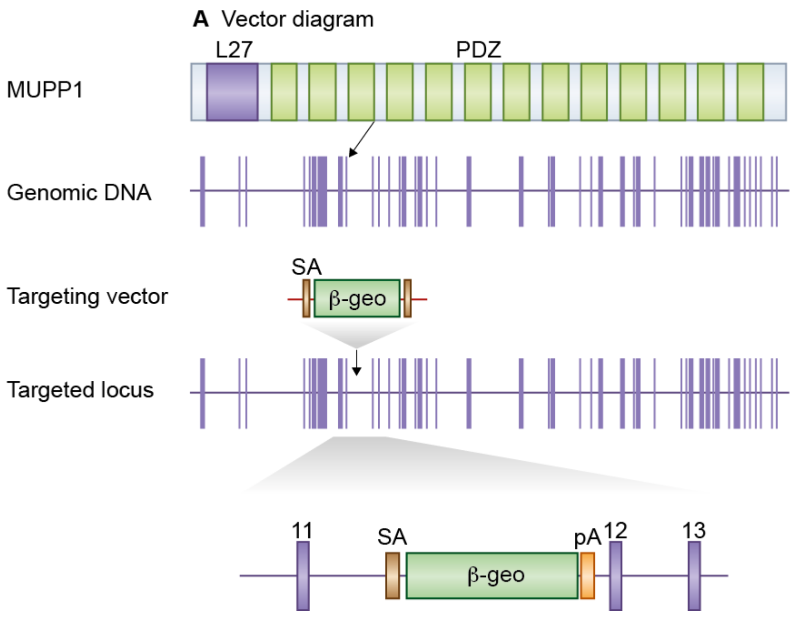


**Figure 1:** *Details of the mutation*

A mouse embryonic stem (ES) cell line (XG734, strain 129P2/OlaHsd) with an insertional mutation in *Mpdz* was obtained from BayGenomics (baygenomics.ucsf.edu/). The insertional mutation in XG734, by the gene-trapping vector, pGT1lxf, that was designed to create an in-frame fusion between the 5' exons of the trapped gene and a reporter, ß-geo (a fusion of beta-galactosidase and neomycin phosphotransferase II) occurred in intron 11-12. Thus, the gene-trapped locus is predicted to yield a fusion transcript containing exons 1-11 of *Mpdz* and ß-geo. The ES cells were injected into C57BL/6 blastocysts to create chimeric mice, which were bred with C57BL/6 mice to generate heterozygous *Mpdz* mutant mice. The severe hydrocephalic phenotype of homozygous neonates resulted in generating them in the heterozygous state. Location of *Mpdz* gene trap. *Mpdz* is a 47 exon gene encoding the protein MUPP1 which contains an L27 domain and 13 PDZ domains (top). The *Mpdz* gene trap is located in intron 11-12.

#### 1.2 Genotyping

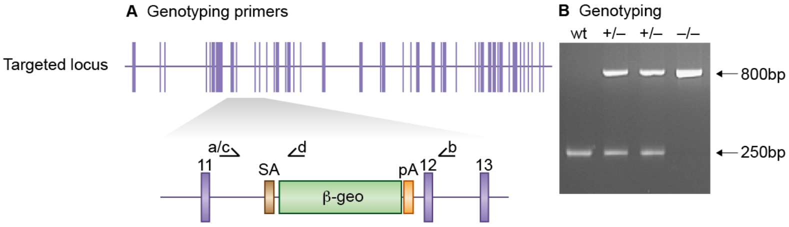


**Figure 2:** *Genotyping details*

Genomic DNA was isolated from ES cells or mouse tissues by Wizard SV 96 Genomic DNA purification system (Promega Cat A2371). Genotyping PCR consisted of 2 independent reactions. A 250bp product amplified from the wild-type (wt) allele using a forward primer A (5'- GTTCATACGGTTACTGTGGAG -3') upstream of the cassette and a reverse primer B (5'- CATAATGAAATCCTGAGCCTG -3') in the wt sequence deleted by targeted mutation. A 800bp product was amplified from the targeted allele using forward primer C (5'- GTGCGTCTGACACTGATGAG-3') upstream of the insertion point with reverse primer D (5'- CTCTTCACATCCATGCTGAG -3'), within the ß-geo cassette. After enzymatic amplification for 35 cycles (45 seconds at 94 degC, 45 seconds at 55 degC, and 1 minute at 72 degC), the PCR products were size-fractionated on a 2% agarose gel in 1x Tris borate-EDTA buffer. Primers used for genotyping (a,b,c, d). PCR genotyping of gene trap *Mpdz* mice using primers a and b to amplify wild type alleles and primers c and d to amplify mutant alleles.

#### 1.3 Breeding

Birth of *Mpdz*$^{-/-}$ mice did not follow Mendelian ratios with 6% of offspring being homozygous knockouts; these progeny displayed increased perinatal mortality rates from birth to 4 weeks but tissue was collected for post mortem genotyping. Genotypes from *Mpdz*$^{+/-}$ intercrosses identified 75 wt, 152 *Mpdz*$^{+/-}$ and 14 *Mpdz*$^{-/-}$ progeny (Chi-squared p= <0.001). Backcrosses onto the C57BL/6 background were used to maintain the *Mpdz* colony and to generate heterozygous and wildtype mice to study. Genomic composition of the majority of mice used in experiments was approximately 25% 129P2/OlaHsd and 75% C57BL/6.

### *Sorbs2* (nArgBP2)

#### 1 Mouse generation

##### 1.1 Mutation

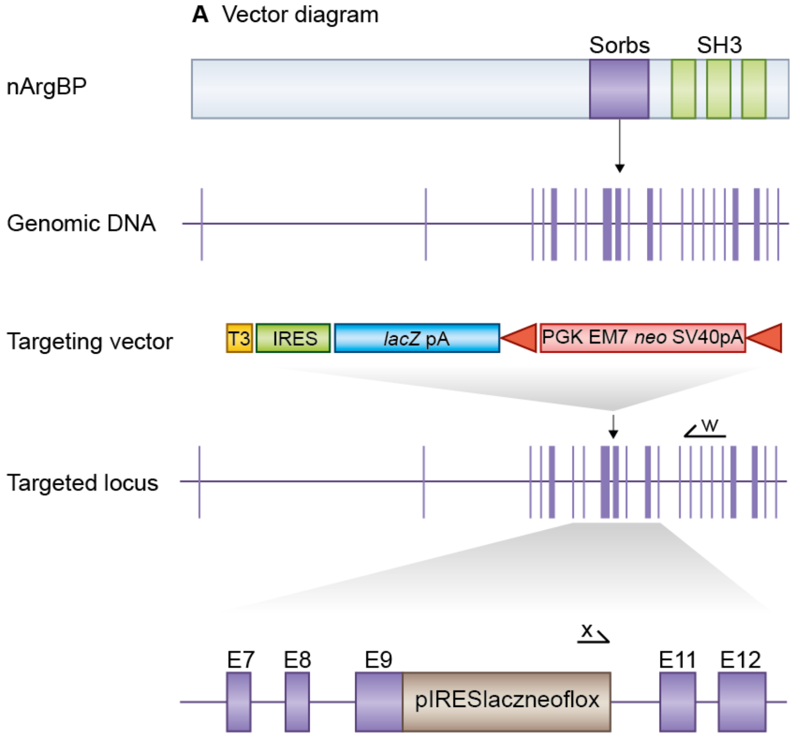


**Figure 1:** *Details of the mutation*

E14TG2a mouse embryonic stem (ES) cells were targeted with a vector containing 6.7kb and 3.3kb of flanking genomic DNA. This replaced 940bp of *Sorbs2* genomic DNA (X46855222 to X46856162; Ensemble Build 55) with IRES-lacZ-neo cassette. Correctly targeted ES cells were identified by long range PCR using Expand Long Template PCR system (Roche Cat 11681842001). The PCR contained primer X (5'-GAGCTATTCCAGAAGTAGTGAG-3') and primer W (5'- CTACCTGTACAATACTGTGTC -3') that correspond to sequence in the IRES-lacZ-neo cassette and sequence outside the 3.3kb flanking region respectively. The correctly targeted ES cells were injected into C57BL/6 blastocysts to create chimeric mice, which were bred with 129S5 mice to generate heterozygous *Sorbs2* mutant mice. Location of *Sorbs2* gene trap. *Sorbs2*, a 26 exon gene, encodes the protein ArgBP2 which contains a Sorbs domain and 3 SH3 domains (top). We replaced *Sorbs2* exons 9-10 with a selection cassette in targeted mice and created a frameshift between exons 8 and 11. Primers used for targeted clone identification (w,x) are shown.

#### 1.2 Genotyping

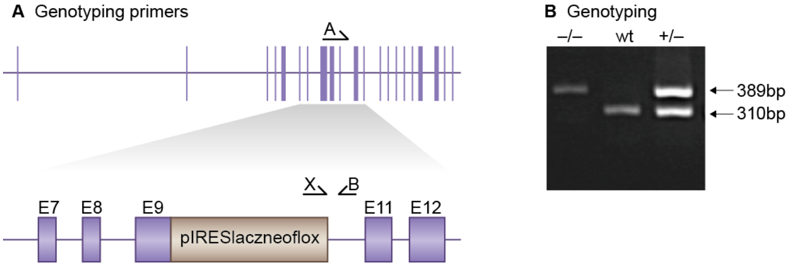


**Figure 2:** *Genotyping details*

Genomic DNA was isolated from ES cells by Wizard SV 96 Genomic DNA purification system (Promega Cat A2371). Genotyping PCR consisted of a 310bp product amplified from the wild-type (wt) allele using a forward primer A (GATGACCAATCACACTCCAC) in the wt sequence deleted by targeted mutation and a reverse primer B (GTCTTATGCAACCTTAGTCAC) downstream of the cassette. A 389bp product was amplified from the targeted allele using primer B with forward primer X, within the selection cassette. After enzymatic amplification for 35 cycles (45 seconds at 94 degC, 45 seconds at 55 degC, and 1 minute at 72 degC), the PCR products were size-fractionated on a 2% agarose gel in 1x Tris borate-EDTA buffer. Primers used for genotyping (A,B, X). PCR genotyping of targeted nArgBP2 mice using a common reverse primer B and forward primers A and X to amplify the wt and mutant alleles respectively.

#### 1.3 Breeding

Birth of *Sorbs2*$^{-/-}$ mice followed Mendelian ratios with 20% of offspring being homozygous knockouts. Genotypes of 3-week-old pups from *Sorbs2*$^{+/-}$ intercrosses identified 75 wt, 145 *Sorbs2*$^{+/-}$ and 55 *Sorbs2*$^{-/-}$ progeny (Chi-squared p= 0.155). Male and female *Sorbs2*$^{-/-}$ mice developed normally to adulthood, exhibited normal body size and no gross abnormalities but displayed an increased mortality rate between 14-21 weeks. *Sorbs2* mice were maintained by backcrossing onto the 129S5/SvEvBrd background; heterozygous males and females were fertile and used to set up intercrosses to generate homozygous and wildtype mice to study.

# *Nf1* (NF1)

#### 1 Mouse generation

##### 1.1 Mutation

Generation of *Nf1* mutant mice described in PMID:7920653, Jacks *et al* (1994)

##### 1.2 Genotyping

*Nf1* genotyping described in PMID:7920653, Jacks *et al* (1994)

##### 1.3 Breeding

No *NF1*$^{-/-}$ mice were produced from *NF1*$^{+/-}$ intercrosses. Male and female *NF1* $^{+/-}$ mice developed normally to adulthood, were fertile, exhibited normal body size and no gross abnormalities. Genotypes of 3-week-old pups from *NF1*$^{+/-}$ intercrosses identified 14 wt and 16 *NF1*$^{+/-}$ progeny (Chi-squared p= $<$0.001). Back-crosses onto the 129S5/SvEvBrd background were used to maintain the colony and to generate heterozygous and wildtype mice to study.

### *Rasal2* (nGAP)

#### 1 Mouse generation

##### 1.1 Mutation

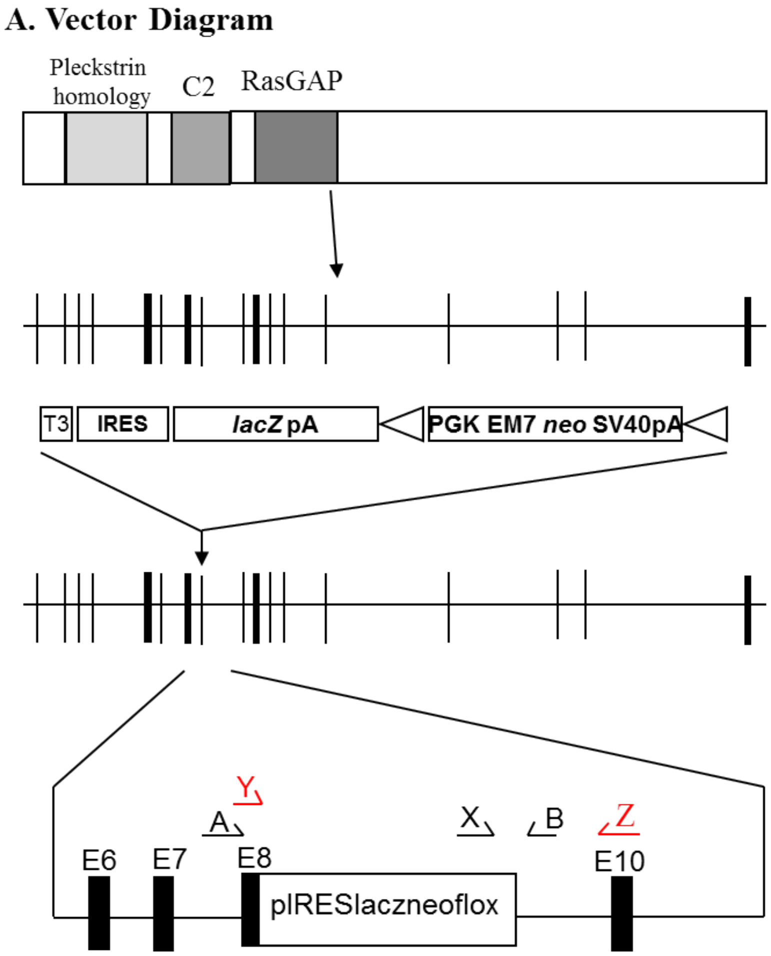


**Figure 1:** *Details of the mutation*

E14TG2a mouse embryonic stem (ES) cells were targeted with a vector containing 2.3kb and 2.8kb of flanking genomic DNA. This replaced 1.9kb of *Rasal2* genomic DNA (X157173891 to X157175814; Ensemble Build 75) with an IRES-lacZ-neo cassette. Genomic DNA was isolated from ES cells by Wizard SV 96 Genomic DNA purification system (Promega Cat A2371). Correctly targeted ES cells were identified by long range PCR using Expand Long Template PCR system (Roche Cat 11681842001). The correctly targeted ES cells were injected into C57BL/6 blastocysts to create chimeric mice, which were bred with 129S5 mice to generate heterozygous *Rasal2* mutant mice. Generation of *Rasal2* targeted mice *Rasal2* is an 18 exon gene that encodes a protein containing Pleckstrin homology, C2 and RasGAP domains (top). We replaced *Rasal2* exons 8-9 with a selection cassette in targeted mice and created a frameshift between exons 8 and 10. Primers used for genotyping(A,B, X) and RT-PCR (Y,Z).

#### 1.2 Genotyping

Genotyping PCR consisted of a 2.7kb product amplified from the wild-type (wt) allele using a forward primer A (GACGAGCTGAAAATGTTCTCC) in the wt sequence upstream of the cassette and a reverse primer B (CAGCACTTTCTTCAAGGAACTC) downstream of the cassette. A 640bp product was amplified from the targeted allele using primer B with forward primer X (CATGTCTGGATCGATATCCC), within the selection cassette. After enzymatic amplification for 35 cycles (45 seconds at 94 degC, 45 seconds at 55 degC, and 1 minute at 72 degC), the PCR products were size-fractionated on a 1% agarose gel in 1x Tris borate-EDTA buffer (Image not shown).

#### 1.3 Breeding

Birth of *Rasal2*$^{-/-}$ mice followed Mendelian ratios with 17% of offspring being homozygous knockouts. Genotypes of 3-week old pups from *Rasal2*$^{+/-}$ intercrosses identified 86 wt, 121 *Rasal2*$^{+/-}$ and 42 *Rasal2*$^{-/-}$ progeny (Chi-squared $p<0.001$). Male and female *Rasal2*$^{-/-}$ mice developed normally to adulthood, exhibited normal body size and no gross abnormalities. *Rasal2* mice were maintained by backcrossing onto the 129S5/SvEvBrd background; heterozygous males and females were fertile and used to set up intercrosses to generate homozygous and wildtype mice to study.

### *Grin2a* (NR2A-2BCTR)

#### 1 Mouse generation

##### 1.1 Mutation

Generation of *Grin2a* mutant mice described in PMCID:PMC3979286, *Ryan et al* (2013)

##### 1.2 Genotyping

*Grin2*a genotyping described in PMCID:PMC3979286, *Ryan et al* (2013)

##### 1.3 Breeding

Birth of mutant mice followed approximately Mendelian ratios with 21% of offspring being homozygous knockouts. Genotypes of 3-week-old pups from heterozygous intercrosses identified 89 wt, 154 heterozygous, and 63 homozygous progeny (Chi-squared p=0.11). Male and female homozygous mice developed normally to adulthood, exhibited normal body size and no gross abnormalities. Backcrosses onto the C57BL/6J background were used to maintain the colony and to generate homozygous and wildtype mice to study.

### *Grin2a* (NR2A-dC)

#### 1 Mouse generation

##### 1.1 Mutation

Generation of *Grin2a* mutant mice described in PMID:9458051, Sprengel *et al* (1998)

##### 1.2 Genotyping

*Grin2a* genotyping described in PMID:9458051, Sprengel *et al* (1998)

##### 1.3 Breeding

Birth of NR2A-dC$^{-/-}$ mice followed Mendelian ratios with 21% of offspring being homozygous knockouts. Genotypes of 3-week-old pups from NR2A-dC$^{+/-}$ intercrosses identified 47 wt, 98 NR2A-dC$^{+/-}$ and 39 NR2A-dC$^{-/-}$ progeny (Chi-squared p= 0.478). Male and female NR2A-dC$^{-/-}$ mice developed normally to adulthood, exhibited normal body size and no gross abnormalities NR2A-dC mice were maintained by backcrossing onto the C57BL/6J background; heterozygous males and females were fertile and used to set up intercrosses to generate homozygous and wildtype mice to study.

### *Grin2b* (NR2B-2ACTR)

#### 1 Mouse generation

##### 1.1 Mutation

Generation of *Grin2b* mutant mice described in PMCID:PMC3979286, Ryan *et al* (2013)

##### 1.2 Genotyping

*Grin2b* genotyping described in PMCID:PMC3979286, Ryan *et al* (2013)

##### 1.3 Breeding

Birth of mutant mice followed approximately Mendelian ratios with 19% of offspring being homozygous knock-outs. Genotypes of 3-week-old pups from heterozygous intercrosses identified 79 wt, 136 heterozygous, and 52 homozygous progeny (Chi-squared p=0.062). Male and female homozygous mice developed normally to adult-hood, exhibited normal body size and no gross abnormalities. Backcrosses onto the C57BL/6 background were used to maintain the colony and to generate homozygous and wildtype mice to study.

### *Grin2b* (NR2B-CK2)

#### 1 Mouse generation

##### 1.1 Mutation

A detailed description of these mutant mice will be published elsewhere.

##### 1.2 Genotyping

A detailed description of these mutant mice will be published elsewhere.

##### 1.3 Breeding

A detailed description of these mutant mice will be published elsewhere.

### *Grin2b* (NR2B-dC)

#### 1 Mouse generation

##### 1.1 Mutation

Generation of *Grin2b* mutant mice described in PMID:9458051, Sprengel *et al* (1998)

##### 1.2 Genotyping

*Grin2b* genotyping described in PMID:9458051, Sprengel *et al* (1998)

##### 1.3 Breeding

No NR2B-dC$^{-/-}$ mice were produced from NR2B-dC$^{+/-}$ intercrosses. Male and female NR2B-dC$^{+/-}$ mice developed normally to adulthood, were fertile, exhibited normal body size and no gross abnormalities. Genotypes of 3-week-old pups from NR2B-dC$^{+/-}$ intercrosses identified 9 wt and 18 NR2B-dC$^{+/-}$ progeny (Chi-squared p= $<$0.001). Backcrosses onto the C57BL/6J background were used to maintain the colony and to generate heterozygous and wildtype mice to study.

### *Grin2b* (NR2B-dV)

#### 1 Mouse generation

##### 1.1 Mutation

A detailed description of these mutant mice will be published elsewhere.

##### 1.2 Genotyping

A detailed description of these mutant mice will be published elsewhere.

##### 1.3 Breeding

A detailed description of these mutant mice will be published elsewhere.

### *Rapgef2* (nRapGEP)

#### 1 Mouse generation

##### 1.1 Mutation

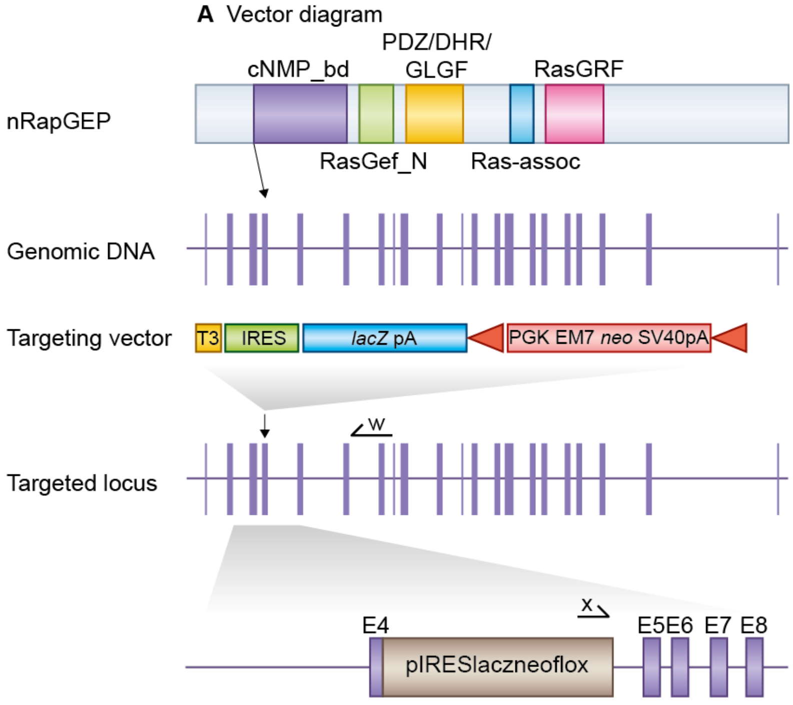


**Figure 1:** *Details of the mutation*

E14TG2a mouse embryonic stem (ES) cells were targeted with a vector containing 6.6Kb and 3.6Kb of flanking genomic DNA. This replaced 934bp of *Rapgef2* genomic DNA (3 78,903,114 to 3 78,904,048; Ensemble Build 50) with IRES-lacZ-neo cassette. Correctly targeted ES cells were identified by long range PCR using Expand Long Template PCR system (Roche Cat 11681842001). The PCR contained primer X (5'-CTATGAGTGGATCAGTGATACG -3') and primer W (5'- CAAGATCAGTTGTCATCGGAG -3') that correspond to sequence outside the 3.6kb flanking region sequence, and in the IRES-lacZ-neo cassette respectively. The correctly targeted ES cells were injected into C57BL/6 blastocysts to create chimeric mice, which were bred with 129S5 mice to generate heterozygous *Rapgef2* mutant mice. Location of *Rapgef2* gene trap. *Rapgef2* is a 24 exon gene encoding the protein nRapGEP which contains cNMP-bd, RasGef N, PDZ/DHR/GLGF, Ras-assoc and RasGRF domains (top). We replaced most of *Rapgef2* exon 4 with a selection cassette in targeted mice and created a frameshift between exons 4 and 5. Primers used for targeted clone identification (w,x) are shown.

#### 1.2 Genotyping

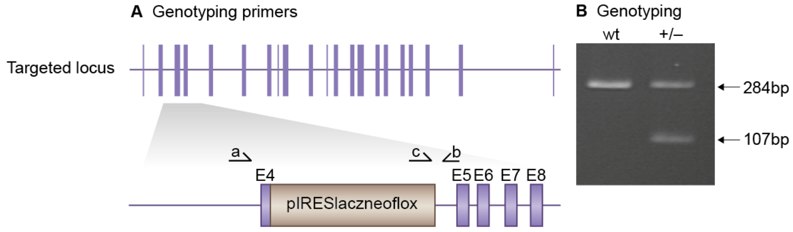


**Figure 2:** *Genotyping details*

Genomic DNA was isolated from ES cells by Wizard SV 96 Genomic DNA purification system (Promega Cat A2371). Genotyping PCR consisted of a 284bp product amplified from the wild-type (wt) allele using a forward primer A (CTAGTTTAGGACACGTCTAG) in the wt sequence deleted by targeted mutation and a reverse primer B (GTGTTCGTTACTGAGGGTC) downstream of the cassette. A 107bp product was amplified from the targeted allele using reverse primer B with forward primer C (CACTGCATTCTAGTTGTGG), within the selection cassette. After enzymatic amplification for 35 cycles (45 seconds at 94 degC, 45 seconds at 55 degC, and 1 minute at 72 degC), the PCR products were size-fractionated on a 2% agarose gel in 1x Tris borate-EDTA buffer. Primers used for genotyping (a,b, c). PCR genotyping of targeted *Rapgef2* mice using a common reverse primer, b, and forward primers a and c to amplify the wt and mutant alleles respectively.

#### 1.3 Breeding

No *Rapgef2*$^{-/-}$ mice were produced from *Rapgef2*$^{+/-}$ intercrosses. Male and female *Rapgef2* $^{+/-}$ mice developed normally to adulthood, were fertile, exhibited normal body size and no gross abnormalities. Genotypes of 3-week-old pups from *Rapgef2*$^{+/-}$ intercrosses identified 39 wt and 107 *Rapgef2*$^{+/-}$ progeny (Chi-squared p= $<$0.001). Backcrosses onto the 129S5/SvEvBrd background were used to maintain the colony and to generate heterozygous and wildtype mice to study.

### *Nsf* (NSF)

#### 1 Mouse generation

##### 1.1 Mutation

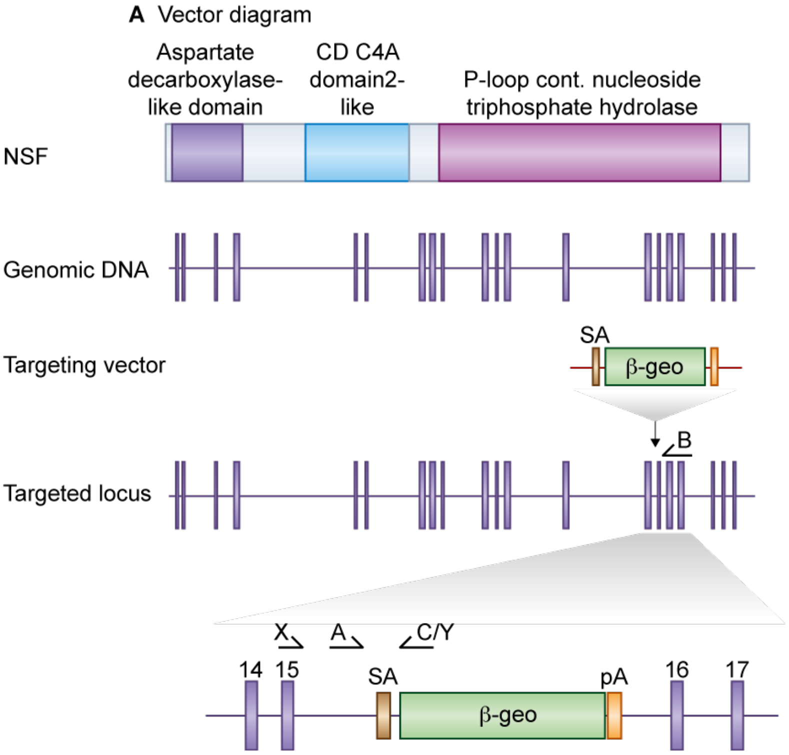


**Figure 1:** *Details of the mutation*

A mouse embryonic stem (ES) cell line (AK0189, strain 129/Ola) with an insertional mutation in *Nsf* was obtained from Sanger Institute Gene Trap Resource (SIGTR - sanger.ac.uk/PostGenomics/genetrap/). The insertional mutation in AK0189, by the gene-trapping vector, pGT0lxr, that was designed to create an in-frame fusion between the 5' exons of the trapped gene and a reporter, ß-geo (a fusion of ß-galactosidase and neomycin phosphotransferase II), occurred in intron 15-16. Thus, the gene-trapped locus is predicted to yield a fusion transcript containing exons 1-15 of *Nsf* and ß-geo. The ES cells were injected into C57BL/6 blastocysts to create chimeric mice, which were bred with 129S5 mice to generate heterozygous *Nsf*-mutant mice. Location of *Nsf* gene trap. *Nsf* is a 21 exon gene encoding a protein containing an aspartate decarboxylase, CD C4A and a P-loop containing nucleoside triphosphate hydrolase domain (top). The *Nsf* gene trap is located in intron 15-16. Primers used for genotyping (A,B, C) and RT-PCR (X,Y) are shown.

#### 1.2 Genotyping

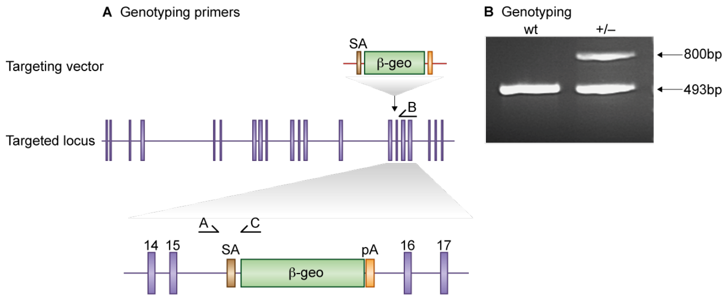


**Figure 2:** *Genotyping details*

Genomic DNA was isolated from ES cells or mouse tissues by Wizard SV 96 Genomic DNA purification system (Promega Cat A2371). Genotyping PCR consisted of a 493 bp product amplified from the wild-type (wt) allele using a forward primer A (5'- CACATCGACAGCTTCTTCTG -3') upstream of the cassette and a reverse primer B (5'- TCAATGTCATCCACGACCAC -3'). A 800bp product was amplified from the targeted allele using primer A with reverse primer C (5'- CAGGCTTCACTGAGTCTCTG -3') within the ß-geo cassette. After enzymatic amplification for 35 cycles (45 seconds at 94 degC, 45 seconds at 55 degC, and 2 minutes at 72 degC), the PCR products were size-fractionated on a 2% agarose gel in 1x Tris borate-EDTA buffer. Primers used for genotyping (A,B, C). PCR genotyping of gene trap NSF mice using a common forward primer, A, and reverse primers B and C to amplify the wt and mutant alleles respectively.

#### 1.3 Breeding

No *Nsf*$^{-/-}$ mice were produced from *Nsf*$^{+/-}$ intercrosses. Male and female *Nsf*$^{+/-}$ mice developed normally to adulthood, were fertile, exhibited normal body size and no gross abnormalities. Genotypes of 3-week-old pups from *Nsf*$^{+/-}$ intercrosses identified 21 wt and 57 *Nsf*$^{+/-}$ progeny (Chi-squared $p = <0.001$). Back-crosses onto the 129S5/SvEvBrd background were used to maintain the colony and to generate heterozygous and wildtype mice to study.

### *Agap2* (PIKE)

#### 1 Mouse generation

##### 1.1 Mutation

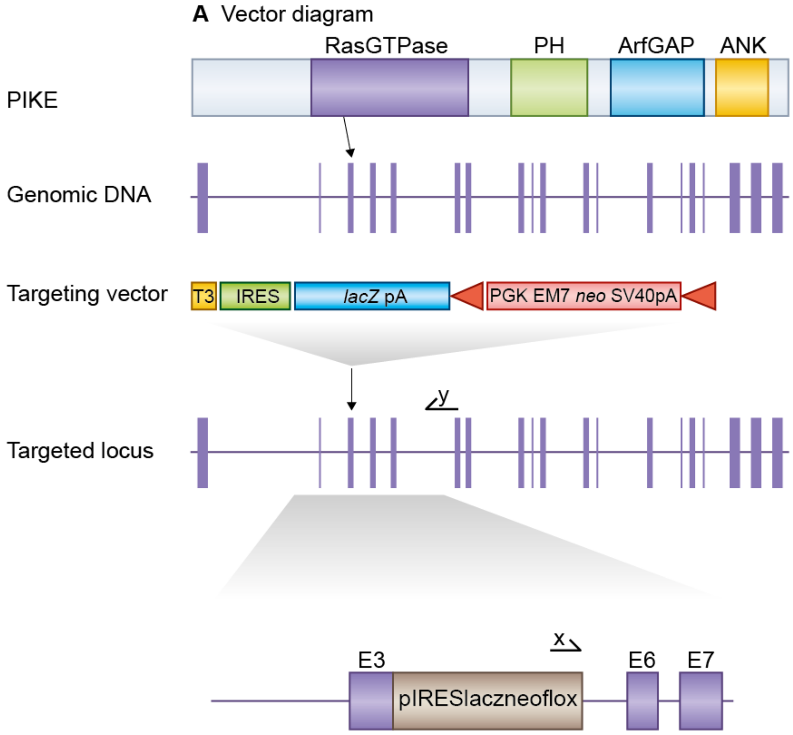


**Figure 1:** *Details of the mutation*

E14TG2a mouse embryonic stem (ES) cells were targeted with a vector containing 6.8kb and 3.2kb of flanking genomic DNA. This replaced 945bp of *Agap2* genomic DNA (X126520053 to X126520998; Ensemble Build 50) with IRES-lacZ-neo cassette. Correctly targeted ES cells were identified by long range PCR using Expand Long Template PCR system (Roche Cat 11681842001). The PCR contained primer X (5'-GAGCTATTCCAGAAGTAGTGAG-3') and primer Y (5'- GTTTCCCAGTGTGACTGTCTC -3') that correspond to sequence in the IRES-lac-Zneo cassette and sequence outside the 3.2kb flanking region respectively. The correctly targeted ES cells were injected into C57BL/6 blastocysts to create chimeric mice, which were bred with 129S5 mice to generate heterozygous *Agap2* mutant mice. Location of *Agap2* gene trap. *Agap2* is a 19 exon gene which encodes Ras GTPase, PH, ArfGAP and ANK domains (top). We replaced *Agap2* exons 4 and 5 with a selection cassette in targeted mice and created a frameshift between exons 3 and 6. Primers used for targeted clone identification (x,y) are shown.

#### 1.2 Genotyping

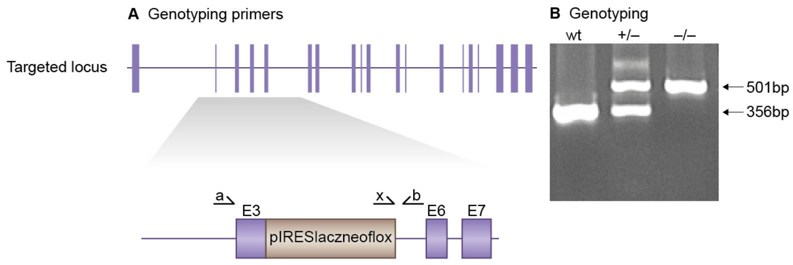


**Figure 2:** *Genotyping details*

Genomic DNA was isolated from ES cells by Wizard SV 96 Genomic DNA purification system (Promega Cat A2371). Genotyping PCR consisted of a 356bp product amplified from the wild-type (wt) allele using a forward primer A (GTTCACATTGCCATTTCTGAC) in the wt sequence deleted by targeted mutation and a reverse primer B (GGAGAACAATGAGAGGTGAAC) downstream of the cassette. A 501bp product was amplified from the targeted allele using primer B with forward primer X, within the selection cassette. After enzymatic amplification for 35 cycles (45 seconds at 94 degC, 45 seconds at 55 degC, and 1 minute at 72 degC), the PCR products were size-fractionated on a 2% agarose gel in 1x Tris borate-EDTA buffer. Primers used for genotyping (a,b, x). PCR genotyping of targeted *Agap2* mice using a common reverse primer, b, and forward primers a and x to amplify the wildtype and mutant alleles respectively.

#### 1.3 Breeding

Birth of *Agap2*$^{-/-}$ mice followed Mendelian ratios with 20% of offspring being homozygous knockouts. Genotypes of 3-week-old pups from *Agap2*$^{+/-}$ intercrosses identified 70 wt, 170 *Agap2*$^{+/-}$ and 63 *Agap2*$^{-/-}$ progeny (Chi-squared p= 0.089). Male and female *Agap2*$^{-/-}$ mice developed normally to adulthood, exhibited normal body size and no gross abnormalities. *Agap2* mice were maintained by backcrossing onto the 129S5/SvEvBrd background; heterozygous males and females were fertile and used to set up intercrosses to generate homozygous and wildtype mice to study.

### *Prr7* (Prr7)

#### 1 Mouse generation

##### 1.1 Mutation

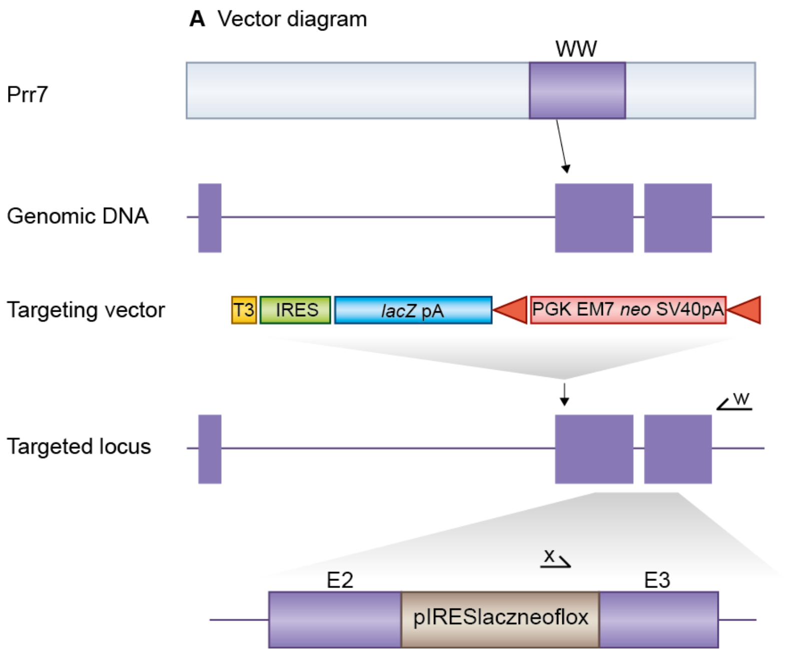


**Figure 1:** *Details of the mutation*

E14TG2a mouse embryonic stem (ES) cells were targeted with a vector containing 5.4kb and 2.9kb of flanking genomic DNA. This replaced 700bp of *Prr7* genomic DNA (X55573423 to X55574123; Ensemble Build 55) with IRES-lacZ-neo cassette. Correctly targeted ES cells were identified by long range PCR using Expand Long Template PCR system (Roche Cat 11681842001). The PCR contained primer X (5'-GAGCTATTCCAGAAGTAGTGAG-3') and primer W (5'- CTCACAACCACCTATAATGAG -3') that correspond to sequence in the IRES-lacZ-neo cassette and sequence outside the 2.9kb flanking region respectively. The correctly targeted ES cells were injected into C57BL/6 blastocysts to create chimeric mice, which were bred with 129S5 mice to generate heterozygous *Prr7* mutant mice. Location of *Prr7* gene trap. *Prr7* is a 3 exon gene, encoding the Prr7 protein which contains a WW binding domain (top). We replaced most of *Prr7* exons 2,3 with a selection cassette in targeted mice and created a frameshift between exons 2 and 3. Primers used for targeted clone identification (x,w) are shown.

#### 1.2 Genotyping

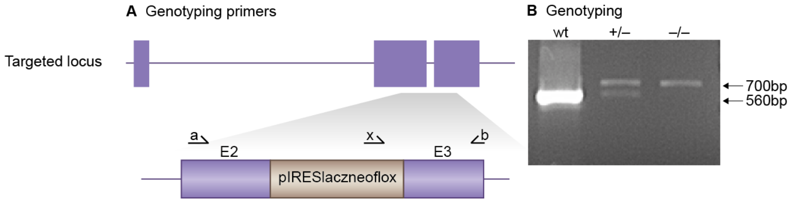


**Figure 2:** *Genotyping details*

Genomic DNA was isolated from ES cells by Wizard SV 96 Genomic DNA purification system (Promega Cat A2371). Genotyping PCR consisted of a 560bp product amplified from the wild-type (wt) allele using a forward primer A (GGAATCGGACATGTCTAAG) in the wt sequence deleted by targeted mutation and a reverse primer B (GTACCAAAGCAGATCACACAC) downstream of the cassette. A 700bp product was amplified from the targeted allele using primer B with forward primer X, within the selection cassette. After enzymatic amplification for 35 cycles (45 seconds at 94 degC, 45 seconds at 55 degC, and 1 minute at 72 degC), the PCR products were size-fractionated on a 2% agarose gel in 1x Tris borate-EDTA buffer. Primers used for genotyping (a,b, x). PCR genotyping of targeted *Prr7* mice using a common reverse primer, b, and forward primers a and x to amplify the wt and mutant alleles respectively.

#### 1.3 Breeding

Birth of *Prr7*$^{-/-}$ mice followed Mendelian ratios with 27% of offspring being homozygous knockouts. Genotypes of 3-week-old pups from *Prr7*$^{+/-}$ intercrosses identified 62 wt, 99 *Prr7*$^{+/-}$ and 61 *Prr7*$^{-/-}$ progeny (Chi-squared p= 0.272). Male and female *Prr7*$^{-/-}$ mice developed normally to adulthood, exhibited normal body size and no gross abnormalities. *Prr7* mice were maintained by backcrossing onto the 129S5/SvEvBrd background; heterozygous males and females were fertile and used to set up intercrosses to generate homozygous and wildtype mice to study.

### *Dlg2* (PSD-93)

#### 1 Mouse generation

##### 1.1 Mutation

Generation of PSD-93 mutant mice described in PMID:11312293, McGee *et al* (2001)

##### 1.2 Genotyping

*Dlg2* genotyping described in PMID:11312293, McGee *et al* (2001)

##### 1.3 Breeding

Birth of *Dlg2*$^{-/-}$ mice followed Mendelian ratios with 20% of offspring being homozygous knockouts. Genotypes of 3-week-old pups from *Dlg2*$^{+/-}$ intercrosses identified 50 wt, 106 *Dlg2*$^{+/-}$ and 40 *Dlg2*$^{-/-}$ progeny (Chi-squared p= 0.312). Male and female Dlg2$^{-/-}$ mice developed normally to adulthood, exhibited normal body size and no gross abnormalities. *Dlg2* mice were maintained by backcrossing onto the C57BL/6J background; heterozygous males and females were fertile and used to set up intercrosses to generate homozygous and wildtype mice to study.

### *Dlg4* (PSD-95-GK)

### 1 Mouse generation

#### 1.1 Mutation

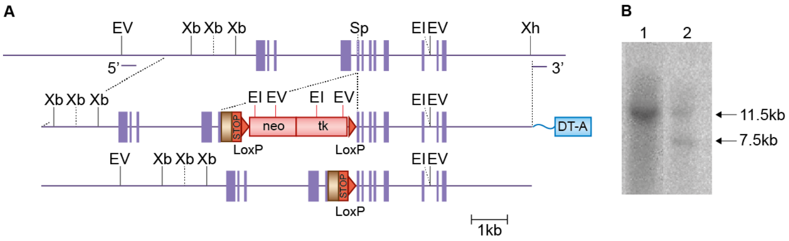


**Figure 1:** *Details of the mutation*

E14TG2a mouse embryonic stem (ES) cells were targeted with a vector containing 5.2kb and 5kb of flanking genomic DNA. This was inserted in frame into exon 13 of *Dlg4* with neomycin/tk cassette. Correctly targeted ES cells were identified by long range PCR and southern blot. For southern blot, genomic DNA was cut with EcoRV, XbaI + HindIII or Xba alone for null mutants. The correctly targeted ES cells were injected into C57BL/6 blastocysts to create chimeric mice, which were further bred with C57BL/6J mice to generate heterozygous *Dlg4* mutant mice. Targeted mutation of the PSD-95 gene. a. PSD-95 genomic DNA with restriction enzyme sites (EI, EcoRI; EV, EcoRV; Sp, SpeI; Xb, XbaI; Xh, XhoI.); boxes, exons. With integrated targeting vector (middle) and targeted locus (bottom). tk, thymidine kinase gene; neo, neomycin resistance gene; HA, haemaglutinin epitope tag; STOP, stop codon; arrowheads, loxP sites; DT-A, Diptheria toxin A based vector. b. Genomic southern blot of XbaI digested ES cell DNA following Cre excision of selection cassette probed with 3' flanking probe. Lane 1, wild type (wt) DNA showing 11.5 kb band; lane 2, wt and 7.5 kb mutant band as predicted.

#### 1.2 Genotyping

Genotyping of mice was carried out by PCR using the following primers: A PSD-95 DNA fragment of 2400bp was amplified using a forward primer 5'-CCGACTGCTCACTAGTATTTTCTCCC-3' with reverse primer 5'-CCGTAGAGGTGGCTGTTGTA-3'. Wild type and mutant alleles were distinguished by digesting with XbaI giving rise to a 1400bp mutant fragment.

#### 1.3 Breeding

Birth of *Dlg4(GK)*$^{-/-}$ mice followed approximately Mendelian ratios with 21% of offspring being homozygous knockouts. Genotypes of 3-week-old pups from *Dlg4(GK)*$^{+/-}$ intercrosses identified 31 wt, 79 Dlg4(GK)$^{+/-}$ and 29 *Dlg4(GK)*$^{-/-}$ progeny (Chi-squared p= 0.265). Male and female *Dlg4(GK)*$^{-/-}$ mice developed normally to adulthood, exhibited normal body size and no gross abnormalities. *Dlg4(GK)* mice were maintained by backcrossing onto the C57BL/6J background; heterozygous males and females were fertile and used to set up intercrosses to generate homozygous

### *Dlg4* (PSD-95-SH3)

#### 1 Mouse generation

##### 1.1 Mutation

Generation of *Dlg4* mutant mice described in PMID:20467438, Arbuckle *et al* (2010)

##### 1.2 Genotyping

*Dlg4* genotyping described in PMID:20467438, Arbuckle *et al* (2010)

##### 1.3 Breeding

Birth of PSD-95-SH3$^{-/-}$ departed only slightly from Mendelian ratios with 17% of offspring being homozygous knockouts. Genotypes of 3-week-old pups from PSD-95-SH3$^{+/-}$ intercrosses identified 48 wt, 72 PSD-95-SH3 $^{+/-}$, and 25 PSD-95-SH3$^{-/-}$ progeny (Chi-squared p=0.026). Male and female PSD-95-SH3$^{-/-}$ mice exhibited no gross abnormalities. Backcrosses onto the C57BL/6J background were used to maintain the colony and to generate homozygous and wildtype mice to study.

### *Rap1gap* (Rap1gap)

#### 1 Mouse generation

##### 1.1 Mutation

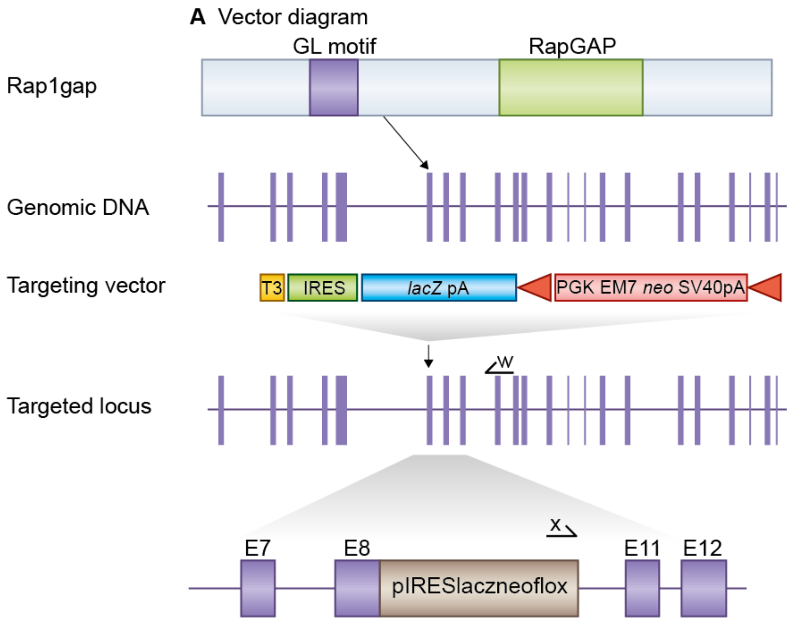


**Figure 1:** *Details of the mutation*

E14TG2a mouse embryonic stem (ES) cells were targeted with a vector containing 6.9kb and 3.2kb of flanking genomic DNA. This replaced 908bp of *Rap1gap* genomic DNA (X137271961 to X137272599; Ensemble Build 55) with IRES-lacZ-neo cassette. Correctly targeted ES cells were identified by long range PCR using Expand Long Template PCR system (Roche Cat 11681842001). The PCR contained primer X (5'-GAGCTATTCCAGAAGTAGTGAG-3') and primer W (5'- GTACCTTTACAGATAGCTGTG -3') that correspond to sequence in the IRES-lacZ-neo cassette and sequence outside the 3.2kb flanking region respectively. The correctly targeted ES cells were injected into C57BL/6 blastocysts to create chimeric mice, which were bred with 129S5 mice to generate heterozygous *Rap1gap* mutant mice. Location of *Rap1gap* gene trap. *Rap1gap* is a 25 exon gene which encodes a GoLoco Motif and RapGAP domain(top). We replaced exons 9,10 with a selection cassette in targeted mice and created a frameshift between exons 8 and 11. Primers used for targeted clone identification (W,X) are shown.

#### 1.2 Genotyping

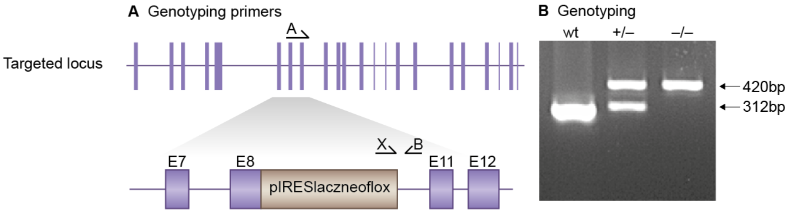


**Figure 2:** *Genotyping details*

Genomic DNA was isolated from ES cells by Wizard SV 96 Genomic DNA purification system (Promega Cat A2371). Genotyping PCR consisted of a 312bp product amplified from the wild-type (wt) allele using a forward primer A (GTATGTGAGGATGTCAATGTG) in the wt sequence deleted by targeted mutation and a reverse primer B (CTGTACCTCATGTGCTTTAAG) downstream of the cassette. A 420bp product was amplified from the targeted allele using primer B with forward primer X, within the selection cassette. After enzymatic amplification for 35 cycles (45 seconds at 94 degC, 45 seconds at 55 degC, and 1 minute at 72 degC), the PCR products were size-fractionated on a 2% agarose gel in 1x Tris borate-EDTA buffer. Primers used for genotyping (A,B, X). PCR genotyping of targeted Rap1gap mice using a common reverse primer, B, and forward primers A and X to amplify the wt and mutant alleles respectively.

#### 1.3 Breeding

Birth of *Rap1gap*-/- mice followed Mendelian ratios with 25% of offspring being homozygous knockouts. Genotypes of 3-week-old pups from *Rap1gap*+/- intercrosses identified 39 wt, 76 *Rap1gap*+/- and 38 *Rap1gap*-/- progeny (Chi-squared p= 0.879). Male and female *Rap1gap*-/- mice developed normally to adulthood, exhibited normal body size and no gross abnormalities Rap1gap mice were maintained by backcrossing onto the 129S5/SvEvBrd background; heterozygous males and females were fertile and used to set up intercrosses to generate homozygous and wildtype mice to study.

### *Rassf1* (Rassf1)

#### 1 Mouse generation

##### 1.1 Mutation

Generation of *Rassf1* mutant mice described in PMID:16135822, van der Weyden et al (2005)

##### 1.2 Genotyping

*Rassf1* genotyping described in PMID:16135822, van der Weyden et al (2005)

##### 1.3 Breeding

Cohorts of *Rassf1a* mice were provided by the Wellcome Trust Sanger Institute (colony code MAMP). Birth of *Rassf1a*-/- mice is reported to follow Mendelian ratios (van der Weyden, L, et al, Mol Cell Biol 25:8356-8367 (2005)). Male and female *Rassf1a*-/- mice are also reported to develop normally to adulthood, exhibited normal body size and no gross abnormalities. Heterozygous males and females were fertile and used to set up intercrosses to generate homozygous and wildtype mice to study.

### *Dlg1* (SAP97)

#### 1 Mouse generation

##### 1.1 Mutation

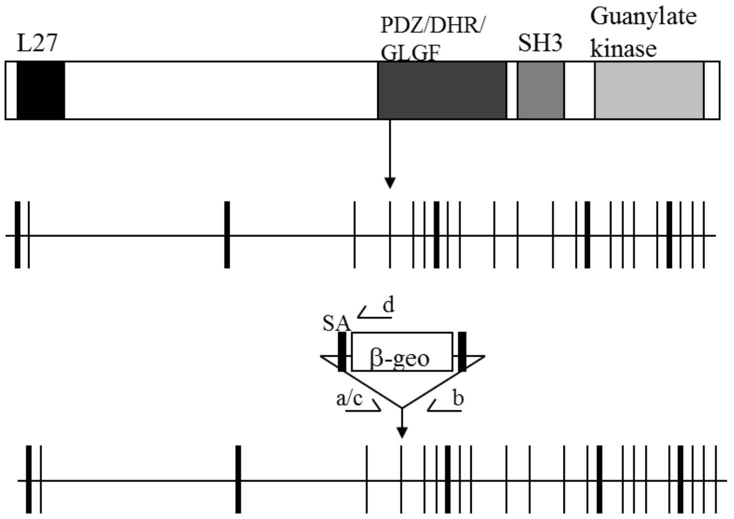


**Figure 1:** *Details of the mutation*

A mouse embryonic stem (ES) cell line (AJ0497, strain 129/Ola) with an insertional mutation in *Dlg1* was obtained from Sanger Institute Gene Trap Resource (SIGTR - sanger.ac.uk/PostGenomics/genetrap/). The insertional mutation in AJ0497, by the gene-trapping vector, pGT0lxr, that was designed to create an in-frame fusion between the 5' exons of the trapped gene and a reporter, ß-geo (a fusion of ß-galactosidase and neomycin phosphotransferase II), occurred in intron 5-6. Thus, the gene-trapped locus is predicted to yield a fusion transcript containing exons 1-5 of *Dlg1* and ß-geo. The ES cells were injected into C57BL/6 blastocysts to create chimeric mice, which were bred with 129S5 mice to generate heterozygous*Dlg1* mutant mice. Primers used for genotyping (a, b, c and d). PCR genotyping of gene trap SAP97 mice using forward primers, a and c, and reverse primers, b and d, to amplify the wt and mutant alleles respectively.

#### 1.2 Genotyping

Genomic DNA was isolated from ES cells or mouse tissues by Wizard SV 96 DNA purification system (Promega Cat A2371). Genotyping PCR consisted of 2 independent reactions. A 421bp product amplified from the wild-type (wt) allele using a forward primer a (5'- CAGTCCTCAAATCCCATCGG -3') upstream of the cassette and a reverse primer b (5'- CACTGTCTGAAGGAACACTG -3') in the wt sequence deleted by targeted mutation. The targeted allele is detected using forward primer c (5'- CTGTCCTGGAACTCACTCTG - 3') with reverse primer d (5'- CTCAAAGTCAGGGTCACAAGG -3') within the gene trap cassette and amplifies a 500bp product. After enzymatic amplification for 35 cycles (45 seconds at 94 degC, 45 seconds at 55 degC, and 1 minute at 72 degC), the PCR products were size-fractionated on a 2% agarose gel in 1x Tris borate-EDTA buffer (image not shown).

#### 1.3 Breeding

No *Dlg1*$^{-/-}$ mice were produced from *Dlg1*$^{+/-}$ intercrosses. Male and female *Dlg1*$^{+/-}$ mice developed normally to adulthood, were fertile, exhibited normal body size and no gross abnormalities. Genotypes of 3-week-old pups from *Dlg1*$^{+/-}$ intercrosses identified 70 wt and 99 Dlg1$^{+/-}$ progeny (Chi-squared $p = <0.001$). Backcrosses onto the 129S5/SvEvBrd background were used to maintain the colony and to generate heterozygous and wildtype mice to study.

### *Dlg3* (SAP102)

#### 1 Mouse generation

##### 1.1 Mutation

Generation of *Dlg3* mutant mice described in PMCID:PMC2851144, Cuthbert *et al* (2007)

##### 1.2 Genotyping

*Dlg3* genotyping described in PMCID:PMC2851144, Cuthbert *et al* (2007)

##### 1.3 Breeding

*Dlg3* resides on the X chromosome. Birth of SAP102-/Y mice followed approximately Mendelian ratios with 25% of offspring being hemizygous knockouts. Birth of homozygous knockouts also followed Mendelian ratios with 19% of offspring being homozygous (female) knockouts. Genotypes of 3-week-old pups from heterozygous intercrosses identified 34 male wt, 21 female wt, 32 SAP102+/-, 17 SAP102-/Y, and 13 homozygous (female) progeny (Chi-squared p=0.25). Male SAP102-/Y and female homozygous mice developed normally to adulthood, exhibited normal body size and no gross abnormalities. Backcrosses onto the C57BL/6J background were used to maintain the colony and to generate homozygous, hemizygous and wildtype mice to study.

### *Shank2* (SHANK2)

#### 1 Mouse generation

##### 1.1 Mutation

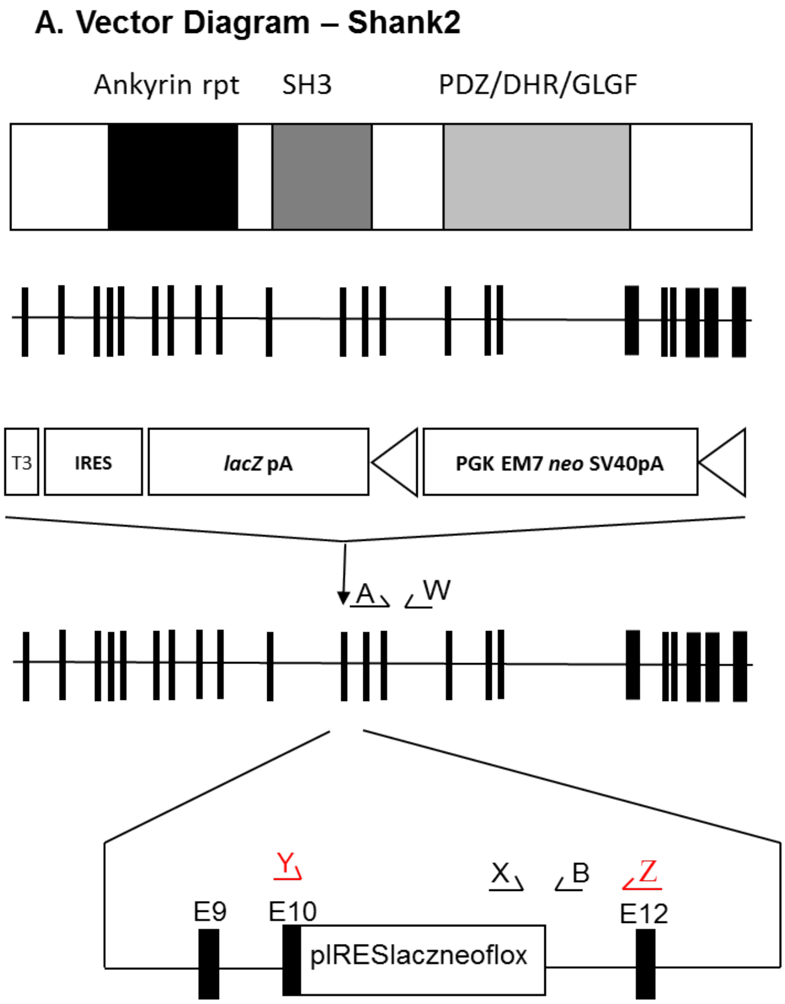


**Figure 1:** *Details of the mutation*

E14TG2a mouse embryonic stem (ES) cells were targeted with a vector containing 6.2kb and 2.9kb of flanking genomic DNA. This replaced 1.1kb of Shank2 genomic DNA (X144179906 to X144180964; Ensemble Build 75) with IRES-lacZ-neo cassette. Genomic DNA was isolated from ES cells by Wizard SV 96 Genomic DNA purification system (Promega Cat A2371). Correctly targeted ES cells were identified by long range PCR using Expand Long Template PCR system (Roche Cat 11681842001). The PCR contained forward primer X (5'-GAGCTATTCCAGAAGTAGTGAG-3') and reverse primer W (5'- CACATAGTGCAGTGTCAACTG -3') that correspond to sequence in the IRES-lacZ-neo cassette and sequence outside the 2.9kb flanking region respectively. The correctly targeted ES cells were injected into C57BL/6 blastocysts to create chimeric mice, which were bred with 129S5 mice to generate heterozygous *Shank2* mutant mice.

Generation of *Shank2* targeted mice. *Shank2* is a 23 exon gene encoding a protein which contains Ankyrin repeat, SH3 and PDZ domains (top). We replaced most of Shank2 exon 11 with a selection cassette in targeted mice and created a frameshift between exons 11 and 12. Primers used for genotyping(A,B, X), RT-PCR (Y,Z) and targeted clone identification (W,X) are shown.

#### 1.2 Genotyping

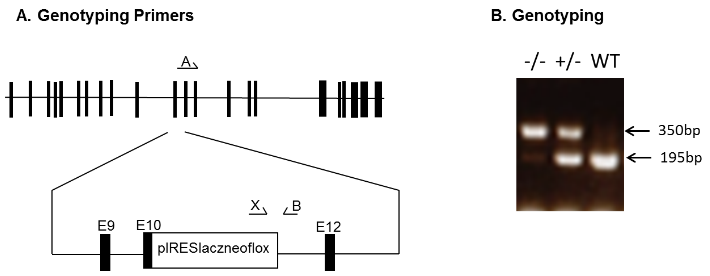


**Figure 2:** *Genotyping details*

Genomic DNA was isolated from ES cells or mouse tissues by Wizard SV 96 Genomic DNA purification system (Promega Cat A2371). Genotyping PCR consisted of a 195bp product amplified from the wild-type (wt) allele using a forward primer A (CTGCCTTCCTAAGTTTCTTAG) in the wt sequence upstream of the cassette and a reverse primer B (GTGAAGACATTGCTATGGCAG) downstream of the cassette. A 350bp product was amplified from the targeted allele using primer B with forward primer X, within the selection cassette. After enzymatic amplification for 35 cycles (45 seconds at 94 degC, 45 seconds at 55 degC, and 1 minute at 72 degC), the PCR products were size-fractionated on a 1% agarose gel in 1x Tris borate-EDTA buffer. Primers used for genotyping (A,B, X).PCR genotyping of targeted SHANK2 mice using a common reverse primer, B, and forward primers A and X to amplify the wt and mutant alleles respectively.

#### 1.3 Breeding

Birth of *Shank2*$^{-/-}$ mice followed Mendelian ratios with 22% of offspring being homozygous knockouts. Genotypes of 3-week-old pups from *Shank2*$^{+/-}$ intercrosses identified 73 wt, 147 *Shank2*$^{+/-}$ and 62 *Shank2*$^{-/-}$ progeny (Chi-squared p= 0.504). Male and female Shank2$^{-/-}$ mice showed no gross abnormalities. *Shank2* mice were maintained by backcrossing onto the 129S5/SvEvBrd background; heterozygous males and females were fertile and used to set up intercrosses to generate homozygous and wildtype mice to study.

### *Sipa1l1* (SPAR)

### 1 Mouse generation

#### 1.1 Mutation

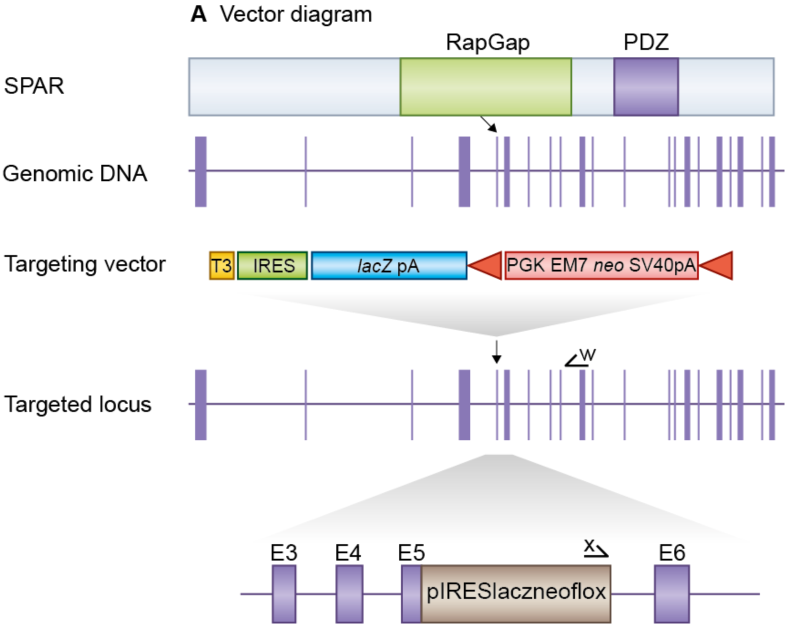


**Figure 1:** *Details of the mutation*

E14TG2a mouse embryonic stem (ES) cells were targeted with a vector containing 7.2kb and 3kb of flanking genomic DNA. This replaced 1.2kb of *Sipa1l1* genomic DNA (X83442041 to X83443195; Ensemble Build 50) with IRES-lacZ-neo cassette. Correctly targeted ES cells were identified by long range PCR using Expand Long Template PCR system (Roche Cat 11681842001). The PCR contained primer X (5'- CAGC-TAAGTGGAATGAGGACTC -3') and primer W (5'- CACTACTTCTGGAATAGCTCAG -3') that correspond to sequence in the IRES-lacZ-neo cassette and sequence outside the 3kb flanking region respectively. The correctly targeted ES cells were injected into C57BL/6 blastocysts to create chimeric mice, which were bred with 129S5 mice to generate heterozygous *Sipa1l1* mutant mice. Location of *Sipa1l1* gene trap. *Sipa1l1* is a 24 exon gene, encoding the SPAR protein which contains a RapGAP and PDZ domain(top). We replaced most of exon 5 with a selection cassette in targeted mice and created a frameshift between exons 5 and 6. Primers used for targeted clone identification (w,x) are shown.

#### 1.2 Genotyping

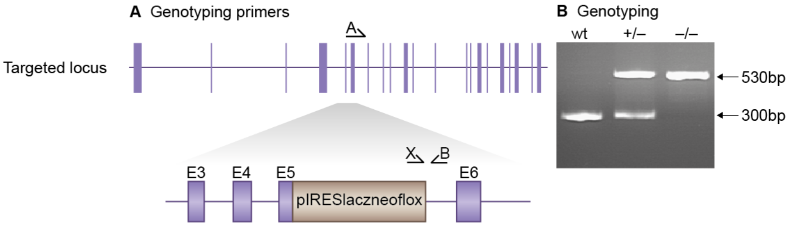


**Figure 2:** *Genotyping details*

Genomic DNA was isolated from ES cells by Wizard SV 96 Genomic DNA purification system (Promega Cat A2371).Genotyping PCR consisted of a 303bp product amplified from the wild-type (wt) allele using a forward primer A (CCACTATGATGTCCAGAG) in the wt sequence deleted by targeted mutation and a reverse primer B (CTTGGACAGGCTGATCTTC) downstream of the cassette. A 532bp product was amplified from the targeted allele using primer B with forward primer X (CTTCTTGACGAGTTCTTCTG), within the selection cassette. After enzymatic amplification for 35 cycles (45 seconds at 94 degC, 45 seconds at 55 degC, and 1 minute at 72 degC), the PCR products were size-fractionated on a 2% agarose gel in 1x Tris borate-EDTA buffer. Primers used for genotyping (A,B, X). PCR genotyping of targeted *Sipa1l1* mice using a common reverse primer, B, and forward primers A and X to amplify the wt and mutant alleles respectively.

#### 1.3 Breeding

Birth of *Sipa1l1*$^{-/-}$ mice did not follow Mendelian ratios with 21% of offspring being homozygous knockouts. Genotypes of 3-week-old pups from *Sipa1l1*$^{+/-}$ intercrosses identified 71 wt, 104 *Sipa1l1*$^{+/-}$ and 47 *Sipa1l1*$^{-/-}$ progeny (Chi-squared p= 0.048). Male and female *Sipa1l1*$^{-/-}$ mice developed normally to adulthood, exhibited normal body size and no gross abnormalities. *Sipa1l1* mice were maintained by backcrossing onto the 129S5/SvEvBrd background; heterozygous males and females were fertile and used to set up intercrosses to generate homozygous and wildtype mice to study.

### *Magi2* (S-SCAM)

### 1 Mouse generation

#### 1.1 Mutation

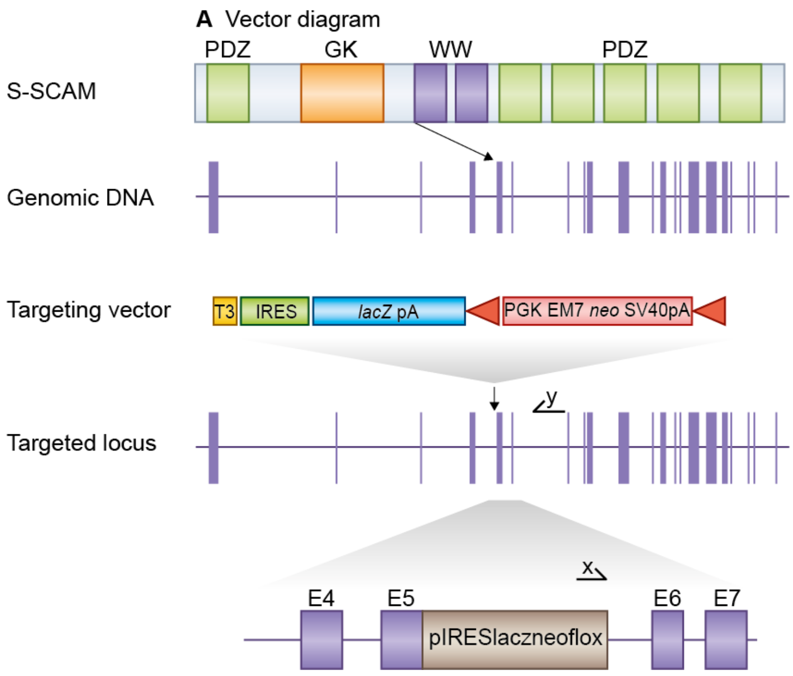


**Figure 1:** *Details of the mutation*

E14TG2a mouse embryonic stem (ES) cells were targeted with a vector containing 5.9kb and 2.9kb of flanking genomic DNA. This replaced 2.3 kb of *Magi2* genomic DNA (X19,721,359 to X19723711; Ensemble Build 50) with IRES-lacZ-neo cassette. Correctly targeted ES cells were identified by long range PCR using Expand Long Template PCR system (Roche Cat 11681842001). The PCR contained primer X (5'-GAGCTATTCCAGAAGTAGTGAG-3') and primer Y (5'- CAAGATCAGTTGTCATCGGAG -3') that correspond to sequence in the IRES-lacZ-neo cassette and sequence outside the 2.9kb flanking region respectively. The correctly targeted ES cells were injected into C57BL/6 blastocysts to create chimeric mice, which were bred with 129S5 mice to generate heterozygous *Magi2* mutant mice. Those F1 heterozygous mice had been backcrossed with 129S5 mice for 1-2 times before being used for intercrossing.Location of *Magi2* gene trap. *Magi2*, a 23 exon gene, encodes the protein S-SCAM which contains GK, WW and multiple PDZ domains (top). We replaced most of *Magi2* exon 5 with a selection cassette in targeted mice and created a frameshift between exons 5 and 6. Primers used for targeted clone identification (X,Y) are shown.

#### 1.2 Genotyping

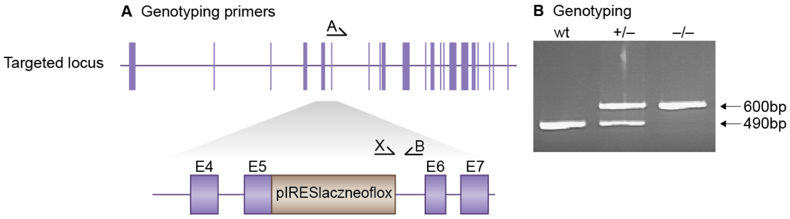


**Figure 2:** *Genotyping details*

Genomic DNA was isolated from ES cells by Wizard SV 96 Genomic DNA purification system (Promega Cat A2371). Genotyping PCR consisted of a 490bp product amplified from the wild-type (wt) allele using a forward primer A (GTCTGACTTTGTGCTTATCAG) in the wt sequence deleted by targeted mutation and a reverse primer B (CTCAGCTAGTACTCTAGAAG) downstream of the cassette. A 600bp product was amplified from the targeted allele using primer B with forward primer X, within the selection cassette. After enzymatic amplification for 35 cycles (45 seconds at 94 degC, 45 seconds at 55 degC, and 1 minute at 72 degC), the PCR products were size-fractionated on a 2% agarose gel in 1x Tris borate-EDTA buffer. Primers used for genotyping (A,B, X). PCR genotyping of targeted S-SCAM mice using a common reverse primer, B, and forward primers A and X to amplify the wt and mutant alleles respectively.

#### 1.3 Breeding

Birth of *Magi2*$^{-/-}$ mice did not follow Mendelian ratios with 5% of offspring being homozygous knockouts; these progeny did not survive beyond birth but tissue was collected for post mortem genotyping. Genotypes from *Magi2*$^{+/-}$ intercrosses identified 49 wt, 89 *Magi2*$^{+/-}$ and 7 *Magi2*$^{-/-}$ progeny (Chi-squared p= $<$0.001). Male and female *Magi2*$^{+/-}$ mice developed normally to adulthood, exhibited normal body size and no gross abnormalities. Backcrosses onto the 129S5/SvEvBrd background were used to maintain the *Magi2* colony and to generate heterozygous and wildtype mice to study.

### *Cadm1* (SynCam)

#### 1 Mouse generation

##### 1.1 Mutation

Generation of *Cadm1* mutant mice described in PMID:16611999, van der Weyden *et al* (2006)

##### 1.2 Genotyping

*Cadm1* genotyping described in PMID:16611999, van der Weyden *et al* (2006)

##### 1.3 Breeding

Cohorts of *Cadm1* mice were provided by the Wellcome Trust Sanger Institute (colony code MAMQ). Birth of *Cadm1*-/- mice is reported to follow Mendelian ratios (van der Weyden, L, et al, Mol Cell Biol 26:3595-3609 (2006)). Male and female *Cadm1*-/- mice are also reported to develop normally to adulthood, exhibited normal body size and no gross abnormalities. *Cadm1* mice are maintained by backcrossing onto a mixed 129S5/SvEvBrd and C57BL/6 background; heterozygous males and females were fertile and used to set up intercrosses to generate homozygous and wildtype mice to study. Male *Cadm1*-/- mice are reported to be infertile. Cadm1 is also known as Tslc1.

### *Syngap1* (SynGAP)

#### 1 Mouse generation

##### 1.1 Mutation

Generation of *Syngap1* mutant mice described in PMID:12427827, Komiyama *et al* (2002)

##### 1.2 Genotyping

*Syngap1* genotyping described in PMID:12427827, Komiyama *et al* (2002)

##### 1.3 Breeding

No *Syngap1*$^{-/-}$ mice were produced from *Syngap1*+/- intercrosses. Male and female *Syngap1*$^{+/-}$ mice developed normally to adulthood, were fertile, exhibited normal body size and no gross abnormalities. Genotypes of 3-week-old pups from *Syngap1*$^{+/-}$ intercrosses identified 7 wt and 16 *Syngap1*$^{+/-}$ progeny (Chi-squared p= 0.020). Backcrosses onto the C57BL/6 background were used to maintain the colony and to generate heterozygous and wildtype mice to study.

### *Tanc1* (TANC1)

#### 1 Mouse generation

##### 1.1 Mutation

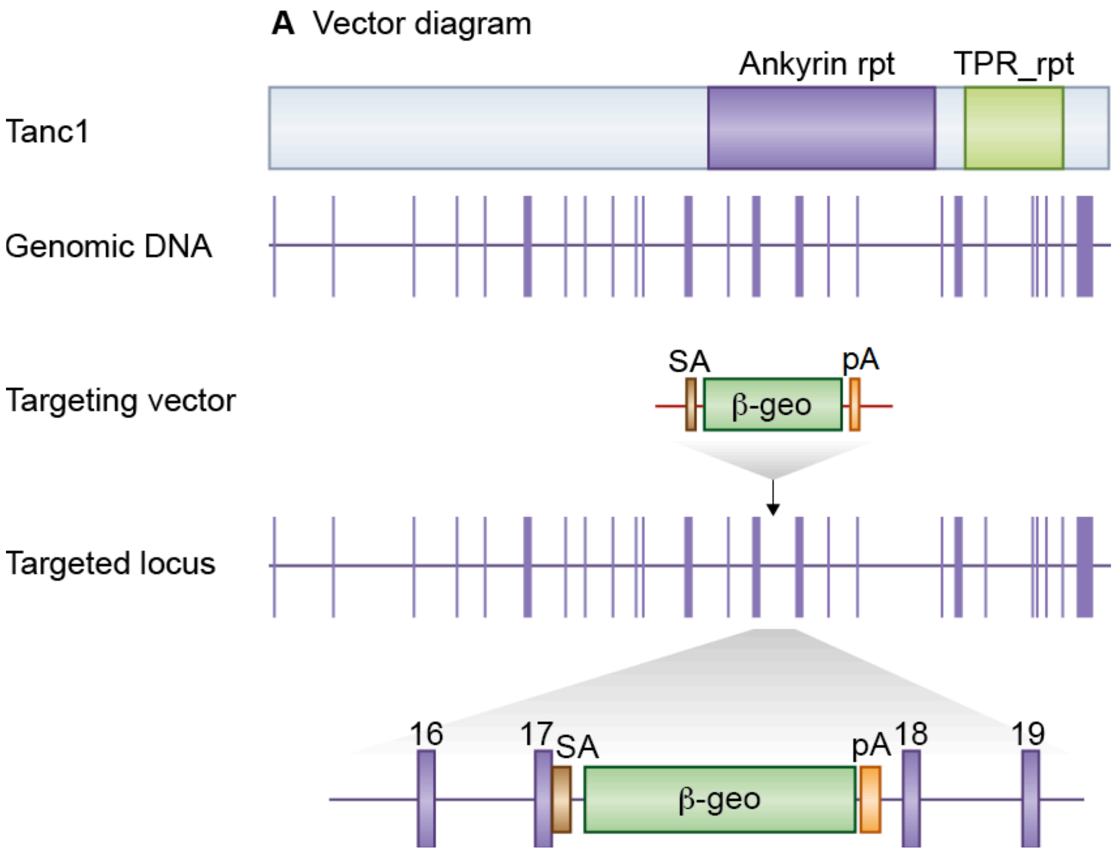


**Figure 1:** *Details of the mutation*

A mouse embryonic stem (ES) cell line (CE0101, strain 129/Ola) with an insertional mutation in *Tanc1* was obtained from Sanger Institute Gene Trap Resource (SIGTR - sanger.ac.uk/PostGenomics/genetrap/). The insertional mutation in CE0101 by the gene-trapping vector, pGT1lxf, that was designed to create an in-frame fusion between the 5' exons of the trapped gene and a reporter, β-geo (a fusion of β-galactosidase and neomycin phosphotransferase II) occurred within exon 17. Thus, the gene-trapped locus is predicted to yield a fusion transcript containing exons 1-17 of *Tanc1* and β-geo. The ES cells were injected into C57BL/6 blastocysts to create chimeric mice, which were bred with 129S5 mice to generate heterozygous (+/-) Tanc1-mutant mice. Those F1 heterozygous mice had been backcrossed with 129S5 mice for 1-2 times before being used for intercrossing.

**Location of *Tanc1* gene trap mice** *Tanc1* is a 27 exon gene encoding the Tanc1 protein and contain ankyrin repeat and TPR repeat domains (top). The *Tanc1* gene trap is located in exon 17.

#### 1.2 Genotyping

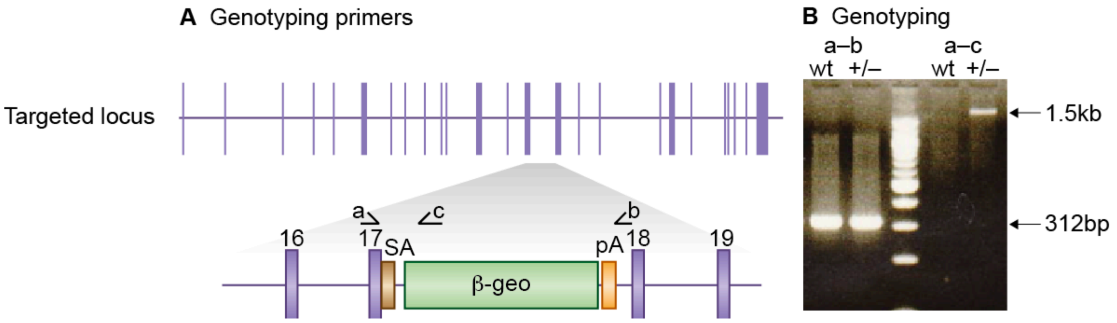


#### Figure 2: *Genotyping details*

Genomic DNA was isolated from ES cells or mouse tissues by Wizard SV 96 Genomic DNA purification system (Promega Cat A2371). Genotyping PCR consisted of a 312bp product amplified from the wild-type (wt) allele using a forward primer A (5'- CATGAAGAAGTTGTCACTCTC -3') and a reverse primer B (5'- CAGTCAGACACTGAATCTTC -3'). A 1.5kb product was amplified from the targeted allele using primer A with reverse primer C (5'- CAGTCCTCTTCACATCCATG -3'), within the β-geo cassette. After enzymatic amplification for 35 cycles (45 seconds at 94 °C, 45 seconds at 55 °C, and 1 minute at 72 °C), the PCR products were size-fractionated on a 2% agarose gel in 1x Tris borate-EDTA buffer.

**Primers used for genotyping (a,b&c)** PCR genotyping of gene trap *Tanc1* mice using a common forward primer, a, and reverse primers b and c to amplify the wt and mutant alleles respectively.

#### 1.3 Breeding

Birth of *Tanc1* mutant mice followed non-Mendelian ratios with no offspring being homozygous knockouts. Genotypes of 3-week-old pups from *Tanc1*+/- intercrosses identified 11 wt, 20 *Tanc1*+/- and 0 *Tanc1*-/- progeny (X2 p= 0.0055). Male and female *Tanc1*+/- mice showed no gross abnormalities. *Tanc1* mice were maintained by backcrossing onto the C57BL/6J background.

### *Tnik* (TNiK)

#### 1 Mouse generation

##### 1.1 Mutation

Generation of *Tnik* mutant mice described in PMID:23035106, Coba *et al* (2012)

##### 1.2 Genotyping

*Tnik* genotyping described in PMID:23035106, Coba *et al* (2012)

##### 1.3 Breeding

Birth of *Tnik*$^{-/-}$ mice followed Mendelian ratios with 21% of offspring being homozygous knockouts. Genotypes of 3-week-old pups from *Tnik*$^{+/-}$ intercrosses identified 46 wt, 113 *Tnik*$^{+/-}$ and 43 *Tnik*$^{-/-}$ progeny (Chi-squared p= 0.23). Male and female *Tnik*$^{-/-}$ mice developed normally to adulthood, exhibited normal body size and no gross abnormalities. *Tnik* mice were maintained by backcrossing onto the 129S5/SvEvBrd background; heterozygous males and females were fertile and used to set up intercrosses to generate homozygous and wildtype mice to study.

### *Tjp1* (ZO-1)

#### 1 Mouse generation

##### 1.1 Mutation

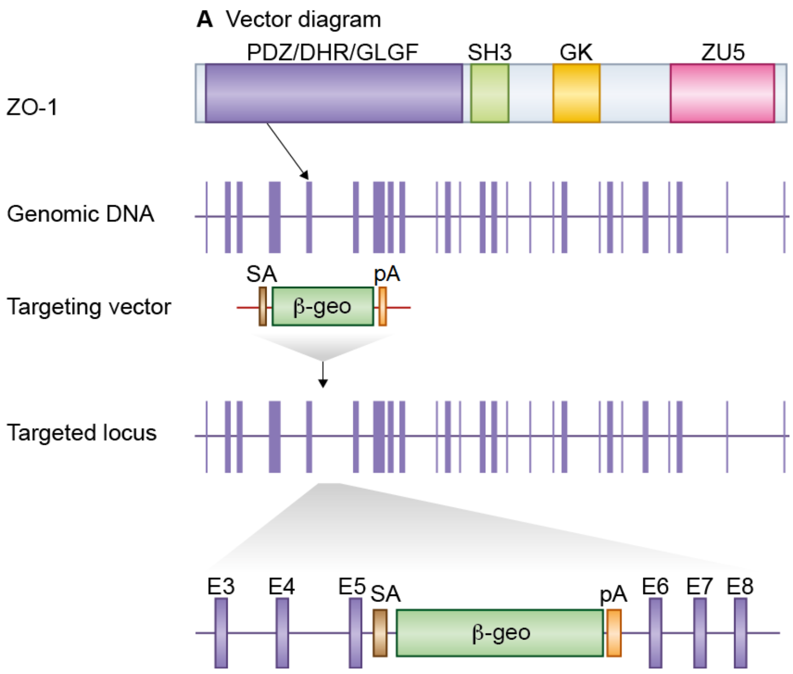


**Figure 1:** *Details of the mutation*

A mouse embryonic stem (ES) cell line (XH852, strain 129/Ola) with an insertional mutation in *Tjp1* was obtained from BayGenomics (baygenomics.ucsf.edu/). The insertional mutation in XH852, by the gene-trapping vector, pGT1lxf, that was designed to create an in-frame fusion between the 5' exons of the trapped gene and a reporter, ß-geo (a fusion of ß-galactosidase and neomycin phosphotransferase II), occurred in intron 4-5. Thus, the gene-trapped locus is predicted to yield a fusion transcript containing exons 1-4 of *Tjp1* and ß-geo. The ES cells were injected into C57BL/6 blastocysts to create chimeric mice, which were bred with 129S5 mice to generate heterozygous*Tjp1* mutant mice. Location of *Tjp1* gene trap. *Tjp1* is a 28 exon gene encoding the ZO-1 protein which contains SH3, GK, ZU-5 and three PDZ domains (top). The *Tjp1* gene trap is located within exon 5.

#### 1.2 Genotyping

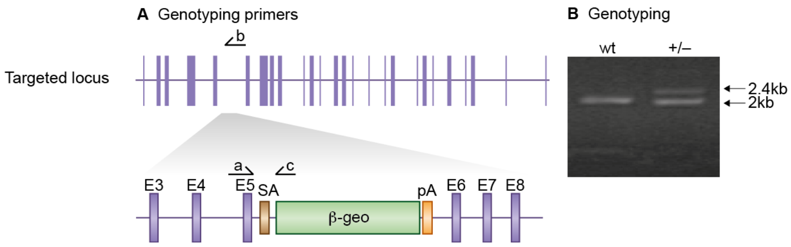


**Figure 2:** *Genotyping details*

Genomic DNA was isolated from ES cells or mouse tissues by Wizard SV 96 Genomic DNA purification system (Promega Cat A2371). Genotyping PCR consisted of a 2kb product amplified from the wild-type (wt) allele using a forward primer A (5'- GAAAGTTCAGATCCCTGTAA -3') upstream of the cassette and a reverse primer B (5'- CTGGTCTACAGAACTAGTTC -3') in the wt sequence deleted by targeted mutation. A 2.4kb product was amplified from the targeted allele using primer A with reverse primer C (5'- GGTTACGTTGGT-GTAGATGG -3'), within the ß-geo cassette. After enzymatic amplification for 35 cycles (30 seconds at 94 degC, 1 minute at 55 degC, and 3 minutes at 72 degC), the PCR products were size-fractionated on a 0.6%agarose gel in 1x Tris borate-EDTA buffer. Primers used for genotyping (a,b, c). PCR genotyping of gene trap ZO-1 mice using a common forward primer, a, and reverse primers b and c to amplify the wt and mutant alleles respectively.

#### 1.3 Breeding

No *Tjp1*$^{-/-}$ mice were produced from *Tjp1*$^{+/-}$ intercrosses. Male and female *Tjp1*$^{+/-}$ mice developed normally to adulthood, were fertile, exhibited normal body size and no gross abnormalities. Genotypes of 3-week-old pups from *Tjp1*$^{+/-}$ intercrosses identified 26 wt and 49 *Tjp1*$^{+/-}$ progeny (Chi-squared p= $<$0.001). Backcrosses onto the 129S5/SvEvBrd background were used to maintain the colony and to generate heterozygous and wildtype mice to study.
