## Supplementary Figures and Tables for "A combinatorial postsynaptic molecular mechanism converts patterns of nerve impulses into the behavioral repertoire"

**Contents:**

Supplementary Figure 1. Genes to Cognition (G2C) Program.	2

Supplementary Table 1. Comparison of G2C phenotypes with previous reports	3

Supplementary Table 2. Human disease gene annotations	3

References for Supplementary Table 2	4

### Supplementary Figure 1. Genes to Cognition (G2C) Program.

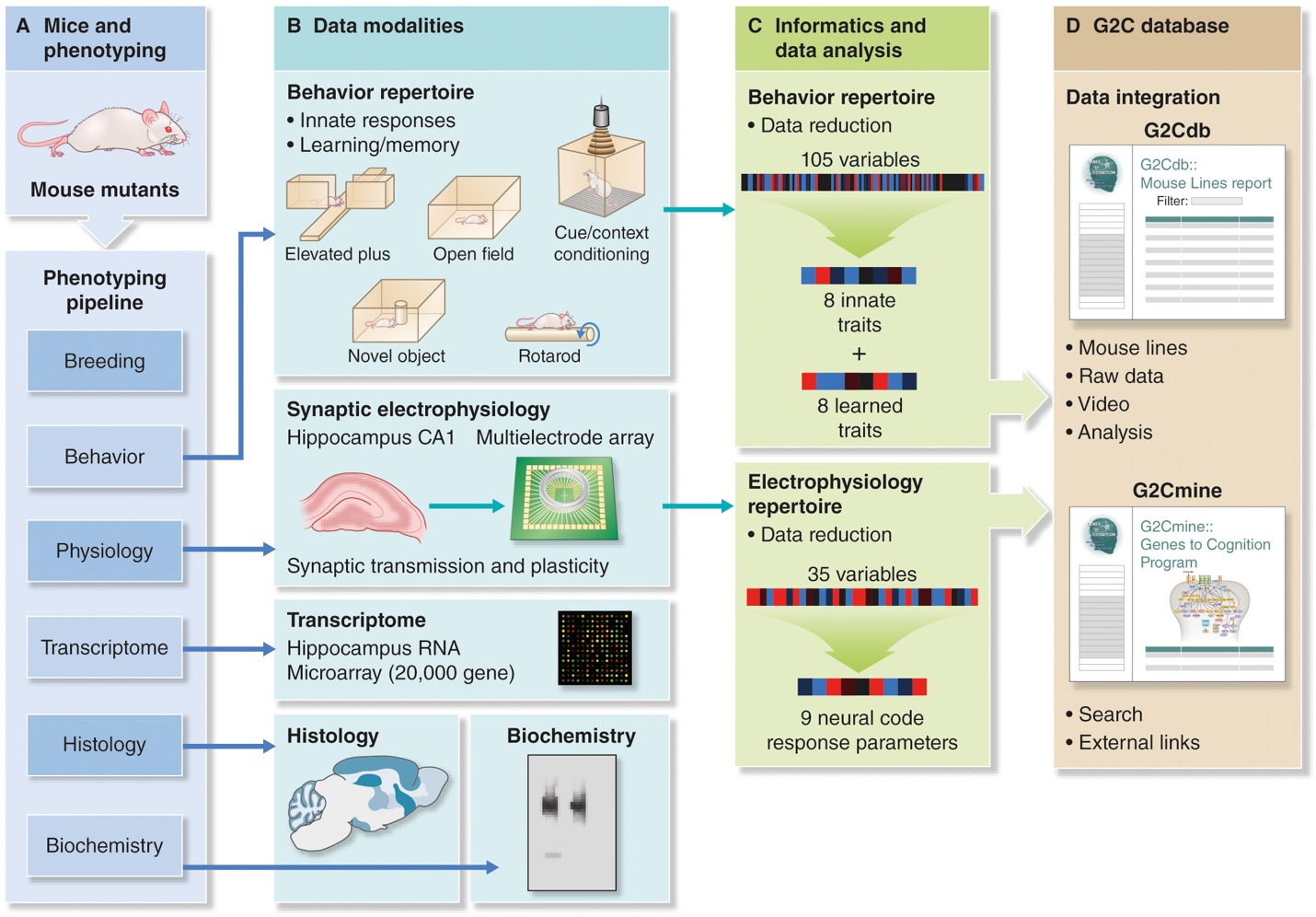

The Genes to Cognition program workflow of mutant mouse generation and standardized data acquisition, bioinformatics analysis and data integration.

### Supplementary Table 1. Comparison of G2C phenotypes with previous reports

Table shows comparison of electrophysiological phenotypes measured with the MEA methodology compared to previously published studies using other recording methods.

### Supplementary Table 2. Human disease gene annotations

Disease gene annotations for the genes/mouse lines studied. References for intellectual disability (ID), autism spectrum disorder (ASD), schizophrenia (SCZ) shown and Online Mendelian Inheritance of Man (OMIM) reference numbers for neural diseases shown.

| **Gene** | **Line** | **ID** | **ASD** | **SCZ** | **OMIM neural disease** |
| --- | --- | --- | --- | --- | --- |
| *Akt2* | Akt2 | 1 |  |  |  |
| *Arhgap32* | GRIT |  |  | 2 |  |
| *Baiap2* | IRSp53 |  | 3 | 4 |  |
| *Cadm1* | SynCam |  | 5 |  |  |
| *Camk2a* | αCaMKII | 6 |  |  |  |
| *Cnksr2* | MAGUIN-1 | 7–9 |  |  |  |
| *Cyfip1* | CYFIP1 | 10,11 | 12 | 13,14 |  |
| *Dlg2* | PSD-93 |  | 15 | 16–19 |  |
| *Dlg3* | SAP102 | 20–23 |  |  | 300189 |
| *Dlg4* | PSD-95-GK | 24 | 25 |  |  |
| *Dlgap1* | GKAP |  |  | 26 |  |
| *Dlgap2* | GKAP2 |  | 27 | 28 |  |
| *Git1* | Git1 | 29 |  | 30 |  |
| *Glul* | Glul | 31 |  |  | 138290 |
| *Gnb1* | Gnb1 | 32 |  |  | 139380 |
| *Gria1* | GluR1 | 33 |  | 34,35 |  |
| *Grin2a* | NR2A-dC | 33,36 |  |  | 138253 |
| *Grin2b* | NR2B-dC | 33,36 |  |  | 138252 |
| *Hras* | H-Ras | 37–40 |  |  | 190020 |
| *Kras* | K-Ras | 41–45 |  |  | 190070 |
| *Nf1* | NF1 | 46–48 |  |  | 162200 |
| *Nsf* | NSF |  |  | 13 |  |
| *Shank2* | SHANK2 | 49 | 27 |  | 603290 |
| *Syngap1* | SynGAP | 33,50–57 | 27,58,59 | 60 | 603384 |
| *Tnik* | TNiK | 61 |  | 62–64 | 610005 |
| *Tspyl2* | CINAP | 65,66 |  |  |  |

### References for Supplementary Table 2

1	Garg, N. *et al.* MORFAN Syndrome: An Infantile Hypoinsulinemic Hypoketotic Hypoglycemia Due to an AKT2 Mutation. *J Pediatr* **167**, 489-491, doi:10.1016/j.jpeds.2015.04.069 (2015).

2	Ohi, K. *et al.* The p250GAP gene is associated with risk for schizophrenia and schizotypal personality traits. *PLoS One* **7**, e35696, doi:10.1371/journal.pone.0035696 (2012).

3	Toma, L. *et al.* Association study of six candidate genes asymmetrically expressed in the two cerebral hemispheres suggests the involvement of BAIAP2 in autism. *J Psychiatr Res* **45**, 280–282, doi: 10.1016/j.jpsychires.2010.09.001 (2011).

4	Purcell, S. M. *et al.* A polygenic burden of rare disruptive mutations in schizophrenia. *Nature* **506**, 185-190, doi:10.1038/nature12975 (2014).

5	Zhiling, Y. *et al.* Mutations in the gene encoding CADM1 are associated with autism spectrum disorder. *Biochem Biophys Res Commun* **377**, 926-929, doi:10.1016/j.bbrc.2008.10.107 (2008).

6	Vincent, M. *et al.* Large deletions encompassing the TCOF1 and CAMK2A genes are responsible for Treacher Collins syndrome with intellectual disability. *Eur J Hum Genet* **22**, 52-56, doi:10.1038/ejhg.2013.98 (2014).

7	Houge, G., Rasmussen, I. H. & Hovland, R. Loss-of-Function CNKSR2 Mutation Is a Likely Cause of Non-Syndromic X-Linked Intellectual Disability. *Mol Syndromol* **2**, 60-63, doi:000335159 (2012).

8	Hu, H. *et al.* X-exome sequencing of 405 unresolved families identifies seven novel intellectual disability genes. *Mol Psychiatry* **21**, 133-148, doi:10.1038/mp.2014.193 (2016).

9	Vaags, A. K. *et al.* Absent CNKSR2 causes seizures and intellectual, attention, and language deficits. *Ann Neurol* **76**, 758-764, doi:10.1002/ana.24274 (2014).

10	Vanlerberghe, C. *et al.* 15q11.2 microdeletion (BP1-BP2) and developmental delay, behavior issues, epilepsy and congenital heart disease: a series of 52 patients. *Eur J Med Genet* **58**, 140-147, doi:10.1016/j.ejmg.2015.01.002 (2015).

11	von der Lippe, C., Rustad, C., Heimdal, K. & Rodningen, O. K. 15q11.2 microdeletion - seven new patients with delayed development and/or behavioral problems. *Eur J Med Genet* **54**, 357-360, doi:10.1016/j.ejmg.2010.12.008 (2011).

12	van der Zwaag, B. *et al.* A co-segregating microduplication of chromosome 15q11.2 pinpoints two risk genes for autism spectrum disorder. *Am J Med Genet B Neuropsychiatr Genet* **153B**, 960-966, doi:10.1002/ajmg.b.31055 (2010).

13	Tam, G. W. *et al.* Confirmed rare copy number variants implicate novel genes in schizophrenia. *Biochem Soc Trans* **38**, 445-451, doi:10.1042/BST0380445 (2010).

14	Zhao, Q. *et al.* Rare CNVs and tag SNPs at 15q11.2 are associated with schizophrenia in the Han Chinese population. *Schizophr Bull* **39**, 712-719, doi:10.1093/schbul/sbr197 (2013).

15	Egger, G. *et al.* Identification of risk genes for autism spectrum disorder through copy number variation analysis in Austrian families. *Neurogenetics* **15**, 117-127, doi:10.1007/s10048-014-0394-0 (2014).

16	International Schizophrenia, C. Rare chromosomal deletions and duplications increase risk of schizophrenia. *Nature* **455**, 237-241, doi:10.1038/nature07239 (2008).

17	Kirov, G. *et al.* De novo CNV analysis implicates specific abnormalities of postsynaptic signalling complexes in the pathogenesis of schizophrenia. *Mol Psychiatry* **17**, 142-153, doi:10.1038/mp.2011.154 (2012).

18	Walsh, T. *et al.* Rare structural variants disrupt multiple genes in neurodevelopmental pathways in schizophrenia. *Science* **320**, 539-543, doi:10.1126/science.1155174 (2008).

19	Xu, B. *et al.* Strong association of de novo copy number mutations with sporadic schizophrenia. *Nat Genet* **40**, 880-885, doi:10.1038/ng.162 (2008).

20	Philips, A. K. *et al.* X-exome sequencing in Finnish families with intellectual disability--four novel mutations and two novel syndromic phenotypes. *Orphanet J Rare Dis* **9**, 49, doi:10.1186/1750-1172-9-49 (2014).

21	Tarpey, P. *et al.* Mutations in the DLG3 gene cause nonsyndromic X-linked mental retardation. *Am J Hum Genet* **75**, 318-324, doi:10.1086/422703 (2004).

22	Tzschach, A. *et al.* Next-generation sequencing in X-linked intellectual disability. *Eur J Hum Genet* **23**, 1513-1518, doi:10.1038/ejhg.2015.5 (2015).

23	Zanni, G. *et al.* A novel mutation in the DLG3 gene encoding the synapse-associated protein 102 (SAP102) causes non-syndromic mental retardation. *Neurogenetics* **11**, 251-255, doi:10.1007/s10048-009-0224-y (2010).

24	Lelieveld, S. H. *et al.* Meta-analysis of 2,104 trios provides support for 10 new genes for intellectual disability. *Nat Neurosci*, doi:10.1038/nn.4352 (2016).

25	Xing, J. *et al.* Resequencing and Association Analysis of Six PSD-95-Related Genes as Possible Susceptibility Genes for Schizophrenia and Autism Spectrum Disorders. *Sci Rep* **6**, 27491, doi:10.1038/srep27491 (2016).

26	Li, J. M. *et al.* Genetic analysis of the DLGAP1 gene as a candidate gene for schizophrenia. *Psychiatry Res* **205**, 13-17, doi:10.1016/j.psychres.2012.08.014 (2013).

27	Pinto, D. *et al.* Functional impact of global rare copy number variation in autism spectrum disorders. *Nature* **466**, 368-372, doi:10.1038/nature09146 (2010).

28	Greenwood, T. A. *et al.* Genetic assessment of additional endophenotypes from the Consortium on the Genetics of Schizophrenia Family Study. *Schizophr Res* **170**, 30-40, doi:10.1016/j.schres.2015.11.008 (2016).

29	Uddin, M. *et al.* Indexing Effects of Copy Number Variation on Genes Involved in Developmental Delay. *Sci Rep* **6**, 28663, doi:10.1038/srep28663 (2016).

30	Fromer, M. *et al.* De novo mutations in schizophrenia implicate synaptic networks. *Nature* **506**, 179-184, doi:10.1038/nature12929 (2014).

31	Marchese, M., Valvo, G., Moro, F., Sicca, F. & Santorelli, F. M. Targeted Gene Resequencing (Astrochip) to Explore the Tripartite Synapse in Autism-Epilepsy Phenotype with Macrocephaly. *Neuromolecular Med* **18**, 69-80, doi:10.1007/s12017-015-8378-2 (2016).

32	Petrovski, S. *et al.* Germline De Novo Mutations in GNB1 Cause Severe Neurodevelopmental Disability, Hypotonia, and Seizures. *Am J Hum Genet* **98**, 1001-1010, doi:10.1016/j.ajhg.2016.03.011 (2016).

33	de Ligt, J. *et al.* Diagnostic exome sequencing in persons with severe intellectual disability. *N Engl J Med* **367**, 1921-1929, doi:10.1056/NEJMoa1206524 (2012).

34	Kang, W. S. *et al.* Genetic variants of GRIA1 are associated with susceptibility to schizophrenia in Korean population. *Mol Biol Rep* **39**, 10697-10703, doi:10.1007/s11033-012-1960-x (2012).

35	Magri, C. *et al.* Glutamate AMPA receptor subunit 1 gene (GRIA1) and DSM-IV-TR schizophrenia: a pilot case-control association study in an Italian sample. *Am J Med Genet B Neuropsychiatr Genet* **141B**, 287-293, doi:10.1002/ajmg.b.30294 (2006).

36	Endele, S. *et al.* Mutations in GRIN2A and GRIN2B encoding regulatory subunits of NMDA receptors cause variable neurodevelopmental phenotypes. *Nat Genet* **42**, 1021-1026, doi:10.1038/ng.677 (2010).

37	Kerr, B. *et al.* Genotype-phenotype correlation in Costello syndrome: HRAS mutation analysis in 43 cases. *J Med Genet* **43**, 401-405, doi:10.1136/jmg.2005.040352 (2006).

38	Lo, I. F. *et al.* Severe neonatal manifestations of Costello syndrome. *J Med Genet* **45**, 167-171, doi:10.1136/jmg.2007.054411 (2008).

39	Sol-Church, K., Stabley, D. L., Nicholson, L., Gonzalez, I. L. & Gripp, K. W. Paternal bias in parental origin of HRAS mutations in Costello syndrome. *Hum Mutat* **27**, 736-741, doi:10.1002/humu.20381 (2006).

40	Zampino, G. *et al.* Diversity, parental germline origin, and phenotypic spectrum of de novo HRAS missense changes in Costello syndrome. *Hum Mutat* **28**, 265-272, doi:10.1002/humu.20431 (2007).

41	Bertola, D. R. *et al.* Further evidence of genetic heterogeneity in Costello syndrome: involvement of the KRAS gene. *J Hum Genet* **52**, 521-526, doi:10.1007/s10038-007-0146-1 (2007).

42	Carta, C. *et al.* Germline missense mutations affecting KRAS Isoform B are associated with a severe Noonan syndrome phenotype. *Am J Hum Genet* **79**, 129-135, doi:10.1086/504394 (2006).

43	Niihori, T. *et al.* Germline KRAS and BRAF mutations in cardio-facio-cutaneous syndrome. *Nat Genet* **38**, 294-296, doi:10.1038/ng1749 (2006).

44	Schubbert, S. *et al.* Germline KRAS mutations cause Noonan syndrome. *Nat Genet* **38**, 331-336, doi:10.1038/ng1748 (2006).

45	Zenker, M. *et al.* Expansion of the genotypic and phenotypic spectrum in patients with KRAS germline mutations. *J Med Genet* **44**, 131-135, doi:10.1136/jmg.2006.046300 (2007).

46	Kayes, L. M. *et al.* Deletions spanning the neurofibromatosis 1 gene: identification and phenotype of five patients. *Am J Hum Genet* **54**, 424-436 (1994).

47	Kayes, L. M., Riccardi, V. M., Burke, W., Bennett, R. L. & Stephens, K. Large de novo DNA deletion in a patient with sporadic neurofibromatosis 1, mental retardation, and dysmorphism. *J Med Genet* **29**, 686-690 (1992).

48	Wallace, M. R. *et al.* A de novo Alu insertion results in neurofibromatosis type 1. *Nature* **353**, 864-866, doi:10.1038/353864a0 (1991).

49	Berkel, S. *et al.* Mutations in the SHANK2 synaptic scaffolding gene in autism spectrum disorder and mental retardation. *Nat Genet* **42**, 489-491, doi:10.1038/ng.589 (2010).

50	Berryer, M. H. *et al.* Mutations in SYNGAP1 cause intellectual disability, autism, and a specific form of epilepsy by inducing haploinsufficiency. *Hum Mutat* **34**, 385-394, doi:10.1002/humu.22248 (2013).

51	Hamdan, F. F. *et al.* Excess of de novo deleterious mutations in genes associated with glutamatergic systems in nonsyndromic intellectual disability. *Am J Hum Genet* **88**, 306-316, doi:10.1016/j.ajhg.2011.02.001 (2011).

52	Hamdan, F. F. *et al.* Mutations in SYNGAP1 in autosomal nonsyndromic mental retardation. *N Engl J Med* **360**, 599-605, doi:10.1056/NEJMoa0805392 (2009).

53	Klitten, L. L. *et al.* A balanced translocation disrupts SYNGAP1 in a patient with intellectual disability, speech impairment, and epilepsy with myoclonic absences (EMA). *Epilepsia* **52**, e190-193, doi:10.1111/j.1528-1167.2011.03304.x (2011).

54	Krepischi, A. C. *et al.* A novel de novo microdeletion spanning the SYNGAP1 gene on the short arm of chromosome 6 associated with mental retardation. *Am J Med Genet A* **152A**, 2376-2378, doi:10.1002/ajmg.a.33554 (2010).

55	Mignot, C. *et al.* Genetic and neurodevelopmental spectrum of SYNGAP1-associated intellectual disability and epilepsy. *J Med Genet* **53**, 511-522, doi:10.1136/jmedgenet-2015-103451 (2016).

56	Parker, M. J. *et al.* De novo, heterozygous, loss-of-function mutations in SYNGAP1 cause a syndromic form of intellectual disability. *Am J Med Genet A* **167A**, 2231-2237, doi:10.1002/ajmg.a.37189 (2015).

57	Rauch, A. *et al.* Range of genetic mutations associated with severe non-syndromic sporadic intellectual disability: an exome sequencing study. *Lancet* **380**, 1674-1682, doi:10.1016/S0140-6736(12)61480-9 (2012).

58	Iossifov, I. *et al.* The contribution of de novo coding mutations to autism spectrum disorder. *Nature* **515**, 216-221, doi:10.1038/nature13908 (2014).

59	O'Roak, B. J. *et al.* Recurrent de novo mutations implicate novel genes underlying simplex autism risk. *Nat Commun* **5**, 5595, doi:10.1038/ncomms6595 (2014).

60	Xu, B. *et al.* De novo gene mutations highlight patterns of genetic and neural complexity in schizophrenia. *Nat Genet* **44**, 1365-1369, doi:10.1038/ng.2446 (2012).

61	Anazi, S. *et al.* A null mutation in TNIK defines a novel locus for intellectual disability. *Hum Genet* **135**, 773-778, doi:10.1007/s00439-016-1671-9 (2016).

62	Ayalew, M. *et al.* Convergent functional genomics of schizophrenia: from comprehensive understanding to genetic risk prediction. *Mol Psychiatry* **17**, 887-905, doi:10.1038/mp.2012.37 (2012).

63	Potkin, S. G. *et al.* A genome-wide association study of schizophrenia using brain activation as a quantitative phenotype. *Schizophr Bull* **35**, 96-108, doi:10.1093/schbul/sbn155 (2009).

64	Shi, J. *et al.* Common variants on chromosome 6p22.1 are associated with schizophrenia. *Nature* **460**, 753-757, doi:10.1038/nature08192 (2009).

65	Moey, C. *et al.* Xp11.2 microduplications including IQSEC2, TSPYL2 and KDM5C genes in patients with neurodevelopmental disorders. *Eur J Hum Genet* **24**, 373-380, doi:10.1038/ejhg.2015.123 (2016).

66	Vasli, N. *et al.* Identification of a homozygous missense mutation in LRP2 and a hemizygous missense mutation in TSPYL2 in a family with mild intellectual disability. *Psychiatr Genet* **26**, 66-73, doi:10.1097/YPG.0000000000000114 (2016).
